# New estimates indicate that males are not larger than females in most mammals

**DOI:** 10.1101/2023.02.23.529628

**Authors:** Kaia J. Tombak, Severine B. S. W. Hex, Daniel I. Rubenstein

**Affiliations:** Department of Anthropology, Hunter College of the City University of New York, New York, NY, USA; Department of Ecology and Evolutionary Biology, Princeton University, Princeton, NJ, USA

## Abstract

Sexual size dimorphism (SSD) has motivated a large body of research on mammalian mating strategies and sexual selection. Despite some contrary evidence, the narrative that larger males are the norm in mammals – upheld since Darwin’s *Descent of Man* – still dominates today, supported by meta-analyses that use crude measures of dimorphism and taxonomically-biased data. With newly-available datasets and primary sources reporting sex-segregated means and variances in adult body mass, we estimated statistically-determined rates of SSD in mammals, sampling taxa by their species richness at the family level. Our analyses of >400 species indicate that although males tend to be larger than females *when* dimorphism occurs, males are *not* larger in most mammals, and suggest a need to revisit other assumptions in sexual selection research.

**One-Sentence Summary:** Taxonomically-balanced estimates of rates of sexual size dimorphism in mammals refute the ‘larger males’ narrative.

## MAIN TEXT

A long-standing narrative postulates that in mammals, males are typically larger than females. Darwin treated it as a matter of common knowledge (1), as have many subsequent evolutionary biologists studying sexual selection (*2–6*). The principal hypothesis predicting a prevalence of larger males in mammals is that the reproductive investment that females commit to their offspring (via gestation, lactation, and often parental care) results in a skewed operational sex ratio, leading to greater mate competition and selection for competitive ability among males (*1, 7, 8*). This pattern should be especially strong under polygyny, presumed to be the most common mating system in mammals (*7, 9*). In the 1970’s, Ralls contributed the first review on rates of sexual size dimorphism (SSD) in mammals and found weak support for this hypothesis. She concluded that most mammals are ‘not extremely dimorphic’, that species with little sexual size dimorphism were extremely numerous in the most species-rich mammalian orders (*10*), and that larger females were surprisingly common in mammals (*11*). Nonetheless, her findings have been overpowered by the continuation of the ‘larger males’ narrative (*2–4, 6, 12–14*), despite some additional evidence supporting her conclusions (*15, 16*).

Issues with data availability and taxonomic biases have hindered efforts to accurately estimate the rate of SSD in mammals. Meta-analyses have so far been limited to using mean adult body mass values for each sex, a measure that is widely available in the literature but typically reported without measures of variance that would allow for a statistical assessment of dimorphism. To designate species as dimorphic or monomorphic, researchers have therefore used either arbitrary cut-offs (including a 5% (*17*), 10% (*13*), and up to a 20% difference in mean body mass (*18*)) or ratios below or above 1 between mean male and female body masses (*13, 19*). Ralls herself used mean body mass ratios to assess the modal degree of dimorphism in each mammalian order (*10*). Using different criteria influences the conclusions. For instance, Lindenfors et al. (*13*) concluded that mammals generally had male-biased size dimorphism because the average body mass ratio was >1 across their sample, but their other analyses using a 10% body mass difference cut-off indicated that less than half of mammalian species had male-biased SSD (*13*). Neither criterion is based on sufficient information to assign a species as dimorphic or monomorphic: body mass difference thresholds are both arbitrary and inconsistent, and a body mass ratio >1 can indicate either more frequent dimorphism *or* more extreme dimorphism in one sex than the other. In addition, research on SSD in mammals has tended to focus on a few taxa, namely artiodactyls, carnivores (especially pinnipeds), and primates (*7, 19–23*): clades with high rates of male-biased SSD (*24*). However, most mammals, by far, are rodents and bats (*25*), which are often under-represented in studies of SSD. The phylogenetic signal for SSD is strong (*6*), calling for updated estimates with more balanced taxonomic representation.

Fortunately, some recently-published large datasets report mean body mass as well as measures of variance for each sex across mammalian taxa. We combined these datasets with data from primary sources to revisit Ralls’s original question, estimating the rates of sexual size dimorphism in wild, non-provisioned mammalian populations using statistical determinations of dimorphism for each taxon and sampling each mammalian order and family according to their species richness.

## Results

Our final dataset included 405 species with a minimum sample size of 9 for each sex, the minimum sample size that mitigates for the inflation of confidence intervals with low sample size (Fig. S1). We achieved at least 5% representation for each mammalian order except for Eulipotyphla (3.8%), and Perissodactyla (for which no study met our criteria, see Materials and Methods). We also achieved at least 5% representation for 58 out of the 76 mammalian families that contain at least 10 species (Fig. S2, Table S1, Data S1). Our estimates, based on the frequency with which the 95% confidence interval for the between-sex difference in mean body mass straddles zero, and weighted by species richness in each family, indicated that 38.7% of mammalian species are sexually monomorphic in body mass, while 44.5% of species are male-biased dimorphic and 16.8% are female-biased dimorphic (Fig. 1).

**Figure 1:**
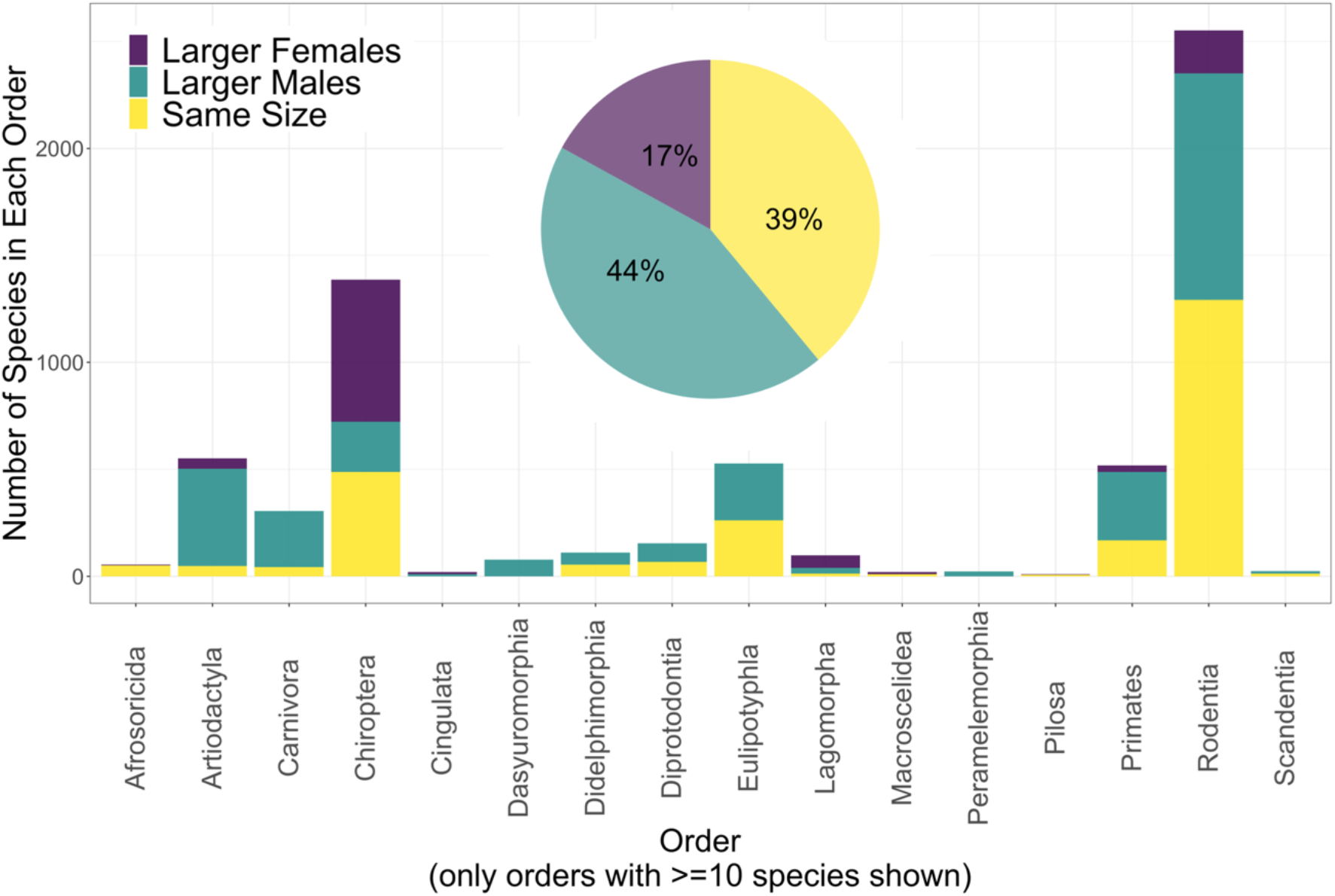
Estimated rates of SSD in mammals as a whole (pie chart), and in each mammalian order (bar chart), weighted by species richness in each family

Male-biased dimorphism was somewhat more extreme on average than female-biased dimorphism (mean male/female body mass ratio in male-biased dimorphic species = 1.28, N=169; mean female/male body mass ratio in female-biased dimorphic species = 1.13, N=66). This confirms that average male/female mass ratios >1 are inappropriate indicators of the frequency of dimorphism. The most dimorphic species was the northern elephant seal, where males had a mean mass 3.2 times that of females (*26*). The most extreme female-biased dimorphism was found in the peninsular tube-nosed bat (*Murina peninsularis*), in which mean female mass was 1.4 times that of males (*27*). However, most dimorphisms were not extreme (Fig. 2), as Ralls concluded almost 50 years ago (*10*). When we reran the analyses on rates of SSD on body length instead of body mass in the subset of our data with body length measurements (see Methods and Materials), our estimates shifted towards more monomorphism and female-biased dimorphism (48.0% monomorphic, 29.7% male-biased dimorphic, 22.3% female-biased dimorphic; N=172).

**Figure 2:**
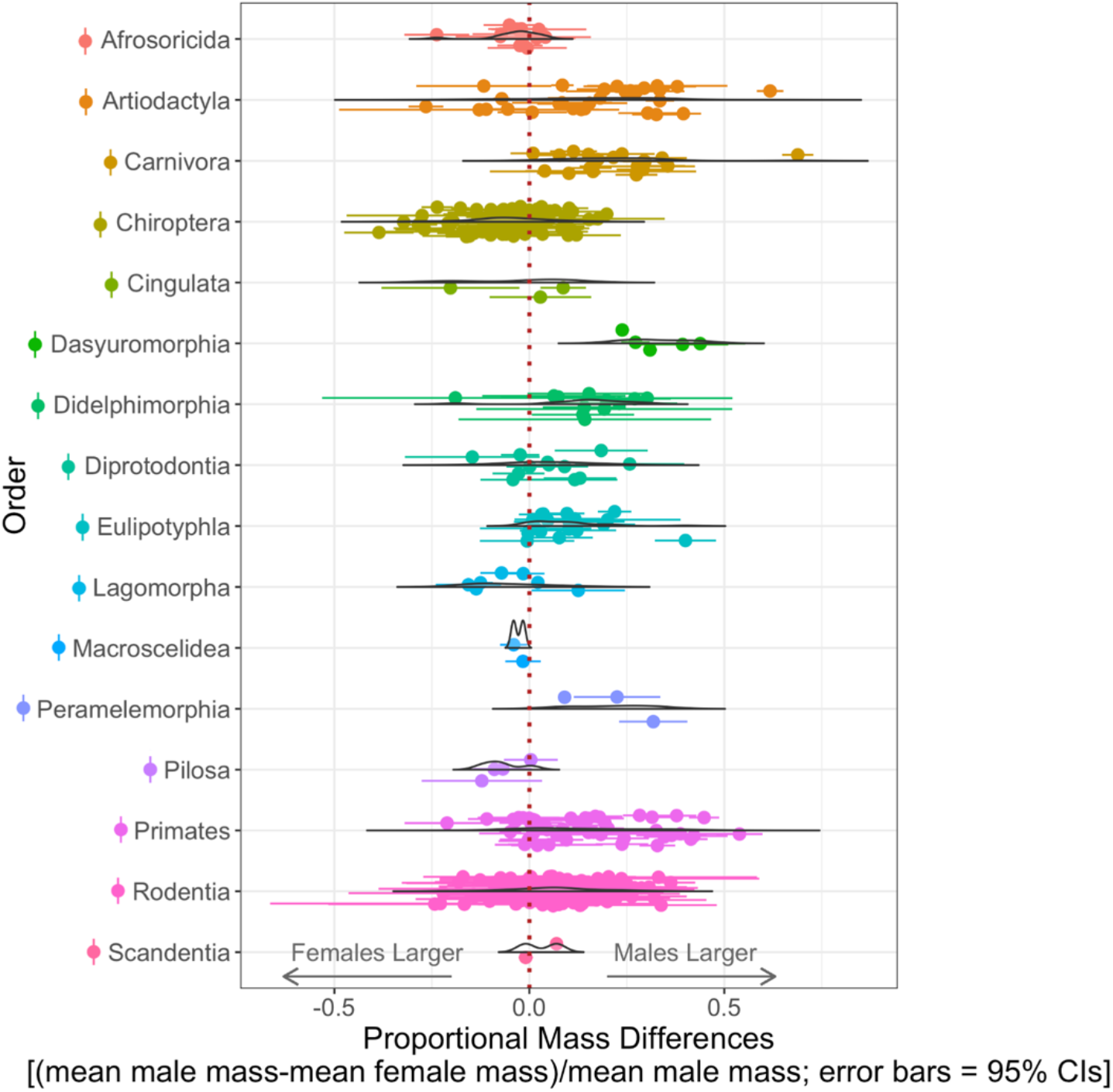
Distribution of mean mass differences, relative to male body mass, and 95% confidence intervals of the mass differences for all data collected that met our criteria for inclusion, plotted separately by order. Data density distributions are scaled such that the area under the curve is the same for each order. A proportional mass difference of −0.2 indicates that females are 20% larger than males, on average.

Overall, standard deviation in body mass was greater in males (mean SD=1920.5, median SD=12.9) than in females (mean SD=1171.5, median SD=11.1), so our results were unlikely to be seriously confounded by data that may have included pregnant females without our knowledge (paired Wilcoxon Rank Sum test V=52536, *p*<0.0001). Further, standard deviation in body mass was greater in males in male-dimorphic species (median male SD=62.7, median female SD=39.4, V=12494, *p*<0.0001), greater in females in female-dimorphic species (median male SD=2.6, median female SD=3.2, V=446, *p*<0.0001), and no different between the sexes in monomorphic species (median male SD=8.4, median female SD=9.3, V=6468, *p*=0.17).

Patterns of SSD differed markedly between orders (Figs. 1, 2, S2). About half of the species in Rodentia (the most species-rich order) were monomorphic, whereas about half of Chiroptera (the second-most species-rich order) had larger females (Table S1). Larger males were the norm for several of the less species-rich orders, while several others were evenly divided between larger males and monomorphism, and larger females were the norm for Lagomorpha (Figs. 2, S2). Notably, the orders that had the most prevalent male-biased dimorphism included Artiodactyla, Carnivora, and Primates: the orders that dominate the SSD literature for mammals (*7, 19–23*). Differences in rates of SSD at the family level were also evident, indicating that weighting our estimates based on species richness in each family was important. For example, the famously larger females in Lagomorpha were so only in Leporidae (rabbits and hares), while Ochotonidae (pikas) have more male-biased dimorphism. In Primates, larger males are the norm overall, but strepsirrhine primates are mostly monomorphic, as are about half of Cebidae (Fig. S2, Table S1).

## Discussion

Our results did not support the ‘larger males’ narrative. While species with larger males were the largest single category, we found that males are not larger than females in most mammalian species, and that sexual monomorphism was almost as frequent as larger males (and *more* frequent if body length is used instead of body mass). Our results accord with Ralls’s original reviews (*10, 11*), with smaller-scale meta-analyses on species-rich mammalian taxa (*15, 28*), and with a very large-scale meta-analysis that found the same rate of male-biased SSD as we did using a 10% body mass difference cut-off (*13*). Yet the latter study fell back on male/female body mass ratios to conclude that mammals generally have larger males and Ralls’s review – which was the only review of rates of SSD in mammals for many years – has been miscited several times as having supported the ‘larger males’ narrative (*6, 13, 14*).

Why has this narrative persisted so stubbornly? It may be ascribed to the long-time focus of SSD research on species with conspicuous dimorphisms, as suggested by Bondrup-Nielsen and Ims (*29*) and by Dewsbury *et al*. (*30*). However, given the well-established variation in dimorphism across mammalian taxa, it is surprising that so many would accept generalizations based on a few, relatively species-poor taxa. The narrative may also be traced to a long-standing research focus on male mating strategies in the study of evolution (*31, 32*), particularly in mammals (*33*). Darwin himself focused almost entirely on how sexual selection operated on males in the form of mate competition when discussing mammals (*1*). Competitive males and choosy females are a recurring theme in animal behavior research, based on the argument that females invest more energy in gametes and are therefore the less reproductively available sex: the controversial ‘Darwin-Bateman-Trivers’ paradigm (*1, 2, 34*). The dominance of this paradigm and general androcentrism in sexual selection research are likely to have influenced which narratives are readily accepted and amplified and which are overlooked or subjected to heavier scrutiny (*32, 35*).

Importantly, ours should not be the last word on rates of sexual size dimorphism in mammals. We prioritized data quality over quantity, and our conclusions are based on data covering only 5% of mammalian species. However, our results align well with those based on a 10% body mass difference across >1,300 mammalian species (*13*). In addition, some minor taxonomic biases persist in our dataset. Mammals of very high body mass are difficult to weigh and, when such data are reported, often have low sample sizes. However, the most underrepresented taxa by species richness were still small-bodied, speciose clades (certain families in Eulipotyphla, Chiroptera, and Rodentia; Fig. S2). In general, given the suppressed reporting of non-significant results in science (and we frequently saw statements of a lack of sexual dimorphism unaccompanied by descriptive statistics in primary sources) (*36*), our estimated rates of monomorphism are probably underestimates. Even in many statistically dimorphic species, the dimorphism was minimal, and what constitutes biologically meaningful dimorphism will vary according to the physiology of a species and its ecological niche. Finally, body mass varies by body condition and is not an ideal measure of size for many taxa (*15, 37, 38*). Our preliminary results showing a prevalence of sexual monomorphism in body *length* in mammals reinforce the idea that it may be time to retire the ‘larger males’ narrative.

More nuanced language and thought around dimorphism would also improve sexual selection research. Many species do not fit neatly into a dimorphic or monomorphic category (*39*). There can be great intraspecific variation in both body size and dimorphism in mammals, whether at the individual level (e.g., extreme seasonal body mass fluctuations in both males and females in prairie dogs, *Cynomys* spp., with the result that males are much larger than females in the beginning of the breeding season but statistically the same size by the end of it (*40*)) or population level (e.g., a latitudinal cline in the short-nosed fruit bat, *Cynopterus sphinx*, ranging from male-biased to female-biased dimorphism (*9*)). High variance in body mass within a sex also presents complications for categorizing species as simply dimorphic or monomorphic; for example, the fossa (*Cryptoprocta ferox*) has two male morphs, one of which is the same size as adult females, and the other of which is larger (*42*). Our results indicate that the sex that is larger on average tends to be the sex with greater variance in body mass, challenging the notion that absolute size is strongly linked to sex in most mammals (i.e., a greater average body mass may be driven by a relatively small subset of individuals of the larger sex in many cases). This variety also underlines the potential for multiple alternative reproductive strategies in either sex in mammalian species (*39*).

Given the building evidence for a greater prevalence of sexual size monomorphism than is commonly recognized, the theoretical basis for the evolution of SSD in mammals deserves some reframing. If mate competition is expected to be stronger in the more reproductively available sex (*43*), our findings suggest either that the relative availability of males is not so much greater than females in most mammals as is traditionally assumed, or that greater availability does not typically result in greater size. For instance, selection for agility in mate competition instead of size is one hypothesis for the prevalence of smaller males in bats (*44*) (however, *greater* size in female bats has also been argued to enhance agility in flight while carrying offspring (*14*)). Meanwhile, how sexual selection, and mate competition in particular, operate among female mammals is generally understudied (*32*). In animal behavior research it is commonly assumed that all available females will choose the strongest, most dominant male as a mate, or else be coerced by him into copulation – but many populations have shown great variation in female mate preferences, as well as aggressive competition among females for mates (*45, 46*), some instances of which have been passed off as capricious behavior rather than adaptive and strategic (*32*). The biased predictions in sexual selection research have thus potentially generated blind spots in terms of what the patterns in nature really are on multiple counts. SSD research needs to refocus on how multiple selection pressures act on body size in both sexes, including how sex-specific pressures (e.g., fecundity selection and size selection for carrying offspring load in females) (*1, 14, 47*) balance with natural selection acting on body size in both sexes (e.g., heat stress, agility and detectability with respect to predators) (*48*). In addition, we call for rigorous investigation of other long-standing and related narratives, such as the often-repeated claim that polygyny is by far the most prevalent mating system in mammals, with some estimating rates of 90% of species or more (*3, 9*). As old assumptions are revisited with larger datasets and greater scrutiny, we see great potential in new breakthroughs in sexual selection theory.

## Materials and Methods

### Data Collection

We searched Google Scholar for datasets on sex-segregated body mass data for mammalian species that reported measures of variance, standard deviation, or 95% confidence intervals as well as sample size for each sex within a population. Extremely large datasets on mean body mass values are available but do not report measures of variance for each sex (e.g., the PanTHERIA database, the Handbook of Mammalian Body Masses, the Handbook of Mammals of the World, AnimalTraits) (*49–51*). Many others do not report body mass data in a sex-segregated format (e.g., the Malagasy Animal trait Data Archive, EltonTraits1.0) (*52, 53*). However, we found several published datasets that included data that met our criteria (*16, 29, 38, 54–58*). We then searched Google Scholar for primary sources to top up sample sizes for underrepresented orders and families in our dataset, using Burgin et al.’s (*25*) estimations of species richness in each mammalian order. For primates, we found additional data on underrepresented taxa from the *All the World’s Primates* online database (*59*). When we found more than one study to report body mass values for the same species, we used the data from the study with greater sample sizes in analyses. Subspecies were treated as different populations of the same species and these data were not combined even if reported in the same study. We set a goal of achieving 5% representation of the extant species in every mammalian order with at least 10 species (the lowest number for which 5% can be rounded up to 1 species), and further aimed to sample 5% of all families within each of these orders to minimize taxonomic biases.

We excluded any measures from sexually immature, pregnant, or captive-bred animals when these were distinguished (or when it was noted that these were mixed in with the data). One exception was that we accepted weight measurements from Fischer’s pygmy fruit bats (*Haplonycteris fischeri*) for which weights were reported for females with very early-stage fetuses (1.5-5mm in length). When data were presented for non-breeding seasons separately, data from breeding seasons were excluded. Only data from wild animals were used, and free-ranging animals that were provisioned with food were excluded. Estimates based on museum specimens were generally not used as we wished to make within-population comparisons of body mass between the sexes, and domestic species were also excluded, although we did use data from semi-domesticated, free-ranging reindeer (*Rangifer tarandus*). Only direct body mass measurements were included, with the exception of Baird’s beaked whales (*Berardius bairdii*), and male northern elephant seals (*Mirounga angustirostris*), for which we accepted body mass estimated based on body lengths, girths, and/or ultrasound measurements due to the logistical difficulty of weighing such heavy animals in the wild (*26*). Where means, sample sizes, and measures of variance were reported separately for different seasons or sites for one population, these were combined using Baker and Nissim’s (*60*) equation to calculate the standard error for the combined sample.

We focused our data search on body mass because these data are the most available measure of body size in the literature. However, we did collect body length measurements wherever these were reported for the same population for which we obtained body mass data, to serve as verification that our conclusions would not be very different if body length were used. For these, we used head and body length (excluding the tail, when possible) for most taxa, but forearm length was used for bats, hindfoot length for lagomorphs, and head length for dasyuromorphs and peramelemorphs, according to the conventions and data availability for these taxa.

### Statistical Analyses

We calculated the 95% confidence interval of the difference in mean mass between males and females for each species (*61*) and labeled each species as either monomorphic (95% CI of mass difference straddles zero) or dimorphic (95% CI does not straddle zero). Our initial dataset included a total of 609 mammalian species, but this included some populations with a sample size of only 2 for each sex. Lower sample sizes decrease confidence in the mean masses, broadening the 95% confidence intervals and the likelihood of being assigned as monomorphic. We therefore plotted, for each sex in turn, the difference in mean mass between the sexes, divided by mean body mass, against sample size and found the elbow of the exponential decay function in this relationship (N=9.6 for males, N=9.2 for females) using the *findCutoff* function in the ‘KneeArrower’ package (*62*) (Fig. S1). A minimum sample size of 9 for each sex was thus used as a criterion for inclusion in the analyses, producing a final count of 405 species included.

Some orders and families within orders were highly overrepresented relative to their species richness in our final dataset. To ensure that these did not contribute disproportionately to our estimates of rates of dimorphism, we randomly sampled rows in our dataset within each overrepresented family such that they were sampled exactly according to their species richness. In other words, each family was assigned the number of rows corresponding to 5% of the number of species in the family (rounded to the closest integer), and only this many rows were randomly sampled from the family to produce estimated rates of dimorphism for each order. This random sampling within mammalian families was performed with replacement 1000 times and the frequencies of female-biased dimorphism, male-biased dimorphism, and monomorphism were tabulated each time to enable the calculation of species-richness-adjusted average frequencies of dimorphism for each order and for mammals as a whole. The order Pilosa contains only ten species but is divided into four families, which were represented in the dataset roughly in proportion to their species richness and from which only one row for the order was sampled randomly for each iteration.

To verify that our estimates of the rates of dimorphism do not considerably change if we use body length data, we similarly sampled the subset of species from the final dataset for which the source reported body length measurements (N=172) by species richness. However, since fewer body length data were available than body mass data, we set the goal to 1% representation and sampled by species richness only at the level of the order, rather than for each family.

## Supporting information

Data Sources

## Acknowledgements

We thank Dr. Sara Weinstein and all authors of van Schaik et al. (2015) for generously sharing their data on their study taxa through personal communications. This research was supported by the National Science Foundation (IBN-9874523, CNS-025214, and IOB-9874523) and the Simons Foundation (grant #638529).

## Data Availability Statement

All data used for analyses will be published in a public repository upon acceptance of this manuscript. For the time-being, we have appended a summary table of the results (Table S1) and a datasheet listing the species included and sources used (Data S1).

## References

1. C. Darwin, The descent of man and selection in relation to sex (John Murray, London, United Kingdom, 1871).

2. R. L. Trivers, “Parental investment and sexual selection” in Sexual Selection and the Descent of Man, B. G. Campbell, Ed. (Harvard University Press, 1st editio., 1972), pp. 136–179.

3. E. Mori, G. Mazza, S. Lovari, “Sexual Dimorphism” in Encyclopedia of Animal Cognition and Behavior, J. Vonk, T. K. Shackelford, Eds. (Springer International Publishing, 2017).

4. E. Dinerstein, “Size and sexual dimorphism in greater one-horned rhinoceros” in The Return of the Unicorns: The Natural History and Conservation of the Greater One-Horned Rhinoceros, E. Dinerstein, G. Schaller, Eds. (Columbia University Press, 2003), pp. 61–80.

5. H. K. Kurki, Pelvic dimorphism in relation to body size and body size dimorphism in humans. J. Hum. Evol. 61, 631–643 (2011).

6. E. Abouheif, D. J. Fairbairn, A comparative analysis of allometry for sexual size dimorphism: Assessing Rensch’s rule. Am. Nat. 149, 540–562 (1997).

7. R. D. Alexander, J. L. Hoogland, R. D. Howard, K. M. Noonan, P. W. Sherman, “Sexual dimorphisms and breeding systems in pinnipeds, ungulates, primates, and humans” in Evolutionary Biology and Human Social Behavior, N. A. Chagnon, W. Irons, Eds. (Duxbury Press, North Scituate, Massachusetts, 1979), pp. 402–435.

8. A. V. Hedrick, E. J. Temeles, The evolution of sexual dimorphism in animals: hypotheses and tests. Trends Ecol. Evol. 4, 136–138 (1989).

9. T. H. Clutton-Brock, Mammalian mating systems. Proc. R. Soc. B Biol. Sci. 236, 339–372 (1989).

10. K. Ralls, Sexual Dimorphism in Mammals: Avian Models and Unanswered Questions. Am. Nat. 111, 917–938 (1977).

11. K. Ralls, Mammals in which females are larger than males. Q. Rev. Biol. 5, 245–276 (1976).

12. R. D. Alexander, G. Borgia, “On the origin and basis of the male-female phenomenon” in Sexual selection and reproductive competition in insects, M. F. Blum, N. Blum, Eds. (Academic Press, New York, 1979), pp. 417–440.

13. P. Lindenfors, J. L. Gittleman, K. E. Jones, “Sexual size dimorphism in mammals” in Sex, Size and Gender Roles: Evolutionary Studies of Sexual Size Dimorphism (University of Oxford Press, 2007), pp. 16–26.

14. P. J. Greenwood, P. Wheeler, “The evolution of sexual size dimorphism in birds and mammals: a “hot blooded” hypothesis” in Evolution: Essays in hoour of John Maynard Smith (1985), pp. 287–299.

15. D. Lu, C. Q. Zhou, W. B. Liao, Sexual size dimorphism lacking in small mammals. North. West. J. Zool. 10, 53–59 (2014).

16. P. M. Kappeler, C. L. Nunn, A. Q. Vining, S. M. Goodman, Evolutionary dynamics of sexual size dimorphism in non-volant mammals following their independent colonization of Madagascar. Sci. Rep. 9, 1–14 (2019).

17. J. Carranza, F. J. Pérez-Barbería, Sexual selection and senescence: Male size-dimorphic ungulates evolved relatively smaller molars than females. Am. Nat. 170, 370–380 (2007).

18. K. E. Ruckstuhl, P. Neuhaus, Sexual segregation in ungulates: A comparative test of three hypotheses. Biol. Rev. Camb. Philos. Soc. 77, 77–96 (2002).

19. E. Ranta, A. Laurila, J. Elmberg, Reinventing the Wheel: Analysis of Sexual Dimorphism in Body Size. Oikos. 70, 313–321 (1994).

20. A. Mysterud, The relationship between ecological segregation and sexual body size dimorphism in large herbivores. Oecologia. 124, 40–54 (2000).

21. F. J. Pérez-Barbería, I. J. Gordon, M. Pagel, The origins of sexual dimorphism in body size in ungulates. Evolution (N. Y). 56, 1276–1285 (2002).

22. J. M. Plavcan, Sexual dimorphism in primate evolution. Yearb. Phys. Anthropol. 44, 25–53 (2001).

23. T. H. Clutton-Brock, P. H. Harvey, Primate ecology and social organization. J. Zool. 183, 1–39 (1977).

24. F. W. Weckerly, Sexual-size dimorphism: Influence of mass and mating systems in the most dimorphic mammals. J. Mammal. 79, 33–52 (1998).

25. C. J. Burgin, J. P. Colella, P. L. Kahn, N. S. Upham, How many species of mammals are there? J. Mammal. 99, 1–14 (2018).

26. S. S. Kienle, A. S. Friedlaender, D. E. Crocker, R. S. Mehta, D. P. Costa, Trade-offs between foraging reward and mortality risk drive sex-specific foraging strategies in sexually dimorphic northern elephant seals. R. Soc. Open Sci. 9 (2022), doi:10.1098/rsos.210522.

27. P. Soisook, S. Karapan, C. Satasook, V. D. Thong, F. A. A. Khan, I. Maryanto, G. Csorba, N. Furey, B. Aul, P. J. J. Bates, A Review of the Murina cyclotis complex (Chiroptera: Vespertilionidae) with descriptions of a new species and subspecies. Acta Chiropterologica. 15, 271–292 (2013).

28. P. M. Kappeler, A framework for studying social complexity. Behav. Ecol. Sociobiol. 73 (2019), doi:10.1007/s00265-018-2601-8.

29. S. Bondrup-Nielsen, R. A. Ims, Reversed sexual size dimorphism in microtines: Are females larger than males or are males smaller than females? Evol. Ecol. 4, 261–272 (1990).

30. D. A. Dewsbury, D. J. Baumgardner, R. L. E. Evans, G. W. Daniel, Sexual Dimorphism for Body Mass in 13 Taxa of Muroid Rodents under Laboratory Conditions. J. Mammal. 61, 146–149 (1980).

31. T. Clutton-Brock, Sexual selection in males and females. Science (80-.). 318, 1882–1885 (2007).

32. M. Ah-King, The history of sexual selection research provides insights as to why females are still understudied. Nat. Commun. 13, 6976 (2022).

33. T. H. Clutton-Brock, K. Mcauliffe, Female choice in mammals. Q. Rev. Biol. 84, 3–27 (2009).

34. A. J. Bateman, Intra-sexual selection in Drosophila. Heredity (Edinb). 2, 349–368 (1948).

35. Z. Tang-Martinez, Rethinking Bateman’s Principles: Challenging Persistent Myths of Sexually Reluctant Females and Promiscuous Males. J. Sex Res. 53, 532–559 (2016).

36. E. W. van Zwet, E. A. Cator, The significance filter, the winner’s curse and the need to shrink. Stat. Neerl. 75, 437–452 (2021).

37. M. T. O’Mara, K. Bauer, D. Blank, J. W. Baldwin, K. Dina, N. Dechmann, Common Noctule Bats Are Sexually Dimorphic in Migratory Behaviour and Body Size but Not Wing Shape. PLoS One. 11, e0167027 (2016).

38. A. I. Schulte-Hostedde, “Sexual size dimorphism in rodents” in Rodent Societies: An Ecological and Evolutionary Perspective, J. O. Wolff, P. W. Sherman, Eds. (University of Chicago Press, Chicago, 2007), pp. 115–128.

39. J. E. Mank, Sex-specific morphs: the genetics and evolution of intra-sexual variation. Nat. Rev. Genet. 24, 44–52 (2023).

40. J. L. Hoogland, Sexual dimorphism of prairie dogs. J. Mammal. 84, 1254–1266 (2003).

41. J. F. Storz, J. Balasingh, H. R. Bhat, T. P. Nathan, P. D. Swami Doss, A. A. Prakash, T. H. Kunz, Clinal variation in body size and sexual dimorphism in an Indian fruit bat, Cynopterus sphinx (Chiroptera: Pteropodidae). Biol. J. Linn. Soc. 72, 17–31 (2001).

42. M. L. Lührs, M. Dammhahn, P. Kappeler, Strength in numbers: Males in a carnivore grow bigger when they associate and hunt cooperatively. Behav. Ecol. 24, 21–28 (2013).

43. T. H. Clutton-Brock, A. C. J. Vincent, Sexual selection and the potential reproductive rates of males and females. Nature. 351, 58–60 (1991).

44. M. Reiss, “Sexual dimorphism in body size” in The Allometry of Growth and Reproduction (Cambridge University Press, Cambridge, 1989).

45. F. Widemo, S. A. Sæther, Beauty is in the eye of the beholder: Causes and consequences of variation in mating preferences. Trends Ecol. Evol. 14, 26–31 (1999).

46. M. D. Jennions, M. Petrie, Variation in mate choice and mating preferences: A review of causes and consequences. Biol. Rev. 72, 283–327 (1997).

47. R. Shine, Ecological causes for the evolution of sexual dimorphism: a review of the evidence. Q. Rev. Biol. 64, 419–461 (1989).

48. W. U. Blanckenhorn, The evolution of body size: What keeps organisms small? Q. Rev. Biol. 75, 385–407 (2000).

49. M. Silva, J. A. Downing, CRC Handbook of Mammalian Body Masses (CRC Press, Boca Raton, FL, 1995).

50. K. E. Jones, J. Bielby, M. Cardillo, S. A. Fritz, J. O’Dell, C. D. L. Orme, K. Safi, W. Sechrest, E. H. Boakes, C. Carbone, C. Connolly, M. J. Cutts, J. K. Foster, R. Grenyer, M. Habib, C. A. Plaster, PanTHERIA: a species-level database of life history, ecology, and geography of extant and recently extinct mammals. Ecology. 90 (2009).

51. M. E. Herberstein, D. J. Mclean, E. Lowe, J. O. Wol, K. Khan, K. Smith, A. P. Allen, M. Bulbert, B. A. Buzatto, M. D. B. Eldridge, D. Falster, L. F. Winzer, S. C. Gri, J. S. Madin, A. Narendra, M. Westoby, M. J. Whiting, I. J. Wright, A. J. R. Carthey, AnimalTraits - a curated animal trait database for body mass, metabolic rate and brain size, 1–11 (2022).

52. O. H. Razafindratsima, Y. Yacoby, D. S. Park, MADA: Malagasy Animal trait Data Archive. Ecology. 99, 990 (2018).

53. H. Wilman, W. Jetz, J. Belmaker, J. Simpson, J. Hagberry, C. De, C. Anderson, M. Rivadeneira, EltonTraits 1.0: Species-level foraging attributes of the world’s birds and mammals. Ecology. 95, 2027 (2014).

54. D. Ocampo, K. G. Borja-Acosta, J. Lozano-Flórez, S. Cifuentes-Acevedo, E. Arbeláez-Cortés, N. J. Bayly, Á. Caguazango, B. Coral-Jaramillo, D. Cueva, F. Forero, J. P. Gómez, C. Gómez, M. A. Loaiza-Muñoz, G. A. Londoño, S. Losada-Prado, S. Pérez-Peña, H. E. Ramírez-Chaves, M. E. Rodríguez-Posada, J. Sanabria-Mejía, M. Sánchez-Martínez, V. H. Serrano-Cardozo, M. S. Sierra-Buitrago, J. Soto-Patiño, O. Acevedo-Charry, Body mass data set for 1,317 bird and 270 mammal species from Colombia. Ecology. 102 (2021), doi:10.1002/ecy.3273.

55. F. Gonçalves, R. S. Bovendorp, G. Beca, C. Bello, R. Costa-Pereira, R. L. Muylaert, R. R. Rodarte, N. Villar, R. Souza, M. E. Graipel, J. J. Cherem, D. Faria, J. Baumgarten, M. R. Alvarez, E. M. Vieira, N. Cáceres, R. Pardini, Y. L. R. Leite, L. P. Costa, M. A. R. Mello, E. Fischer, F. C. Passos, L. H. Varzinczak, J. A. Prevedello, A. P. Cruz-Neto, F. Carvalho, A. R. Percequillo, A. Paviolo, A. Nava, J. M. B. Duarte, N. U. de la Sancha, E. Bernard, R. G. Morato, J. F. Ribeiro, R. G. Becker, G. Paise, P. S. Tomasi, F. Vélez-Garcia, G. L. Melo, J. Sponchiado, F. Cerezer, M. A. S. Barros, A. Q. S. de Souza, C. C. dos Santos, G. A. F. Giné, P. Kerches-Rogeri, M. M. Weber, G. Ambar, L. V. Cabrera-Martinez, A. Eriksson, M. Silveira, C. F. Santos, L. Alves, E. Barbier, G. C. Rezende, G. S. T. Garbino, É. O. Rios, A. Silva, A. T. A. Nascimento, R. S. de Carvalho, A. Feijó, J. Arrabal, I. Agostini, D. Lamattina, S. Costa, E. Vanderhoeven, F. R. de Melo, P. de Oliveira Laroque, L. Jerusalinsky, M. M. Valença-Montenegro, A. B. Martins, G. Ludwig, R. B. de Azevedo, A. Anzóategui, M. X. da Silva, M. Figuerêdo Duarte Moraes, A. Vogliotti, A. Gatti, T. Püttker, C. S. Barros, T. K. Martins, A. Keuroghlian, D. P. Eaton, C. L. Neves, M. S. Nardi, C. Braga, P. R. Gonçalves, A. C. Srbek-Araujo, P. Mendes, J. A. de Oliveira, F. A. M. Soares, P. A. Rocha, P. Crawshaw, M. C. Ribeiro, M. Galetti, Atlantic Mammal Traits: a data set of morphological traits of mammals in the Atlantic Forest of South America. Ecology. 99, 498 (2018).

56. R. J. Smith, W. L. Jungers, Body mass in comparative primatology. J. Hum. Evol. 32, 523–559 (1997).

57. M. Colyn, Données pondérales sur les primates Cercopithecidae d’Afrique Centrale (Bassin du Zaire/Congo). Mammalia. 58, 483–487 (1994).

58. P. M. Kappeler, Patterns of sexual dimorphism in body weight among prosimian primates. Folia Primatol. 57, 132–146 (1991).

59. N. Rowe, M. Myers, “All the World’s Primates” (Charlestown RI, 2022), (available at http://www.alltheworldsprimates.org).

60. R. W. R. Baker, J. A. Nissim, Expressions for combining standard errors of two groups and for sequential standard error. Nature. 198, 1020 (1963).

61. Finnstats, Calculate Confidence Intervals in R. R-bloggers (2021).

62. A. Tseng, KneeArrower: Finds Cutoff Points on Knee Curves (2020).

63. J. E. Lovich, J. W. Gibbons, Review of techniques for quantifying sexual size dimorphism. Growth, Dev. Aging. 56, 269–281 (1992).

