## Supplemental Figures and Tables for "New estimates indicate that males are not larger than females in most mammals"

**Fig. S1.**

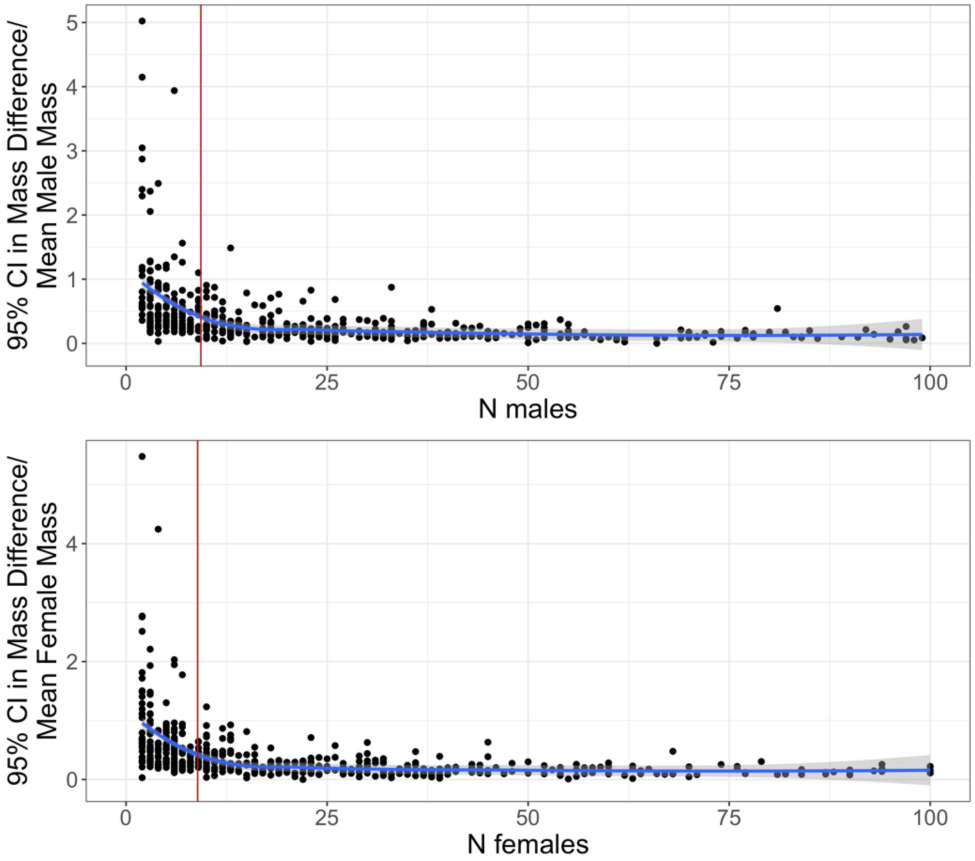


**Sample size and absolute confidence interval, relative to body mass, for each sex (males in the top panel, females in the bottom panel), with the elbow of the curve marked in red. The elbow was used to determine the minimum sample size for a datapoint to be included in analyses (N=9 males and 9 females).**

**Fig. S2.**

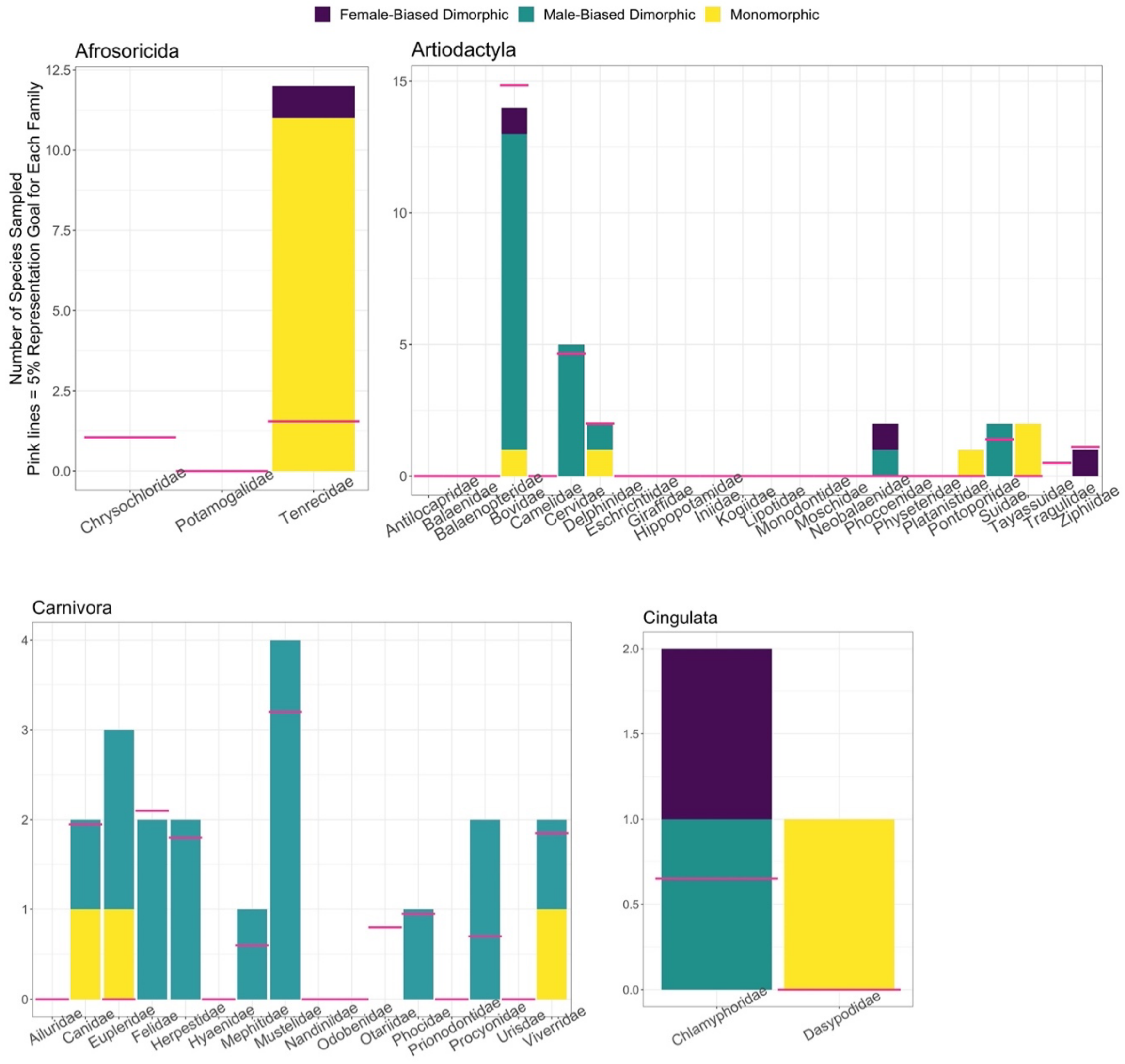
Female-Biased Dimorphic
Male-Biased Dimorphic
Monomorphic
Afrosoricida
Number of Species Sampled
Pink lines = 5% Representation Goal for Each Family
0.0
2.5
5.0
7.5
10.0
12.5
Chrysochloridae
Potamogalidae
Tenrecidae
Artiodactyla
0
5
10
15
Antilocapridae
Balaenidae
Balaenopteridae
Bovidae
Camelidae
Cervidae
Delphinidae
Eschrichtiidae
Giraffidae
Hippopotamidae
Iniidae
Kogiidae
Lipotidae
Monodontidae
Moschidae
Neobalaenidae
Phocoenidae
Physeteridae
Platanistidae
Pontoporiidae
Suidae
Tayassuidae
Tragulidae
Ziphiidae
Carnivora
0
1
2
3
4
Ailuridae
Canidae
Eupleridae
Felidae
Herpestidae
Hyaenidae
Mephitidae
Mustelidae
Nandiniidae
Odobenidae
Otariidae
Phocidae
Prionodontidae
Procyonidae
Urisdae
Viverridae
Cingulata
0.0
0.5
1.0
1.5
2.0
Chlamyphoridae
Dasypodidae

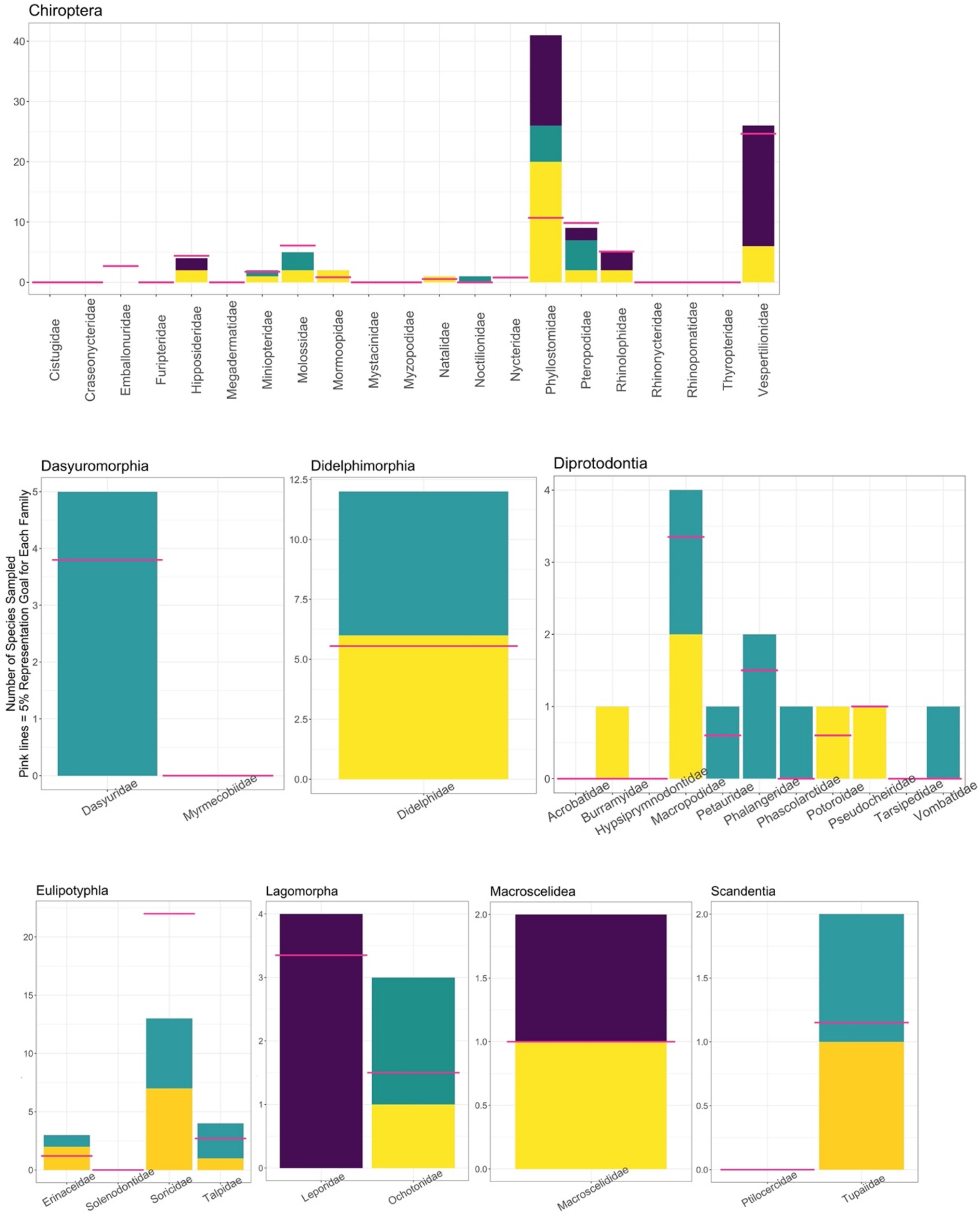
Chiroptera
0
10
20
30
40
Cistugidae
Craseonycteridae
Emballonuridae
Furipteridae
Hipposideridae
Megadermatidae
Miniopteridae
Molossidae
Mormoopidae
Mystacinidae
Myzopodidae
Natalidae
Noctilionidae
Nycteridae
Phyllostomidae
Pteropodidae
Rhinolophidae
Rhinonycteridae
Rhinopomatidae
Thyropteridae
Vespertilionidae
Number of Species Sampled
Pink lines = 5% Representation Goal for Each Family
Dasyuromorphia
Dasyuridae
Myrmecobiidae
Didelphimorphia
Didelphidae
Diprotodontia
Acrobatidae
Burramyidae
Hypsiprymnodontidae
Macropodidae
Petauridae
Phalangeridae
Phascolarctidae
Potoroidae
Pseudocheiridae
Tarsipedidae
Vombatidae
Eulipotyphla
Erinaceidae
Solenodontidae
Soricidae
Talpidae
Lagomorpha
Leporidae
Ochotonidae
Macroscelidea
Macroscelididae
Scandentia
Ptilocercidae
Tupaiidae

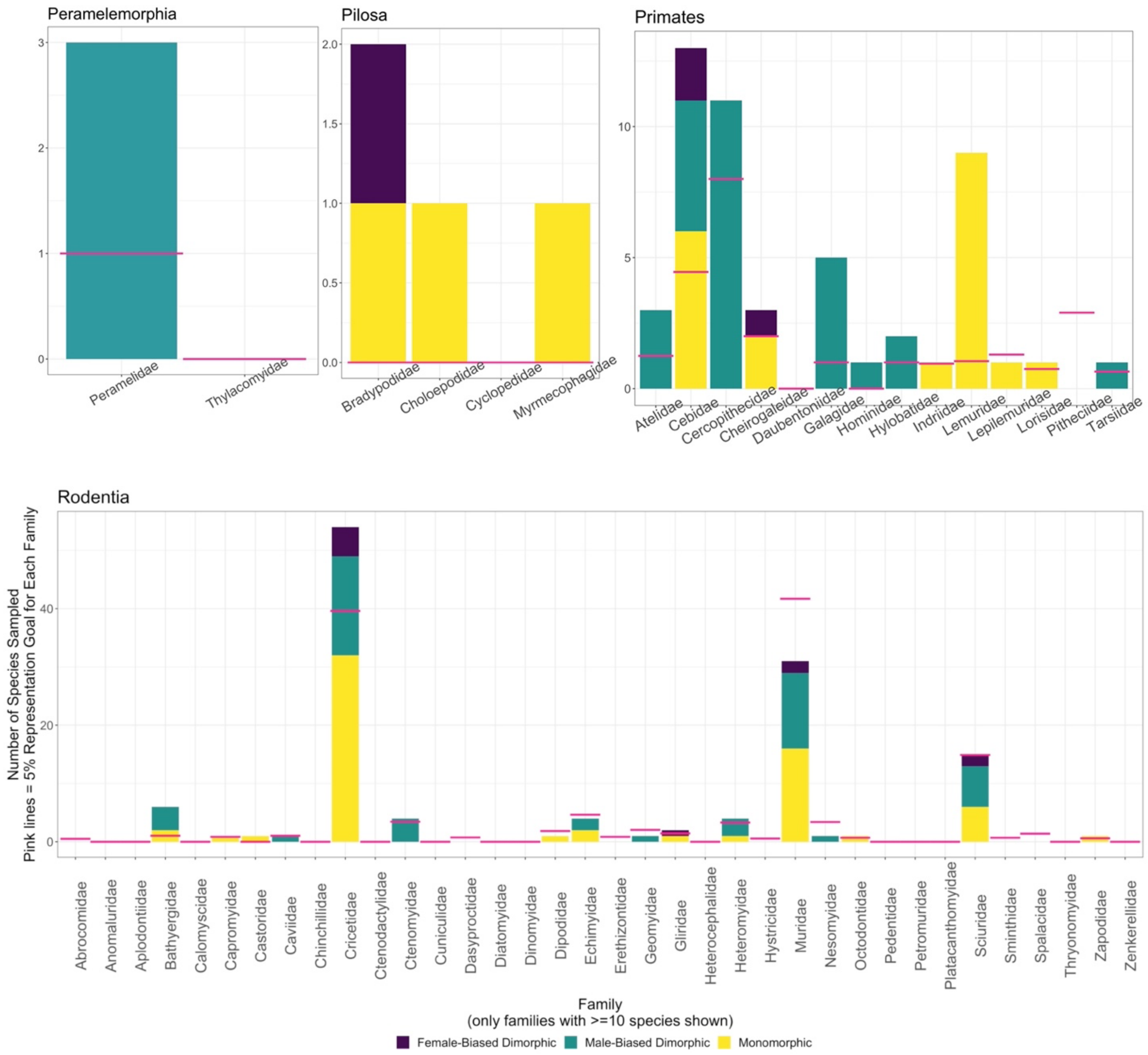


**Observed distribution of sexual size dimorphism across mammalian families in each order. Bars show the number of species in each category in our dataset, floating pink bars the 5% representation goal (= the maximum number of species included in each rate estimation).**

**Table S1.**

| ORDER<br>FAMILY | DESCRIPTION | # SPECIES SAMPLED/ # EXTANT SPECIES (% OF FAMILY) | EST. % OF ORDER & # IN FAMILY WITH LARGER FEMALES | EST. % OF ORDER & # IN FAMILY WITH MONOMORPHISM | EST. % OF ORDER & # IN FAMILY WITH LARGER MALES |
|---|---|---|---|---|---|
| **Rodentia** | **Rats, mice, & allies** | **128/2552 (5%)** | **7.9%** | **50.6%** | **41.5%** |
| Abrocomidae | Chinchilla rats | 0/10 (0%) | | | |
| Bathyergidae | African mole rats | 6/21 (28.6%) | 0 | 2 | 4 |
| Capromyidae | Hutias | 1/17 (5.9%) | 0 | 1 | 0 |
| Castoridae | Beavers | 1/2 (50%) | 0 | 1 | 0 |
| Caviidae | Cavies & maras | 1/21 (4.8%) | 0 | 0 | 1 |
| Cricetidae | Hamsters, voles, lemmings, muskrats, New World mice, & rats | 54/792 (6.8%) | 5 | 32 | 17 |
| Ctenomyidae | Tuco-tucos | 4/69 (5.8%) | 0 | 0 | 4 |
| Dasyproctidae | Agoutis & acouchis | 0/15 (0%) | | | |
| Dipodidae | Jerboas | 1/37 (2.7%) | 0 | 1 | 0 |
| Echimyidae | Shiny rats | 4/93 (4.3%) | 0 | 2 | 2 |
| Erethizontidae | New World porcupines | 0/17 (0%) | | | |
| Geomyidae | Gophers | 1/41 (2.4%) | 0 | 0 | 1 |
| Gliridae | Dormice | 2/29 (6.9%) | 1 | 1 | 0 |
| Heteromyidae | Kangaroo rats, kangaroo mice, pocket mice, & spiny pocket mice | 4/66 (6.1%) | 0 | 1 | 3 |
| Muridae | Mice, rats, & gerbils | 31/834 (3.7%) | 2 | 16 | 13 |
| Nesomyidae | Pouched rats, swamp mice, & allies | 1/68 (1.5%) | 0 | 0 | 1 |
| Octodontidae | Rock rats, coruros, degus, mountain degus, & viscacha rats | 1/14 (7.1%) | 0 | 1 | 0 |
| Sciuridae | Squirrels | 15/298 (5%) | 2 | 6 | 7 |
| Sminthidae | Birch mice | 0/14 (0%) | | | |
| Spalacidae | Blind mole-rats, bamboo rats, mole-rats, & zokors | 0/28 (0%) | | | |

| | | | | | |
|---|---|---|---|---|---|
| Zapodidae | Jumping mice | 1/12 (8.3%) | 0 | 1 | 0 |
| **Chiroptera** | **Bats** | **96/1386 (6.9%)** | **47.9%** | **35.2%** | **16.9%** |
| Emballonuridae | Sac-winged bats | 0/54 (0%) | | | |
| Hipposideridae | Old World leaf-nosed bats | 4/88 (4.5%) | 2 | 2 | 0 |
| Miniopteridae | Bent-winged & long-winged bats | 2/35 (5.7%) | 0 | 1 | 1 |
| Molossidae | Free-tailed bats | 5/122 (4.1%) | 0 | 2 | 3 |
| Mormoopidae | Mustached, ghost-faced, & naked-backed bats | 2/17 (11.8%) | 0 | 2 | 0 |
| Natalidae | Funnel-eared bats | 1/11 (9.1%) | 0 | 1 | 0 |
| Noctilionidae | Bulldog bats | 1/2 (50%) | 0 | 0 | 1 |
| Nycteridae | Slit-faced bats | 0/16 (0%) | | | |
| Phyllostomidae | New World leaf-nosed bats | 41/214 (19.2%) | 15 | 20 | 6 |
| Pteropodidae | Old World fruit bats, flying foxes | 9/197 (4.6%) | 2 | 2 | 5 |
| Rhinolophidae | Horseshoe bats | 5/102 (4.9%) | 3 | 2 | 0 |
| Vespertilionidae | Vesper bats | 26/493 (5.3%) | 20 | 6 | 0 |
| **Artiodactyla** | **Even-toed ungulates** | **29/551 (5.3%)** | **8.7%** | **8.7%** | **82.6%** |
| Bovidae | Buffalo, antelopes, goat-antelopes, & allies | 14/297 (4.7%) | 1 | 1 | 12 |
| Cervidae | True deer & allies | 5/93 (5.4%) | 0 | 0 | 5 |
| Delphinidae | Dolphins | 2/40 (5%) | 0 | 1 | 1 |
| Phocoenidae | Porpoises | 2/7 (28.6%) | 1 | 0 | 1 |
| Pontoporiidae | The La Plata dolphin | 1/1 (100%) | 0 | 1 | 0 |
| Suidae | Pigs | 2/28 (7.1%) | 0 | 0 | 2 |
| Tayassuidae | Peccaries | 2/5 (40%) | 0 | 2 | 0 |
| Tragulidae | Chevrotains | 0/10 (0%) | | | |
| Ziphiidae | Beaked whales | 1/22 (4.5%) | 1 | 0 | 0 |
| **Eulipotyphla** | **Hedgehogs, moles, shrews, & allies** | **20/527 (3.8%)** | **0%** | **49.7%** | **50.3%** |

| | | | | | |
|---|---|---|---|---|---|
| Erinaceidae | Hedgehogs & moonrats | 3/24 (12.5%) | 0 | 2 | 1 |
| Soricidae | Shrews | 13/440 (3%) | 0 | 7 | 6 |
| Talpidae | Moles | 4 /54 (7.4%) | 0 | 1 | 3 |
| **Primates** | **Lemurs, monkeys, apes, & allies** | **51/518 (9.8%)** | **6%** | **32.6%** | **61.4%** |
| Atelidae | Howler, spider, woolly, & woolly spider monkeys | 3/23 (13%) | 0 | 0 | 3 |
| Cebidae | Capuchins & squirrel monkeys, combined here with night monkeys (Aotidae) & marmosets, tamarins, & lion tamarins (Callitrichidae) | 13/89 (14.6%) | 2 | 6 | 5 |
| Cercophithecidae | Old World monkeys | 11/160 (6.9%) | 0 | 0 | 11 |
| Cheirogaleidae | Dwarf & mouse lemurs | 3/40 (7.5%) | 1 | 2 | 0 |
| Galagidae | Bush babies | 5/20 (25%) | 0 | 0 | 5 |
| Hominidae | Great apes | 1/7 (14.3%) | 0 | 0 | 1 |
| Hylobatidae | Gibbons | 2/20 (10%) | 0 | 0 | 2 |
| Indriidae | Wooly lemurs & sifakas | 1/19 (5.3%) | 0 | 1 | 0 |
| Lemuridae | True lemurs, wooly lemurs, & allies | 9/21 (4.3%) | 0 | 9 | 0 |
| Lepilemuridae | Sportive lemurs | 1/26 (3.8%) | 0 | 1 | 0 |
| Lorisidae | Lorises, pottos, & angwantibos | 1/15 (6.7%) | 0 | 1 | 0 |
| Pitheciidae | Titis, sakis, & uakaris | 0/58 (0%) | | | |
| Tarsiidae | Tarsiers | 1/13 (7.7%) | 0 | 0 | 1 |
| **Carnivora** | **Cat-like & dog-like carnivores** | **19/305 (6.2%)** | **0%** | **14.3%** | **85.7%** |
| Canidae | Dogs, foxes, & allies | 2/39 (5.1%) | 0 | 1 | 1 |
| Eupleridae | Malagasy carnivores | 3/8 (37.5%) | 0 | 1 | 2 |
| Felidae | Cats | 2/42 (4.8%) | 0 | 0 | 2 |
| Herpestidae | Mongooses | 2/36 (5.6%) | 0 | 0 | 2 |
| Mephitidae | Skunks | 1/12 (8.3%) | 0 | 0 | 1 |
| Mustelidae | Weasels, otters, badgers & allies | 4/64 (6.3%) | 0 | 0 | 4 |
| Otariidae | Eared seals | 0/16 (0%) | | | |
| Phocidae | Earless seals | 1/19 (5.3%) | 0 | 0 | 1 |

| | | | | | |
|---|---|---|---|---|---|
| Procyonidae | Racoons, ringtails, coatis & allies | 2/14 (14.3%) | 0 | 0 | 2 |
| Viverridae | Civets, genets, & allies | 2/37 (5.4%) | 0 | 1 | 1 |
| **Diprotodontia** | **Kangaroos, koalas, & allies** | **12/155 (7.7%)** | **0%** | **44%** | **56%** |
| Burramyidae | Pygmy possums | 1/5 (20%) | 0 | 1 | 0 |
| Macropodidae | Kangaroos, wallabies, quokkas & allies | 4/67 (6%) | 0 | 2 | 2 |
| Petauridae | Trioks & gliders | 1/12 (8.3%) | 0 | 0 | 1 |
| Phalangeridae | Cuscus & brushtail possums | 2/30 (6.7%) | 0 | 0 | 2 |
| Phascolarctidae | Koalas | 1/1 (100%) | 0 | 0 | 1 |
| Potoroidae | Bettongs, potoroos, & rat-kangaroos | 1/12 (8.3%) | 0 | 1 | 0 |
| Pseudocheiridae | Ringtail possums & allies | 1/20 (5%) | 0 | 1 | 0 |
| Vombatidae | Wombats | 1/3 (33.3%) | 0 | 0 | 1 |
| **Didelphimorphia** | **Opossums** | **12/111 (10.8%)** | **0%** | **49.9%** | **50.1%** |
| Didelphidae | Opossums | 12/111 (10.8%) | 0 | 6 | 6 |
| **Lagomorpha** | **Rabbits, pikas, & hares** | **7/98 (7.1%)** | **60%** | **13.4%** | **26.6%** |
| Leporidae | Rabbits & hares | 4/67 (6%) | 4 | 0 | 0 |
| Ochotonidae | Pikas | 3/30 (10%) | 0 | 1 | 2 |
| **Dasyuromorphia** | **Carnivorous Australian marsupials** | **5/78 (6.4%)** | **0%** | **0%** | **100%** |
| Dasyuridae | Carnivorous Australian marsupials | 5/76 (6.6%) | 0 | 0 | 5 |
| **Afrosoricida** | **Tenrecs, otter shrews, & golden moles** | **12/55 (21.8%)** | **7.7%** | **92.3%** | **0%** |
| Tenrecidae | Tenrecs | 12/31 (38.7%) | 1 | 11 | 0 |
| Chrysochloridae | Golden moles | 0/21 (0%) | | | |
| **Macroscelidea** | **Elephant shrews** | **2/30 (6.7%)** | **50.1%** | **49.9%** | **0%** |
| Macroscelididae | Elephant shrews | 2/30 (6.7%) | 1 | 1 | 0 |

| | | | | | |
|---|---|---|---|---|---|
| **Scandentia** | **Treeshrews** | **2/24 (8.3%)** | **0%** | **50.3%** | **49.7%** |
| Tupaiidae | Treeshrews | 2/23 (8.7%) | 0 | 1 | 1 |
| **Peramelemorphia** | **Bandicoots & bilbies** | **3/23 (13%)** | **0%** | **0%** | **100%** |
| Peramelidae | Bandicoots | 3/20 (15%) | 0 | 0 | 3 |
| **Perissodactyla** | **Odd-toed ungulates** | **0/21 (0%)** | **N/A** | **N/A** | **N/A** |
| Equidae | Horses, zebras, & asses | 0/12 (0%) | | | |
| Rhinocerotidae | Rhinoceroses | 0/5 (0%) | | | |
| Tapiridae | Tapirs | 0/4 (0%) | | | |
| **Cingulata** | **Armadillos** | **3/20 (15%)** | **47.7%** | **0%** | **52.3%** |
| Chlamyphoridae | Armadillos | 2/13 (15%) | 1 | 0 | 1 |
| Dasypodidae | Armadillos | 1/7 (14.3%) | 0 | 1 | 0 |
| **Pilosa** | **Anteaters & sloths** | **4/10 (40%)** | **23.9%** | **76.1%** | **0%** |
| Bradypodidae | Three-toed sloths | 2/4 (50%) | 1 | 1 | 0 |
| Choloepodidae | Two-toed sloths | 1/1 (100%) | 0 | 1 | 0 |
| Cylcopedidae | Silky anteater | 0/1 (0%) | | | |
| Myrmecophagidae | Anteaters | 1/3 (33%) | 0 | 1 | 0 |

**Summary of sample sizes achieved for each order (in order of species richness) and family, the estimated rates of dimorphism & monomorphism in each order (percentages in colored columns, representing rates calibrated by species richness in each family) and the distribution of dimorphism in the whole dataset by family (numbers in colored columns, representing the number of species in the dataset in each category). We only included orders and families with at least 10 species, with the exception of Pilosa, for which no family had more than 4 species, and a few cases where data from families with <10 species were included to help achieve 5% representation for the order. Under Primates, Cebidae includes Aotidae and Callitrichidae, following Burgin et al. (2018).**

**Data S1. (separate file)**

**Species, sample sizes, and sources for all data used in our analyses. The entire dataset was used to calculate the minimum sample size required for inclusion (see Fig. S1), after which only the rows with a minimum sample size of 9 for each sex (i.e., in both the '$N_{males}$' and '$N_{females}$' columns) were used for all other analyses. A more detailed datasheet will be published upon acceptance of this manuscript.**
