## Supplementary material for "New estimates indicate that males are not larger than females in most mammals": Data Sources

| Order | Family | Species | Scientific Name | $N_{males}$ | $N_{females}$ | Source |
|---|---|---|---|---|---|---|
| Afrosoricida | Chrysochloridae | Namib desert golden mole | *Eremitalpa granti namibensis* | 4 | 8 | L. J. Fielden, thesis, University of Natal (1989). |
| Afrosoricida | Tenrecidae | Lesser hedgehog tenrec | *Echinops telfairi* | 24 | 15 | P. M. Kappeler, C. L. Nunn, A. Q. Vining, S. M. Goodman, Evolutionary dynamics of sexual size dimorphism in non-volant mammals following their independent colonization of Madagascar. *Sci. Rep.* **9**, 1–14 (2019). |
| Afrosoricida | Tenrecidae | Large-eared tenrec | *Geogale aurita* | 17 | 15 | P. M. Kappeler, C. L. Nunn, A. Q. Vining, S. M. Goodman, Evolutionary dynamics of sexual size dimorphism in non-volant mammals following their independent colonization of Madagascar. *Sci. Rep.* **9**, 1–14 (2019). |
| Afrosoricida | Tenrecidae | Highland streaked tenrec | *Hemicentetes nigriceps* | 3 | 3 | P. M. Kappeler, C. L. Nunn, A. Q. Vining, S. M. Goodman, Evolutionary dynamics of sexual size dimorphism in non-volant mammals following their independent colonization of Madagascar. *Sci. Rep.* **9**, 1–14 (2019). |
| Afrosoricida | Tenrecidae | Lowland streaked tenrec | *Hemicentetes semispinosus* | 24 | 5 | P. M. Kappeler, C. L. Nunn, A. Q. Vining, S. M. Goodman, Evolutionary dynamics of sexual size dimorphism in non-volant mammals following their independent colonization of Madagascar. *Sci. Rep.* **9**, 1–14 (2019). |
| Afrosoricida | Tenrecidae | Short-tailed shrew tenrec | *Microgale brevicaudata* | 42 | 8 | P. M. Kappeler, C. L. Nunn, A. Q. Vining, S. M. Goodman, Evolutionary dynamics of sexual size dimorphism in non-volant mammals following their independent colonization of Madagascar. *Sci. Rep.* **9**, 1–14 (2019). |

| | | | | | | |
|---|---|---|---|---|---|---|
| Afrosoricida | Tenrecidae | Cowan's shrew tenrec | *Microgale cowani* | 41 | 58 | P. M. Kappeler, C. L. Nunn, A. Q. Vining, S. M. Goodman, Evolutionary dynamics of sexual size dimorphism in non-volant mammals following their independent colonization of Madagascar. *Sci. Rep.* **9**, 1–14 (2019). |
| Afrosoricida | Tenrecidae | Dobson's shrew tenrec | *Microgale dobsoni* | 23 | 18 | P. M. Kappeler, C. L. Nunn, A. Q. Vining, S. M. Goodman, Evolutionary dynamics of sexual size dimorphism in non-volant mammals following their independent colonization of Madagascar. *Sci. Rep.* **9**, 1–14 (2019). |
| Afrosoricida | Tenrecidae | Drouhard's shrew tenrec | *Microgale drouhardi* | 30 | 23 | P. M. Kappeler, C. L. Nunn, A. Q. Vining, S. M. Goodman, Evolutionary dynamics of sexual size dimorphism in non-volant mammals following their independent colonization of Madagascar. *Sci. Rep.* **9**, 1–14 (2019). |
| Afrosoricida | Tenrecidae | Pale shrew tenrec | *Microgale fotsifotsy* | 29 | 8 | P. M. Kappeler, C. L. Nunn, A. Q. Vining, S. M. Goodman, Evolutionary dynamics of sexual size dimorphism in non-volant mammals following their independent colonization of Madagascar. *Sci. Rep.* **9**, 1–14 (2019). |
| Afrosoricida | Tenrecidae | Gracile shrew tenrec | *Microgale gracilis* | 14 | 7 | P. M. Kappeler, C. L. Nunn, A. Q. Vining, S. M. Goodman, Evolutionary dynamics of sexual size dimorphism in non-volant mammals following their independent colonization of Madagascar. *Sci. Rep.* **9**, 1–14 (2019). |
| Afrosoricida | Tenrecidae | Grandidier's shrew tenrec | *Microgale grandidieri* | 11 | 5 | P. M. Kappeler, C. L. Nunn, A. Q. Vining, S. M. Goodman, Evolutionary dynamics of sexual size dimorphism in non-volant mammals following their independent colonization of Madagascar. *Sci. Rep.* **9**, 1–14 (2019). |
| Afrosoricida | Tenrecidae | Naked-nosed shrew tenrec | *Microgale gymnorhyncha* | 17 | 23 | P. M. Kappeler, C. L. Nunn, A. Q. Vining, S. M. Goodman, Evolutionary dynamics of sexual size dimorphism in non-volant mammals following their independent colonization of Madagascar. *Sci. Rep.* **9**, 1–14 (2019). |

| | | | | | | |
|---|---|---|---|---|---|---|
| Afrosoricida | Tenrecidae | Montane shrew tenrec | *Microgale monticola* | 10 | 6 | P. M. Kappeler, C. L. Nunn, A. Q. Vining, S. M. Goodman, Evolutionary dynamics of sexual size dimorphism in non-volant mammals following their independent colonization of Madagascar. *Sci. Rep.* **9**, 1–14 (2019). |
| Afrosoricida | Tenrecidae | Pygmy shrew tenrec | *Microgale parvula* | 40 | 27 | P. M. Kappeler, C. L. Nunn, A. Q. Vining, S. M. Goodman, Evolutionary dynamics of sexual size dimorphism in non-volant mammals following their independent colonization of Madagascar. *Sci. Rep.* **9**, 1–14 (2019). |
| Afrosoricida | Tenrecidae | Shrew-toothed shrew tenrec | *Microgale soricoides* | 28 | 29 | P. M. Kappeler, C. L. Nunn, A. Q. Vining, S. M. Goodman, Evolutionary dynamics of sexual size dimorphism in non-volant mammals following their independent colonization of Madagascar. *Sci. Rep.* **9**, 1–14 (2019). |
| Afrosoricida | Tenrecidae | Taiva shrew tenrec | *Microgale taiva* | 39 | 32 | P. M. Kappeler, C. L. Nunn, A. Q. Vining, S. M. Goodman, Evolutionary dynamics of sexual size dimorphism in non-volant mammals following their independent colonization of Madagascar. *Sci. Rep.* **9**, 1–14 (2019). |
| Afrosoricida | Tenrecidae | Talzac's shrew tenrec | *Microgale talazaci* | 15 | 16 | P. M. Kappeler, C. L. Nunn, A. Q. Vining, S. M. Goodman, Evolutionary dynamics of sexual size dimorphism in non-volant mammals following their independent colonization of Madagascar. *Sci. Rep.* **9**, 1–14 (2019). |
| Afrosoricida | Tenrecidae | Thomas's shrew tenrec | *Microgale thomasi* | 34 | 19 | P. M. Kappeler, C. L. Nunn, A. Q. Vining, S. M. Goodman, Evolutionary dynamics of sexual size dimorphism in non-volant mammals following their independent colonization of Madagascar. *Sci. Rep.* **9**, 1–14 (2019). |
| Afrosoricida | Tenrecidae | Mole-like rice tenrec | *Oryzorictes hova* | 27 | 6 | P. M. Kappeler, C. L. Nunn, A. Q. Vining, S. M. Goodman, Evolutionary dynamics of sexual size dimorphism in non-volant mammals following their independent colonization of Madagascar. *Sci. Rep.* **9**, 1–14 (2019). |

| | | | | | | |
|---|---|---|---|---|---|---|
| Afrosoricida | Tenrecidae | Greater hedgehog tenrec | *Setifer setosus* | 52 | 46 | P. M. Kappeler, C. L. Nunn, A. Q. Vining, S. M. Goodman, Evolutionary dynamics of sexual size dimorphism in non-volant mammals following their independent colonization of Madagascar. *Sci. Rep.* **9**, 1–14 (2019). |
| Afrosoricida | Tenrecidae | Tailless tenrec | *Tenrec ecaudatus* | 4 | 6 | P. M. Kappeler, C. L. Nunn, A. Q. Vining, S. M. Goodman, Evolutionary dynamics of sexual size dimorphism in non-volant mammals following their independent colonization of Madagascar. *Sci. Rep.* **9**, 1–14 (2019). |
| Artiodactyl a | Bovidae | Impala | *Aepyceros melampus* | 10 | 9 | I. G. Anderson, Mass and body measurements of impala *Aepyceros melampus*, from a game ranch. *South African J. Wildl. Res.* **12**, 76–78 (1982). |
| Artiodactyl a | Bovidae | Red hartebeest | *Alcelaphus buselaphus caama* | 13 | 16 | L. C. Hoffman, K. Smit, N. Muller, Chemical characteristics of red hartebeest (*Alcelaphus buselaphus caama*) meat. *South African J. Anim. Sci.* **40**, 221–228 (2010). |
| Artiodactyl a | Bovidae | Springbok | *Antidorcas marsupialis* | 13 | 24 | M. Kroucamp, thesis, University of Stellenbosch (2004). |
| Artiodactyl a | Bovidae | Bison | *Bison bison* | 29 | 23 | J. Berger, M. Peacock, Variability in size-weight relationships of *Bison bison*. *J. Mammal.* **69**, 618–624 (1988). |
| Artiodactyl a | Bovidae | Spanish ibex | *Capra pyrenaica* | 73 | 12 3 | J. E. Granados, J. M. Perez, R. C. Soriguer, P. Fandos, I. Ruiz-Martinez, On the biometry of the Spanish ibex, *Capra pyrenaica*, from Sierra Nevada (southern Spain). *Folia Zool.* **46**, 9–14 (1997). |
| Artiodactyl a | Bovidae | Japanese serow | *Capricornis crispus* | 29 2 | 27 9 | S. Miura, Body and horn growth patterns in the Japanese serow, *Capricornis crispus*. *J. Mammal. Soc. Japan*. **11**, 1–13 (1986). |

| | | | | | | |
|---|---|---|---|---|---|---|
| Artiodactyla | Bovidae | Blesbok | *Damaliscus dorcas phillipsi* | 4 | 4 | L. Van Zyl, A. V. Ferreira, Physical and chemical carcass composition of springbok (*Antidorcas marsupialis*), blesbok (*Damaliscus dorcas phillipsi*) and impala (*Aepyceros melampus*). *Small Rumin. Res.* **53**, 103–109 (2004). |
| Artiodactyla | Bovidae | Dorcas gazelle | *Gazella dorcas* | 8 | 16 | T. Wronski, M. A. Sandouka, M. Plath, P. Cunningham, Differences in sexual dimorphism among four gazelle taxa (*Gazella* spp.) in the Middle East. *Anim. Biol.* **60**, 395–412 (2010). |
| Artiodactyla | Bovidae | Mountain gazelle | *Gazella gazella* | 39 | 94 | T. Wronski, M. A. Sandouka, M. Plath, P. Cunningham, Differences in sexual dimorphism among four gazelle taxa (*Gazella* spp.) in the Middle East. *Anim. Biol.* **60**, 395–412 (2010). |
| Artiodactyla | Bovidae | Red-fronted gazelle | *Gazella rufrons kanuri* | 59 | 36 | A. C. Nchanji, F. O. Amubode, The physical and morphological characteristics of the red-fronted gazelle (*Gazella rufifrons kanuri* Gray 1846) in Waza National Park, Cameroon. *J. Zool.* **256**, 505–509 (2002). |
| Artiodactyla | Bovidae | Sand gazelle | *Gazella subgutturosa marica* | 150 | 229 | T. Wronski, M. A. Sandouka, M. Plath, P. Cunningham, Differences in sexual dimorphism among four gazelle taxa (*Gazella* spp.) in the Middle East. *Anim. Biol.* **60**, 395–412 (2010). |
| Artiodactyla | Bovidae | Soay sheep | *Ovis aries* | 13 | 13 | F. J. Pérez-Barbería, E. Pérez-Fernández, E. Robertson, B. Alvarez-Enríquez, Does the Jarman-Bell principle at intra-specific level explain sexual segregation in polygynous ungulates? Sex differences in forage digestibility in Soay sheep. *Oecologia*. **157**, 21–30 (2008). |
| Artiodactyla | Bovidae | Bighorn sheep | *Ovis canadensis* | 45 | 44 | M. Festa-Bianchet, J. T. Jorgenson, W. J. King, K. G. Smith, W. D. Wishart, The development of sexual dimorphism: Seasonal and lifetime mass changes in bighorn sheep. *Can. J. Zool.* **74**, 330–342 (1996). |

| | | | | | | |
|---|---|---|---|---|---|---|
| Artiodactyl a | Bovidae | Mediterran ean mouflon | *Ovis gmelini* | 17 5 | 62 6 | G. Bourgoin, P. Marchand, A. J. M. Hewison, K. E. Ruckstuhl, M. Garel, Social behaviour as a predominant driver of sexual, age-dependent and reproductive segregation in Mediterranean mouflon. *Anim. Behav.* **136**, 87–100 (2018). |
| Artiodactyl a | Bovidae | Alpine chamois | *Rupicapra rupicapra* | 48 | 37 | B. Bassano, A. Perrone, A. Von Hardenberg, Body weight and horn development in Alpine chamois, *Rupicapra rupicapra* (Bovidae, Caprinae). *Mammalia*. **67**, 65–73 (2003). |
| Artiodactyl a | Bovidae | Greater kudu | *Tragelaphus strepsiceros* | 49 | 12 | N. Fuentes, thesis, Durham University (2021). |
| Artiodactyl a | Cervidae | Moose | *Alces alces andersoni* | 10 7 | 43 | W. F. Jensen, J. R. Smith, J. V McKenzie, J. J. Maskey Jr., R. E. Johnson, Mass, morphology, and growth rates of moose in North Dakota. *Alces*. **49**, 1–15 (2013). |
| Artiodactyl a | Cervidae | Roe deer | *Capreolus capreolus* | 10 1 | 11 2 | J. Ritz, K. Hofer, E. Hofer, K. Hackländer, D. Immekus, D. Codron, M. Clauss, Forestomach pH in hunted roe deer (*Capreolus capreolus*) in relation to forestomach region, time of measurement and supplemental feeding and comparison among wild ruminant species. *Eur. J. Wildl. Res.* **59**, 505–517 (2013). |
| Artiodactyl a | Cervidae | Reeves' muntjac | *Muntiacus reevesi* | 72 | 18 | N. G. Chapman, M. Furlong, S. Harris, Reproductive strategies and the influence of date of birth on growth and sexual development of an aseasonally-breeding ungulate: Reeves’ muntjac (*Muntiacus reevesi*). *J. Zool.* **241**, 551–570 (1997). |
| Artiodactyl a | Cervidae | White-tailed deer | *Odocoileus virginianus* | 71 | 88 | L. Lesage, M. Crête, J. Huot, J. P. Ouellet, Evidence for a trade-off between growth and body reserves in northern white-tailed deer. *Oecologia*. **126**, 30–41 (2001). |

| | | | | | | |
|---|---|---|---|---|---|---|
| Artiodactyla | Cervidae | Pampas deer | *Ozotoceros bezoarticus* | 5 | 9 | W. Perez, R. Ungerfeld, Gross anatomy of the stomach of the pampas deer, *Ozotoceros bezoarticus* (Artiodactyla: Cervidae). *Zoologia*. **29**, 337–342 (2012). |
| Artiodactyla | Cervidae | Reindeer | *Rangifer tarandus* | 10 | 31 | E. Pintus, S. Uccheddu, K. H. Røed, J. P. Gonzaléz, J. Carranza, M. Nieminen, Ø. Holand, Flexible mating tactics and associated reproductive effort during the rutting season in male reindeer (*Rangifer tarandus*, L. 1758). *Curr. Zool.* **61**, 802–810 (2015). |
| Artiodactyla | Delphinidae | Risso's dolphin | *Grampus griseus* | 3 | 3 | D. Bloch, G. Desportes, P. Harvey, C. Lockyer, B. Mikkelsen, Life history of Risso's dolphin (*Grampus griseus*) (G. Cuvier, 1812) in the Faroe Islands. *Aquat. Mamm.* **38**, 250–266 (2012). |
| Artiodactyla | Delphinidae | Striped dolphin | *Stenella coeruleoalba* | 15 | 10 | F. Monaci, A. Borrel, C. Leonzio, L. Marsili, N. Calzada, Trace elements in striped dolphins (*Stenella coeruleoalba*) from the western Mediterranean. *Environ. Pollut.* **99**, 61–68 (1998). |
| Artiodactyla | Delphinidae | Bottle nose dolphin | *Tursiops truncatus* | 14 | 18 | A. Fahlman, M. Brodsky, R. Wells, K. McHugh, J. Allen, A. Barleycorn, J. C. Sweeney, D. Fauquier, M. Moore, Field energetics and lung function in wild bottlenose dolphins, *Tursiops truncatus*, in Sarasota Bay Florida. *R. Soc. Open Sci.* **5** (2018), doi:10.1098/rsos.171280. |
| Artiodactyla | Phocoenidae | Harbour porpoise | *Phocoena phocoena* | 62 | 62 | A. J. Read, K. A. Tolley, Postnatal growth and allometry of harbour porpoises from the Bay of Fundy. *Can. J. Zool.* **75**, 122–130 (1997). |
| Artiodactyla | Phocoenidae | Burmeister's porpoise | *Phocoena spinipinnis* | 9 | 11 | J. C. Reyes, K. Van Waerebeek, "Aspects of the biology of Burmeister's porpoise from Peru" (1995). |

| | | | | | | |
|---|---|---|---|---|---|---|
| Artiodactyla | Pontoporiidae | Franciscana dolphin | *Pontoporia blainvillei* | 129 | 93 | S. Botta, M. M. C. Muelbert, E. R. Secchi, Morphometric relationships of franciscana dolphin, *Pontoporia blainvillei* (Cetacea), off Rio Grande do Sul coast, southern Brazil. *Lat. Am. J. Aquat. Mamm.* **5**, 117–123 (2006). |
| Artiodactyla | Suidae | Warthog | *Phacochoerus athiopicus* | 13 | 16 | D. G. Booyse, B. A. Dehority, Gastrointestinal protozoa and digestive tract parameters of wild South African warthogs (*Phacochoerus aethiopicus*). *African Zool.* **47**, 373 (2012). |
| Artiodactyla | Suidae | Wild boar | *Sus scrofa* | 40 | 76 | S. Focardi, J. M. Gaillard, F. Ronchi, S. Rossi, Survival of wild boars in a variable environment: Unexpected life-history variation in an unusual ungulate. *J. Mammal.* **89**, 1113–1123 (2008). |

| | | | | | | |
|---|---|---|---|---|---|---|
| Carnivora | Eupleridae | Fossa | *Cryptoprocta ferox* | 17 | 11 | C. E. Hawkins, P. A. Racey, Low population density of a tropical forest carnivore, *Cryptoprocta ferox*: Implications for protected area management. *Oryx*. **39**, 35–43 (2005). |
| Carnivora | Eupleridae | Ring-tailed vontsira | *Galidia elegans* | 10 | 2 | P. M. Kappeler, C. L. Nunn, A. Q. Vining, S. M. Goodman, Evolutionary dynamics of sexual size dimorphism in non-volant mammals following their independent colonization of Madagascar. *Sci. Rep.* **9**, 1–14 (2019). |
| Carnivora | Eupleridae | Grandidier's mongoose | *Galidictis grandidieri* | 20 | 10 | P. M. Kappeler, C. L. Nunn, A. Q. Vining, S. M. Goodman, Evolutionary dynamics of sexual size dimorphism in non-volant mammals following their independent colonization of Madagascar. *Sci. Rep.* **9**, 1–14 (2019). |
| Carnivora | Eupleridae | Narrow-striped mongoose | *Mungotictis decemlineata* | 22 | 19 | P. M. Kappeler, C. L. Nunn, A. Q. Vining, S. M. Goodman, Evolutionary dynamics of sexual size dimorphism in non-volant mammals following their independent colonization of Madagascar. *Sci. Rep.* **9**, 1–14 (2019). |

| | | | | | | |
|---|---|---|---|---|---|---|
| Carnivora | Felidae | Margay | *Leopardus wiedii* | 7 | 6 | F. Gonçalves, R. S. Bovendorp, G. Beca, C. Bello, R. Costa-Pereira, R. L. Muylaert, R. R. Rodarte, N. Villar, R. Souza, M. E. Graipel, J. J. Cherem, D. Faria, J. Baumgarten, M. R. Alvarez, E. M. Vieira, N. Cáceres, R. Pardini, Y. L. R. Leite, L. P. Costa, M. A. R. Mello, E. Fischer, F. C. Passos, L. H. Varzinczak, J. A. Prevedello, A. P. Cruz-Neto, F. Carvalho, A. R. Percequillo, A. Paviolo, A. Nava, J. M. B. Duarte, N. U. de la Sancha, E. Bernard, R. G. Morato, J. F. Ribeiro, R. G. Becker, G. Paise, P. S. Tomasi, F. Vélez-Garcia, G. L. Melo, J. Sponchiado, F. Cerezer, M. A. S. Barros, A. Q. S. de Souza, C. C. dos Santos, G. A. F. Giné, P. Kerches-Rogeri, M. M. Weber, G. Ambar, L. V. Cabrera-Martinez, A. Eriksson, M. Silveira, C. F. Santos, L. Alves, E. Barbier, G. C. Rezende, G. S. T. Garbino, É. O. Rios, A. Silva, A. T. A. Nascimento, R. S. de Carvalho, A. Feijó, J. Arrabal, I. Agostini, D. Lamattina, S. Costa, E. Vanderhoeven, F. R. de Melo, P. de Oliveira Laroque, L. Jerusalinsky, M. M. Valença-Montenegro, A. B. Martins, G. Ludwig, R. B. de Azevedo, A. Anzóategui, M. X. da Silva, M. Figuerêdo Duarte Moraes, A. Vogliotti, A. Gatti, T. Püttker, C. S. Barros, T. K. Martins, A. Keuroghlian, D. P. Eaton, C. L. Neves, M. S. Nardi, C. Braga, P. R. Gonçalves, A. C. Srbek-Araujo, P. Mendes, J. A. de Oliveira, F. A. M. Soares, P. A. Rocha, P. Crawshaw, M. C. Ribeiro, M. Galetti, Atlantic Mammal Traits: a data set of morphological traits of mammals in the Atlantic Forest of South America. *Ecology*. **99**, 498 (2018). |
| Carnivora | Felidae | Lion | *Panthera leo* | 4 | 6 | M. Valeix, A. J. Loveridge, D. W. Macdonald, Influence of prey dispersion on territory and group size of African lions: A test of the resource dispersion hypothesis. *Ecology*. **93**, 2490–2496 (2012). |

| | | | | | | |
|---|---|---|---|---|---|---|
| Carnivora | Herpestidae | Yellow mongoose | *Cynictis penicillata* | 7 | 5 | O. A. Mbatyoti, thesis, University of Fort Hare (2012). |
| Carnivora | Herpestidae | Egyptian mongoose | *Herpestes ichneumon* | 9 | 12 | F. Palomares, M. Delibes, Some physical and population characteristics of Egyptian mongooses (*Herpestes ichneumon* L., 1758) in southwestern Spain. *Z. Saugetierkd.* **57** (1992), pp. 94–99. |
| Carnivora | Herpestidae | Indian mongoose | *Herpestes javanicus* | 72 | 56 | W. S. T. Hays, D. Simberloff, A morphometric trend linked to male sociality in the small Indian mongoose *Herpestes javanicus* in Hawaii. *Acta Theriol. (Warsz).* **51**, 303–310 (2006). |
| Carnivora | Mephitidae | Molina's hog-nosed skunk | *Conepatus chinga* | 7 | 11 | D. F. Castillo, M. Lucherini, E. M. L. Vidal, C. Manfredi, E. B. Casanave, Spatial organization of Molina’s hog-nosed skunk (*Conepatus chinga*) in two landscapes of the Pampas grassland of Argentina. *Can. J. Zool.* **89**, 229–238 (2011). |
| Carnivora | Mephitidae | Eastern spotted skunk | *Spilogale putorius* | 94 | 38 | A. E. Kinlaw, L. M. Ehrhart, P. D. Doerr, Spotted skunks (*Spilogale putorius ambarvalis*) trapped at Canaveral National Seashore and Merrit Island, Florida. *Florida F. Nat.* **23**, 57–86 (1995). |

| | | | | | | |
|---|---|---|---|---|---|---|
| Carnivora | Mustelidae | Beech marten | *Martes foina* | 84 | 82 | S. Ruette, J. Larroque, M. Albaret, J. M. Vandel, S. Devillard, Quantifying the age- and sex-dependent morphological variation in two syntopic mustelids: *Martes martes* and *Martes foina*. *Mamm. Biol.* **80**, 414–423 (2015). |
| Carnivora | Mustelidae | European pine marten | *Martes martes* | 47 | 26 | S. Ruette, J. Larroque, M. Albaret, J. M. Vandel, S. Devillard, Quantifying the age- and sex-dependent morphological variation in two syntopic mustelids: *Martes martes* and *Martes foina*. *Mamm. Biol.* **80**, 414–423 (2015). |
| Carnivora | Mustelidae | Honey badger | *Mellivora capensis* | 17 | 13 | C. M. Begg, K. S. Begg, J. T. Du Toit, M. G. L. Mills, Sexual and seasonal variation in the diet and foraging behaviour of a sexually dimorphic carnivore, the honey badger (*Mellivora capensis*). *J. Zool.* **260**, 301–316 (2003). |
| Carnivora | Mustelidae | American mink | *Neovison vison* | 6 | 10 | L. A. Harrington, G. C. Hays, L. Fasola, A. L. Harrington, D. Righton, D. W. MacDonald, Dive performance in a small-bodied, semi-aquatic mammal in the wild. *J. Mammal.* **93**, 198–210 (2012). |
| Carnivora | Otariidae | Australian fur seal | *Arctocephalus pusillus doriferus* | 6 | 6 | L. Kernaléguen, Y. Cherel, T. C. Knox, A. M. M. Baylis, J. P. Y. Arnould, Sexual niche segregation and gender-specific individual specialisation in a highly dimorphic marine mammal. *PLoS One*. **10**, 1–15 (2015). |
| Carnivora | Phocidae | Northern elephant seal | *Mirounga angustirostris* | 32 | 94 | S. S. Kienle, A. S. Friedlaender, D. E. Crocker, R. S. Mehta, D. P. Costa, Trade-offs between foraging reward and mortality risk drive sex-specific foraging strategies in sexually dimorphic northern elephant seals. *R. Soc. Open Sci.* **9** (2022), doi:10.1098/rsos.210522. |
| Carnivora | Procyonidae | White-nosed coati | *Nasua narica* | 51 | 37 | M. E. Gompper, Sociality and asociality in white-nosed coatis (*Nasua narica*): Foraging costs and benefits. *Behav. Ecol.* **7**, 254–263 (1996). |

| | | | | | | |
|---|---|---|---|---|---|---|
| Carnivora | Procyonidae | South American coati | *Nasua nasua* | 50 | 90 | F. Gonçalves, R. S. Bovendorp, G. Beca, C. Bello, R. Costa-Pereira, R. L. Muylaert, R. R. Rodarte, N. Villar, R. Souza, M. E. Graipel, J. J. Cherem, D. Faria, J. Baumgarten, M. R. Alvarez, E. M. Vieira, N. Cáceres, R. Pardini, Y. L. R. Leite, L. P. Costa, M. A. R. Mello, E. Fischer, F. C. Passos, L. H. Varzinczak, J. A. Prevedello, A. P. Cruz-Neto, F. Carvalho, A. R. Percequillo, A. Paviolo, A. Nava, J. M. B. Duarte, N. U. de la Sancha, E. Bernard, R. G. Morato, J. F. Ribeiro, R. G. Becker, G. Paise, P. S. Tomasi, F. Vélez-Garcia, G. L. Melo, J. Sponchiado, F. Cerezer, M. A. S. Barros, A. Q. S. de Souza, C. C. dos Santos, G. A. F. Giné, P. Kerches-Rogeri, M. M. Weber, G. Ambar, L. V. Cabrera-Martinez, A. Eriksson, M. Silveira, C. F. Santos, L. Alves, E. Barbier, G. C. Rezende, G. S. T. Garbino, É. O. Rios, A. Silva, A. T. A. Nascimento, R. S. de Carvalho, A. Feijó, J. Arrabal, I. Agostini, D. Lamattina, S. Costa, E. Vanderhoeven, F. R. de Melo, P. de Oliveira Laroque, L. Jerusalinsky, M. M. Valença-Montenegro, A. B. Martins, G. Ludwig, R. B. de Azevedo, A. Anzóategui, M. X. da Silva, M. Figuerêdo Duarte Moraes, A. Vogliotti, A. Gatti, T. Püttker, C. S. Barros, T. K. Martins, A. Keuroghlian, D. P. Eaton, C. L. Neves, M. S. Nardi, C. Braga, P. R. Gonçalves, A. C. Srbek-Araujo, P. Mendes, J. A. de Oliveira, F. A. M. Soares, P. A. Rocha, P. Crawshaw, M. C. Ribeiro, M. Galetti, Atlantic Mammal Traits: a data set of morphological traits of mammals in the Atlantic Forest of South America. *Ecology*. **99**, 498 (2018). |

| | | | | | | |
|---|---|---|---|---|---|---|
| Carnivora | Procyonidae | Crab-eating raccoon | *Procyon cancrivorus* | 9 | 4 | F. Gonçalves, R. S. Bovendorp, G. Beca, C. Bello, R. Costa-Pereira, R. L. Muylaert, R. R. Rodarte, N. Villar, R. Souza, M. E. Graipel, J. J. Cherem, D. Faria, J. Baumgarten, M. R. Alvarez, E. M. Vieira, N. Cáceres, R. Pardini, Y. L. R. Leite, L. P. Costa, M. A. R. Mello, E. Fischer, F. C. Passos, L. H. Varzinczak, J. A. Prevedello, A. P. Cruz-Neto, F. Carvalho, A. R. Percequillo, A. Paviolo, A. Nava, J. M. B. Duarte, N. U. de la Sancha, E. Bernard, R. G. Morato, J. F. Ribeiro, R. G. Becker, G. Paise, P. S. Tomasi, F. Vélez-Garcia, G. L. Melo, J. Sponchiado, F. Cerezer, M. A. S. Barros, A. Q. S. de Souza, C. C. dos Santos, G. A. F. Giné, P. Kerches-Rogeri, M. M. Weber, G. Ambar, L. V. Cabrera-Martinez, A. Eriksson, M. Silveira, C. F. Santos, L. Alves, E. Barbier, G. C. Rezende, G. S. T. Garbino, É. O. Rios, A. Silva, A. T. A. Nascimento, R. S. de Carvalho, A. Feijó, J. Arrabal, I. Agostini, D. Lamattina, S. Costa, E. Vanderhoeven, F. R. de Melo, P. de Oliveira Laroque, L. Jerusalinsky, M. M. Valença-Montenegro, A. B. Martins, G. Ludwig, R. B. de Azevedo, A. Anzóategui, M. X. da Silva, M. Figuerêdo Duarte Moraes, A. Vogliotti, A. Gatti, T. Püttker, C. S. Barros, T. K. Martins, A. Keuroghlian, D. P. Eaton, C. L. Neves, M. S. Nardi, C. Braga, P. R. Gonçalves, A. C. Srbek-Araujo, P. Mendes, J. A. de Oliveira, F. A. M. Soares, P. A. Rocha, P. Crawshaw, M. C. Ribeiro, M. Galetti, Atlantic Mammal Traits: a data set of morphological traits of mammals in the Atlantic Forest of South America. *Ecology*. **99**, 498 (2018). |
| Carnivora | Viverridae | Bushy-tailed mongoose | *Bdeogale crassicauda* | 3 | 2 | A. Martinoli, D. Preatoni, V. Galanti, P. Codipietro, M. Kilewo, C. A. R. Fernandes, L. A. Wauters, G. Tosi, Species richness and habitat use of small carnivores in the Arusha National Park (Tanzania). *Biodivers. Conserv.* **15**, 1729–1744 (2006). |

| | | | | | | |
|---|---|---|---|---|---|---|
| Carnivora | Viverridae | Common slender mongoose | *Galerella sanguineus* | 3 | 3 | A. Martinoli, D. Preatoni, V. Galanti, P. Codipietro, M. Kilewo, C. A. R. Fernandes, L. A. Wauters, G. Tosi, Species richness and habitat use of small carnivores in the Arusha National Park (Tanzania). *Biodivers. Conserv.* **15**, 1729–1744 (2006). |
| Carnivora | Viverridae | Common genet | *Genetta genetta* | 32 | 35 | C. Rodriguez-Refojos, I. Zuberogoitia, L. M. Rosalino, M. J. Santos, M. Santos-Reis, J. Zabala, D. Camps, Geographical and sexual differences in body size of common genets, *Genetta genetta* (Viverridae, Carnivora), in south-western Europe (Iberian Peninsula). *Folia Zool.* **60**, 54–62 (2011). |
| Carnivora | Viverridae | Rusty-spotted genet | *Genetta maculata* | 2 | 2 | A. Martinoli, D. Preatoni, V. Galanti, P. Codipietro, M. Kilewo, C. A. R. Fernandes, L. A. Wauters, G. Tosi, Species richness and habitat use of small carnivores in the Arusha National Park (Tanzania). *Biodivers. Conserv.* **15**, 1729–1744 (2006). |
| Carnivora | Viverridae | Masked palm civet | *Paguma larvata* | 16 | 17 | Y. Zhou, C. Newman, F. Palomares, S. Zhang, Z. Xie, D. W. Macdonald, Spatial organization and activity patterns of the masked palm civet (*Paguma larvata*) in central-south China. *J. Mammal.* **95**, 534–542 (2014). |

| | | | | | | |
|---|---|---|---|---|---|---|
| Chiroptera | Hipposideridae | Ashy round-leaf bat | *Hipposideros cineraceus* | 25 | 26 | B. Douangboubpha, S. Bumrungsri, P. Soisook, S. W. Murray, S. J. Puechmaille, C. Satasook, S. S. Hla Bu, D. L. Harrison, P. J. J. Bates, A taxonomic review of *Hipposideros halophyllus*, with additional information on *H. ater* and *H. cineraceus* (Chiroptera: Hipposideridae) from Thailand and Myanmar. *Acta Chiropterologica*. **12**, 29–50 (2010). |
| Chiroptera | Hipposideridae | Thailand roundleaf bat | *Hipposideros halophyllus* | 14 | 12 | B. Douangboubpha, S. Bumrungsri, P. Soisook, S. W. Murray, S. J. Puechmaille, C. Satasook, S. S. Hla Bu, D. L. Harrison, P. J. J. Bates, A taxonomic review of *Hipposideros halophyllus*, with additional information on *H. ater* and *H. cineraceus* (Chiroptera: Hipposideridae) from Thailand and Myanmar. *Acta Chiropterologica*. **12**, 29–50 (2010). |
| Chiroptera | Hipposideridae | Intermediate roundleaf bat | *Hipposideros larvatus* | 30 | 21 | A. Thabah, S. J. Rossiter, T. Kingston, S. Zhang, S. Parsons, K. M. Mya, A. Zubaid, G. Jones, Genetic divergence and echolocation call frequency in cryptic species of *Hipposideros larvatus s.l.* (Chiroptera: Hipposideridae) from the Indo-Malayan region. *Biol. J. Linn. Soc.* **88**, 119–130 (2006). |
| Chiroptera | Hipposideridae | Noack's roundleaf bat | *Hipposideros ruber* | 20 | 27 | D. Russo, G. Maglio, A. Rainho, C. F. J. Meyer, J. M. Palmeirim, Out of the dark: Diurnal activity in the bat *Hipposideros ruber* on São Tomé island (West Africa). *Mamm. Biol.* **76**, 701–708 (2011). |
| Chiroptera | Miniopteridae | Least long-fingered bat | *Miniopterus minor* | 26 | 29 | A. N. McWilliam, Mating system of the bat Miniopterus minor (Chiroptera: Vespertilionidae) in Kenya, East Africa: A lek? *Ethology*. **85**, 302–312 (1990). |
| Chiroptera | Miniopteridae | Natal long-fingered bat | *Miniopterus natalensis* | 61 | 31 | M. Pretorius, W. Markotter, T. Kearney, E. Seamark, H. Broders, M. Keith, No evidence of pre-hibernation or pre-migratory body mass gain in *Miniopterus natalensis* in north-eastern South Africa. *J. Vertebr. Biol.* **70** (2021), doi:10.25225/jvb.20088. |

| | | | | | | |
|---|---|---|---|---|---|---|
| Chiroptera | Molossidae | Wrinkle-lipped free-tailed bat | *Chaerephon plicatus* | 20 | 23 | P. Weise, G. A. Czirják, O. Lindecke, S. Bumrungsri, C. C. Voigt, Simulated bacterial infection disrupts the circadian fluctuation of immune cells in wrinkle-lipped bats (*Chaerephon plicatus*). *PeerJ*. **2017**, 1–18 (2017). |
| Chiroptera | Molossidae | Little free-tailed bat | *Chaerephon pumilus* | 6 | 6 | F. Aspetsberger, D. Brandsen, D. S. Jacobs, Geographic variation in the morphology, echolocation and diet of the little free-tailed bat, *Chaerephon pumilus* (Molossidae). *African Zool.* **38**, 245–254 (2003). |

| | | | | | | |
|---|---|---|---|---|---|---|
| Chiroptera | Molossidae | Dwarf dog-faced bat | *Molossops temminckii* | 2 | 3 | F. Gonçalves, R. S. Bovendorp, G. Beca, C. Bello, R. Costa-Pereira, R. L. Muylaert, R. R. Rodarte, N. Villar, R. Souza, M. E. Graipel, J. J. Cherem, D. Faria, J. Baumgarten, M. R. Alvarez, E. M. Vieira, N. Cáceres, R. Pardini, Y. L. R. Leite, L. P. Costa, M. A. R. Mello, E. Fischer, F. C. Passos, L. H. Varzinczak, J. A. Prevedello, A. P. Cruz-Neto, F. Carvalho, A. R. Percequillo, A. Paviolo, A. Nava, J. M. B. Duarte, N. U. de la Sancha, E. Bernard, R. G. Morato, J. F. Ribeiro, R. G. Becker, G. Paise, P. S. Tomasi, F. Vélez-Garcia, G. L. Melo, J. Sponchiado, F. Cerezer, M. A. S. Barros, A. Q. S. de Souza, C. C. dos Santos, G. A. F. Giné, P. Kerches-Rogeri, M. M. Weber, G. Ambar, L. V. Cabrera-Martinez, A. Eriksson, M. Silveira, C. F. Santos, L. Alves, E. Barbier, G. C. Rezende, G. S. T. Garbino, É. O. Rios, A. Silva, A. T. A. Nascimento, R. S. de Carvalho, A. Feijó, J. Arrabal, I. Agostini, D. Lamattina, S. Costa, E. Vanderhoeven, F. R. de Melo, P. de Oliveira Laroque, L. Jerusalinsky, M. M. Valença-Montenegro, A. B. Martins, G. Ludwig, R. B. de Azevedo, A. Anzóategui, M. X. da Silva, M. Figuerêdo Duarte Moraes, A. Vogliotti, A. Gatti, T. Püttker, C. S. Barros, T. K. Martins, A. Keuroghlian, D. P. Eaton, C. L. Neves, M. S. Nardi, C. Braga, P. R. Gonçalves, A. C. Srbek-Araujo, P. Mendes, J. A. de Oliveira, F. A. M. Soares, P. A. Rocha, P. Crawshaw, M. C. Ribeiro, M. Galetti, Atlantic Mammal Traits: a data set of morphological traits of mammals in the Atlantic Forest of South America. *Ecology*. **99**, 498 (2018). |

| | | | | | | |
|---|---|---|---|---|---|---|
| Chiroptera | Molossidae | Molossus bondae | *Molossus bondae* | 13 | 12 | D. Ocampo, K. G. Borja-Acosta, J. Lozano-Flórez, S. Cifuentes-Acevedo, E. Arbeláez-Cortés, N. J. Bayly, Á. Caguazango, B. Coral-Jaramillo, D. Cueva, F. Forero, J. P. Gómez, C. Gómez, M. A. Loaiza-Muñoz, G. A. Londoño, S. Losada-Prado, S. Pérez-Peña, H. E. Ramírez-Chaves, M. E. Rodríguez-Posada, J. Sanabria-Mejía, M. Sánchez-Martínez, V. H. Serrano-Cardozo, M. S. Sierra-Buitrago, J. Soto-Patiño, O. Acevedo-Charry, Body mass data set for 1,317 bird and 270 mammal species from Colombia. *Ecology*. **102** (2021), doi:10.1002/ecy.3273. |
| Chiroptera | Molossidae | Coiban mastiff bat | *Molossus coibensis* | 23 | 41 | Y. Gager, thesis, Universität Konstanz (2016). |
| Chiroptera | Molossidae | Velvety free-tailed bat | *Molossus molossus* | 123 | 403 | Y. Gager, thesis, Universität Konstanz (2016). |

| | | | | | | |
|---|---|---|---|---|---|---|
| Chiroptera | Mormoopidae | Wagner's mustached bat | *Pteronotus personatus* | 54 | 54 | F. Gonçalves, R. S. Bovendorp, G. Beca, C. Bello, R. Costa-Pereira, R. L. Muylaert, R. R. Rodarte, N. Villar, R. Souza, M. E. Graipel, J. J. Cherem, D. Faria, J. Baumgarten, M. R. Alvarez, E. M. Vieira, N. Cáceres, R. Pardini, Y. L. R. Leite, L. P. Costa, M. A. R. Mello, E. Fischer, F. C. Passos, L. H. Varzinczak, J. A. Prevedello, A. P. Cruz-Neto, F. Carvalho, A. R. Percequillo, A. Paviolo, A. Nava, J. M. B. Duarte, N. U. de la Sancha, E. Bernard, R. G. Morato, J. F. Ribeiro, R. G. Becker, G. Paise, P. S. Tomasi, F. Vélez-Garcia, G. L. Melo, J. Sponchiado, F. Cerezer, M. A. S. Barros, A. Q. S. de Souza, C. C. dos Santos, G. A. F. Giné, P. Kerches-Rogeri, M. M. Weber, G. Ambar, L. V. Cabrera-Martinez, A. Eriksson, M. Silveira, C. F. Santos, L. Alves, E. Barbier, G. C. Rezende, G. S. T. Garbino, É. O. Rios, A. Silva, A. T. A. Nascimento, R. S. de Carvalho, A. Feijó, J. Arrabal, I. Agostini, D. Lamattina, S. Costa, E. Vanderhoeven, F. R. de Melo, P. de Oliveira Laroque, L. Jerusalinsky, M. M. Valença-Montenegro, A. B. Martins, G. Ludwig, R. B. de Azevedo, A. Anzóategui, M. X. da Silva, M. Figuerêdo Duarte Moraes, A. Vogliotti, A. Gatti, T. Püttker, C. S. Barros, T. K. Martins, A. Keuroghlian, D. P. Eaton, C. L. Neves, M. S. Nardi, C. Braga, P. R. Gonçalves, A. C. Srbek-Araujo, P. Mendes, J. A. de Oliveira, F. A. M. Soares, P. A. Rocha, P. Crawshaw, M. C. Ribeiro, M. Galetti, Atlantic Mammal Traits: a data set of morphological traits of mammals in the Atlantic Forest of South America. *Ecology*. **99**, 498 (2018). |
| Chiroptera | Natalidae | Mexican funnel-eared bat | *Natalus stramineus* | 40 | 23 | J. B. Akins, M. L. Kennedy, G. D. Schnell, C. Sánchez-Hernández, M. de L. Romero-Almaraz, M. C. Wooten, T. L. Best, Flight speeds of three species of Neotropical bats: *Glossophaga soricina*, *Natalus stramineus*, and *Carollia subrufa*. *Acta Chiropterologica*. **9**, 477–482 (2007). |

| | | | | | | |
|---|---|---|---|---|---|---|
| Chiroptera | Phyllostomidae | Anoura peruana | *Anoura peruana* | 5 | 3 | D. Ocampo, K. G. Borja-Acosta, J. Lozano-Flórez, S. Cifuentes-Acevedo, E. Arbeláez-Cortés, N. J. Bayly, Á. Caguazango, B. Coral-Jaramillo, D. Cueva, F. Forero, J. P. Gómez, C. Gómez, M. A. Loaiza-Muñoz, G. A. Londoño, S. Losada-Prado, S. Pérez-Peña, H. E. Ramírez-Chaves, M. E. Rodríguez-Posada, J. Sanabria-Mejía, M. Sánchez-Martínez, V. H. Serrano-Cardozo, M. S. Sierra-Buitrago, J. Soto-Patiño, O. Acevedo-Charry, Body mass data set for 1,317 bird and 270 mammal species from Colombia. *Ecology*. **102** (2021), doi:10.1002/ecy.3273. |
| Chiroptera | Phyllostomidae | Artibeus aequatorialis | *Artibeus aequatorialis* | 8 | 4 | D. Ocampo, K. G. Borja-Acosta, J. Lozano-Flórez, S. Cifuentes-Acevedo, E. Arbeláez-Cortés, N. J. Bayly, Á. Caguazango, B. Coral-Jaramillo, D. Cueva, F. Forero, J. P. Gómez, C. Gómez, M. A. Loaiza-Muñoz, G. A. Londoño, S. Losada-Prado, S. Pérez-Peña, H. E. Ramírez-Chaves, M. E. Rodríguez-Posada, J. Sanabria-Mejía, M. Sánchez-Martínez, V. H. Serrano-Cardozo, M. S. Sierra-Buitrago, J. Soto-Patiño, O. Acevedo-Charry, Body mass data set for 1,317 bird and 270 mammal species from Colombia. *Ecology*. **102** (2021), doi:10.1002/ecy.3273. |
| Chiroptera | Phyllostomidae | Artibeus amplus | *Artibeus amplus* | 9 | 6 | D. Ocampo, K. G. Borja-Acosta, J. Lozano-Flórez, S. Cifuentes-Acevedo, E. Arbeláez-Cortés, N. J. Bayly, Á. Caguazango, B. Coral-Jaramillo, D. Cueva, F. Forero, J. P. Gómez, C. Gómez, M. A. Loaiza-Muñoz, G. A. Londoño, S. Losada-Prado, S. Pérez-Peña, H. E. Ramírez-Chaves, M. E. Rodríguez-Posada, J. Sanabria-Mejía, M. Sánchez-Martínez, V. H. Serrano-Cardozo, M. S. Sierra-Buitrago, J. Soto-Patiño, O. Acevedo-Charry, Body mass data set for 1,317 bird and 270 mammal species from Colombia. *Ecology*. **102** (2021), doi:10.1002/ecy.3273. |

| | | | | | | |
|---|---|---|---|---|---|---|
| Chiroptera | Phyllostomidae | Carollia perspicillata | *Carollia perspicillata* | 190 | 161 | D. Ocampo, K. G. Borja-Acosta, J. Lozano-Flórez, S. Cifuentes-Acevedo, E. Arbeláez-Cortés, N. J. Bayly, Á. Caguazango, B. Coral-Jaramillo, D. Cueva, F. Forero, J. P. Gómez, C. Gómez, M. A. Loaiza-Muñoz, G. A. Londoño, S. Losada-Prado, S. Pérez-Peña, H. E. Ramírez-Chaves, M. E. Rodríguez-Posada, J. Sanabria-Mejía, M. Sánchez-Martínez, V. H. Serrano-Cardozo, M. S. Sierra-Buitrago, J. Soto-Patiño, O. Acevedo-Charry, Body mass data set for 1,317 bird and 270 mammal species from Colombia. *Ecology*. **102** (2021), doi:10.1002/ecy.3273. |
| Chiroptera | Phyllostomidae | Gray short-tailed bat | *Carollia subrufa* | 3 | 3 | J. B. Akins, M. L. Kennedy, G. D. Schnell, C. Sánchez-Hernández, M. de L. Romero-Almaraz, M. C. Wooten, T. L. Best, Flight speeds of three species of Neotropical bats: *Glossophaga soricina*, *Natalus stramineus*, and *Carollia subrufa*. *Acta Chiropterologica*. **9**, 477–482 (2007). |

| | | | | | | |
|---|---|---|---|---|---|---|
| Chiroptera | Phyllostomidae | Pygmy round-eared bat | *Lophostoma brasiliense* | 15 | 16 | F. Gonçalves, R. S. Bovendorp, G. Beca, C. Bello, R. Costa-Pereira, R. L. Muylaert, R. R. Rodarte, N. Villar, R. Souza, M. E. Graipel, J. J. Cherem, D. Faria, J. Baumgarten, M. R. Alvarez, E. M. Vieira, N. Cáceres, R. Pardini, Y. L. R. Leite, L. P. Costa, M. A. R. Mello, E. Fischer, F. C. Passos, L. H. Varzinczak, J. A. Prevedello, A. P. Cruz-Neto, F. Carvalho, A. R. Percequillo, A. Paviolo, A. Nava, J. M. B. Duarte, N. U. de la Sancha, E. Bernard, R. G. Morato, J. F. Ribeiro, R. G. Becker, G. Paise, P. S. Tomasi, F. Vélez-Garcia, G. L. Melo, J. Sponchiado, F. Cerezer, M. A. S. Barros, A. Q. S. de Souza, C. C. dos Santos, G. A. F. Giné, P. Kerches-Rogeri, M. M. Weber, G. Ambar, L. V. Cabrera-Martinez, A. Eriksson, M. Silveira, C. F. Santos, L. Alves, E. Barbier, G. C. Rezende, G. S. T. Garbino, É. O. Rios, A. Silva, A. T. A. Nascimento, R. S. de Carvalho, A. Feijó, J. Arrabal, I. Agostini, D. Lamattina, S. Costa, E. Vanderhoeven, F. R. de Melo, P. de Oliveira Laroque, L. Jerusalinsky, M. M. Valença-Montenegro, A. B. Martins, G. Ludwig, R. B. de Azevedo, A. Anzóategui, M. X. da Silva, M. Figuerêdo Duarte Moraes, A. Vogliotti, A. Gatti, T. Püttker, C. S. Barros, T. K. Martins, A. Keuroghlian, D. P. Eaton, C. L. Neves, M. S. Nardi, C. Braga, P. R. Gonçalves, A. C. Srbek-Araujo, P. Mendes, J. A. de Oliveira, F. A. M. Soares, P. A. Rocha, P. Crawshaw, M. C. Ribeiro, M. Galetti, Atlantic Mammal Traits: a data set of morphological traits of mammals in the Atlantic Forest of South America. *Ecology*. **99**, 498 (2018). |
| Chiroptera | Phyllostomidae | White-throated round-eared bat | *Lophostoma silvicolum* | 136 | 138 | D. K. N. Dechmann, E. K. V. Kalko, B. König, G. Kerth, Mating system of a neotropical roost-making bat: The white-throated, round-eared bat, *Lophostoma silvicolum* (Chiroptera: Phyllostomidae). *Behav. Ecol. Sociobiol.* **58**, 316–325 (2005). |

| | | | | | | |
|---|---|---|---|---|---|---|
| Chiroptera | Pteropodidae | Spotted-winged fruit bat | *Balionycteris maculata* | 25 | 54 | R. Hodgkison, S. T. Balding, Z. Akbar, T. H. Kunz, Roosting ecology and social organization of the spotted-winged fruit bat, *Balionycteris maculata* (Chiroptera: Pteropodidae), in a Malaysian lowland dipterocarp forest. *J. Trop. Ecol.* **19**, 667–676 (2003). |
| Chiroptera | Pteropodidae | Lesser short-nosed fruit bat | *Cynopterus brachyotis* | 23 | 40 | K. Funakoshi, Z. Akbar, Behavioural and reproductive ecology of the dog-faced fruit bats, *Cynopterus brachyotis* and *C. horsfieldi*, in a Malaysian rainforest. *Mammal Study*. **22** (1997), pp. 95–108. |
| Chiroptera | Pteropodidae | Horsfield's fruit bat | *Cynopterus horsfieldi* | 19 | 26 | K. Funakoshi, Z. Akbar, Behavioural and reproductive ecology of the dog-faced fruit bats, *Cynopterus brachyotis* and *C. horsfieldi*, in a Malaysian rainforest. *Mammal Study*. **22** (1997), pp. 95–108. |
| Chiroptera | Pteropodidae | Greater short-nosed fruit bat | *Cynopterus sphinx* | 30 | 30 | V. Nagarajan-Radha, P. S. D. Devaraj, Sex differences in postprandial blood glucose and body surface temperature are contingent on flight in the fruit bat, *Cynopterus sphinx*. *Biol. Open*. **10** (2021), doi:10.1242/bio.053926. |
| Chiroptera | Pteropodidae | Cave nectar bat | *Eonycteris spelaea* | 7 | 5 | P. R. Acharya, P. A. Racey, S. Sotthibandhu, S. Bumrungsri, Home-range and foraging areas of the dawn bat *Eonycteris spelaea* in agricultural areas of Thailand. *Acta Chiropterologica*. **17**, 307–319 (2015). |
| Chiroptera | Pteropodidae | Fischer's pygmy fruit bat | *Haplonycteris fischeri* | 11 | 10 | L. R. Heaney, B. R. Tabaranza, E. A. Rickart, D. S. Balete, N. R. Ingle, The mammals of Mt. Kitanglad Nature Park, Mindanao, Philippines. *Fieldiana Zool.* **112**, 1–63 (2006). |
| Chiroptera | Pteropodidae | Harpy fruit bat | *Harpyionycteris whiteheadi* | 9 | 2 | L. R. Heaney, B. R. Tabaranza, E. A. Rickart, D. S. Balete, N. R. Ingle, The mammals of Mt. Kitanglad Nature Park, Mindanao, Philippines. *Fieldiana Zool.* **112**, 1–63 (2006). |

| | | | | | | |
|---|---|---|---|---|---|---|
| Chiroptera | Pteropodidae | Salim Ali's fruit bat | *Latidens salimalii* | 16 | 19 | G. Agoramoorthy, M. J. Hsu, Population size, feeding, forearm length and body weight of a less known Indian fruit bat, *Latidens salimalii*. *Curr. Sci.* **88**, 354–356 (2005). |
| Chiroptera | Pteropodidae | Eastern tube-nosed bat | *Nyctimene robinsoni* | 3 | 4 | A. Riek, G. Körtner, F. Geiser, Thermobiology, energetics and activity patterns of the Eastern tube-nosed bat (*Nyctimene robinsoni*) in the Australian tropics: Effect of temperature and lunar cycle. *J. Exp. Biol.* **213**, 2557–2564 (2010). |
| Chiroptera | Pteropodidae | Christmas Island flying fox | *Pteropus natalis* | 15 | 8 | C. M. Todd, D. A. Westcott, J. M. Martin, K. Rose, A. McKeown, J. Hall, J. A. Welbergen, Body-size dependent foraging strategies in the Christmas Island flying-fox: implications for seed and pollen dispersal within a threatened island ecosystem. *Mov. Ecol.* **10**, 1–15 (2022). |
| Chiroptera | Pteropodidae | Mauritian flying fox | *Pteropus niger* | 6 | 6 | R. Z. Oleksy, C. L. Ayady, V. Tatayah, C. Jones, P. W. Howey, J. S. P. Froidevaux, P. A. Racey, G. Jones, The movement ecology of the Mauritian flying fox (*Pteropus niger*): A long-term study using solar-powered GSM/GPS tags. *Mov. Ecol.* **7**, 1–12 (2019). |
| Chiroptera | Pteropodidae | Gray-headed flying fox | *Pteropus poliocephalus* | 87 | 73 | J. A. Welbergen, Growth, bimaturation, and sexual size dimorphism in wild gray-headed flying foxes (*Pteropus poliocephalus*). *J. Mammal.* **91**, 38–47 (2010). |
| Chiroptera | Pteropodidae | Bonin flying fox | *Pteropus pselaphon* | 3 | 2 | N. Sugita, M. Inaba, K. Ueda, Roosting pattern and reproductive cycle of Bonin flying foxes (*Pteropus pselaphon*). *J. Mammal.* **90**, 195–202 (2009). |
| Chiroptera | Pteropodidae | Madagascan flying fox | *Pteropus rufus* | 67 | 82 | H. C. Ranaivoson, J. M. Héraud, H. K. Goethert, S. R. Telford, L. Rabetafika, C. E. Brook, Babesial infection in the Madagascan flying fox, *Pteropus rufus* É. Geoffroy, 1803. *Parasites and Vectors*. **12**, 1–13 (2019). |

| | | | | | | |
|---|---|---|---|---|---|---|
| Chiroptera | Pteropodidae | Madagascan rousette | *Rousettus madagascariensis* | 271 | 289 | A. Riek, G. Körtner, F. Geiser, Thermobiology, energetics and activity patterns of the Eastern tube-nosed bat (*Nyctimene robinsoni*) in the Australian tropics: Effect of temperature and lunar cycle. *J. Exp. Biol.* **213**, 2557–2564 (2010). |
| Chiroptera | Rhinolophidae | Cape horseshoe bat | *Rhinolophus capensis* | 23 | 23 | L. J. Odendaal, D. S. Jacobs, Morphological correlates of echolocation frequency in the endemic Cape horseshoe bat, *Rhinolophus capensis* (Chiroptera: Rhinolophidae). *J. Comp. Physiol. A Neuroethol. Sensory, Neural, Behav. Physiol.* **197**, 435–446 (2011). |
| Chiroptera | Rhinolophidae | Mediterranean horseshoe bat | *Rhinolophus euryale* | 9 | 23 | D. Russo, G. Jones, M. Mucedda, Influence of age, sex and body size on echolocation calls of Mediterranean and Mehely's horseshoe bats, *Rhinolophus euryale* and *R. mehelyi* (Chiroptera: Rhinolophidae). *Mammalia*. **65**, 429–436 (2001). |
| Chiroptera | Rhinolophidae | Greater horseshoe bat | *Rhinolophus ferrumequinum* | 31 | 77 | T. Jiang, J. Wang, H. Wu, G. Csorba, S. J. Puechmaille, P. Benda, J. Boireau, R. Toffoli, J. Y. Courtois, P. Nyssen, R. Colombo, J. Feng, The patterns and possible causes of global geographical variation in the body size of the greater horseshoe bat (*Rhinolophus ferrumequinum*). *J. Biogeogr.* **46**, 2363–2377 (2019). |
| Chiroptera | Rhinolophidae | Lesser horseshoe bat | *Rhinolophus hipposideros* | 19 | 14 | J. Kohyt, E. Pierzchała, A. Koza, K. Piksa, Resting frequency of echolocation calls within a lesser horseshoe bat population (southern Poland) and its relation to body size, condition and mass. *Folia Biol.* **70**, 33–42 (2022). |
| Chiroptera | Rhinolophidae | Philippine forest horseshoe bat | *Rhinolophus inops* | 12 | 4 | H. de G. Bergallo, Ecology of a small mammal community in an Atlantic Forest Area in Southeastern Brazil. *Stud. Neotrop. Fauna Environ.* **29**, 197–217 (1994). |

| | | | | | | |
|---|---|---|---|---|---|---|
| Chiroptera | Rhinolophidae | Mehely's horseshoe bat | *Rhinolophus mehelyi* | 10 | 18 | D. Russo, G. Jones, M. Mucedda, Influence of age, sex and body size on echolocation calls of Mediterranean and Mehely's horseshoe bats, *Rhinolophus euryale* and *R. mehelyi* (Chiroptera: Rhinolophidae). *Mammalia*. **65**, 429–436 (2001). |
| Chiroptera | Vespertilionidae | Western barbastelle | *Barbastella barbastellus* | 21 | 6 | R. Toffoli, P. Culasso, A. G. Locatelli, L. Giraudo, Bats of Alpi Marittime Nature Park (north west Italy) and site of community importance IT1160056: Distribution and status. *Nat. Hist. Sci.* **3**, 3–13 (2016). |
| Chiroptera | Vespertilionidae | Gould's wattled bat | *Chalinolobus gouldii* | 365 | 201 | B. Chruszcz, R. M. Barclay, *Chalinolobus gouldii*. *Mamm. Species*. **690**, 1–4 (2002). |

| | | | | | | |
|---|---|---|---|---|---|---|
| Chiroptera | Vespertilionidae | Serotine bat | *Eptesicus serotinus* | 33 | 6 | Jaap van Schaik, pers. comm. |
| Chiroptera | Vespertilionidae | Pied Butterfly Bat | *Glauconycteris superba* | 6 | 3 | R. Kiri, R. Colombo, G. C. Gembu, Y. Bas, J. F. Julien, Y. Gager, A. Hassanin, Echolocation calls and flight behaviour of the elusive pied butterfly bat (*Glauconycteris superba*), and new data on its morphology and ecology. *Acta Chiropterologica*. **18**, 477–488 (2016). |
| Chiroptera | Vespertilionidae | Savi's pipistrelle | *Hypsugo savii* | 71 | 38 | M. Rughetti, R. Toffoli, Sex-specific seasonal change in body mass in two species of vespertilionid bats. *Acta Chiropterologica*. **16**, 149–155 (2014). |
| Chiroptera | Vespertilionidae | Dark woolly bat | *Kerivoula furva* | 81 | 68 | H. C. Kuo, P. Soisook, Y. Y. Ho, G. Csorba, C. N. Wang, S. J. Rossiter, A taxonomic revision of the Kerivoula hardwickii complex (Chiroptera: Vespertilionidae) with the description of a new species. *Acta Chiropterologica*. **19**, 19–39 (2017). |

| | | | | | | |
|---|---|---|---|---|---|---|
| Chiroptera | Vespertilionidae | Southern red bat | *Lasiurus blossevillii* | 15 | 9 | F. Gonçalves, R. S. Bovendorp, G. Beca, C. Bello, R. Costa-Pereira, R. L. Muylaert, R. R. Rodarte, N. Villar, R. Souza, M. E. Graipel, J. J. Cherem, D. Faria, J. Baumgarten, M. R. Alvarez, E. M. Vieira, N. Cáceres, R. Pardini, Y. L. R. Leite, L. P. Costa, M. A. R. Mello, E. Fischer, F. C. Passos, L. H. Varzinczak, J. A. Prevedello, A. P. Cruz-Neto, F. Carvalho, A. R. Percequillo, A. Paviolo, A. Nava, J. M. B. Duarte, N. U. de la Sancha, E. Bernard, R. G. Morato, J. F. Ribeiro, R. G. Becker, G. Paise, P. S. Tomasi, F. Vélez-Garcia, G. L. Melo, J. Sponchiado, F. Cerezer, M. A. S. Barros, A. Q. S. de Souza, C. C. dos Santos, G. A. F. Giné, P. Kerches-Rogeri, M. M. Weber, G. Ambar, L. V. Cabrera-Martinez, A. Eriksson, M. Silveira, C. F. Santos, L. Alves, E. Barbier, G. C. Rezende, G. S. T. Garbino, É. O. Rios, A. Silva, A. T. A. Nascimento, R. S. de Carvalho, A. Feijó, J. Arrabal, I. Agostini, D. Lamattina, S. Costa, E. Vanderhoeven, F. R. de Melo, P. de Oliveira Laroque, L. Jerusalinsky, M. M. Valença-Montenegro, A. B. Martins, G. Ludwig, R. B. de Azevedo, A. Anzóategui, M. X. da Silva, M. Figuerêdo Duarte Moraes, A. Vogliotti, A. Gatti, T. Püttker, C. S. Barros, T. K. Martins, A. Keuroghlian, D. P. Eaton, C. L. Neves, M. S. Nardi, C. Braga, P. R. Gonçalves, A. C. Srbek-Araujo, P. Mendes, J. A. de Oliveira, F. A. M. Soares, P. A. Rocha, P. Crawshaw, M. C. Ribeiro, M. Galetti, Atlantic Mammal Traits: a data set of morphological traits of mammals in the Atlantic Forest of South America. *Ecology*. **99**, 498 (2018). |
| Chiroptera | Vespertilionidae | Chilean red bat | *Lasiurus varius* | 23 | 4 | G. Ossa, M. M. Díaz, R. M. Barquez, *Lasiurus varius* (Chiroptera: Vespertilionidae). *Mamm. Species*. **51**, 119–127 (2019). |

| | | | | | | |
|---|---|---|---|---|---|---|
| Chiroptera | Vespertilionidae | Fea's tube-nosed bat | *Murina feae* | 3 | 2 | P. Soisook, thesis, Prince of Songkla University (2013). |
| Chiroptera | Vespertilionidae | Long-eared Myotis | *Murina guilleni guilleni* | 6 | 3 | P. Soisook, S. Karapan, C. Satasook, V. D. Thong, F. A. A. Khan, I. Maryanto, G. Csorba, N. Furey, B. Aul, P. J. J. Bates, A Review of the *Murina cyclotis* complex (Chiroptera: Vespertilionidae) with descriptions of a new species and subspecies. *Acta Chiropterologica*. **15**, 271–292 (2013). |
| Chiroptera | Vespertilionidae | Murina peninsularis | *Murina peninsularis* | 23 | 19 | P. Soisook, S. Karapan, C. Satasook, V. D. Thong, F. A. A. Khan, I. Maryanto, G. Csorba, N. Furey, B. Aul, P. J. J. Bates, A Review of the *Murina cyclotis* complex (Chiroptera: Vespertilionidae) with descriptions of a new species and subspecies. *Acta Chiropterologica*. **15**, 271–292 (2013). |
| Chiroptera | Vespertilionidae | Bechstein's bat | *Myotis bechsteinii* | 37 | 8 | Jaap van Schaik, pers. comm. |
| Chiroptera | Vespertilionidae | Brandt's myotis | *Myotis brandtii* | 31 | 68 | R. Lucan, P. Benda, A. Reiter, J. Zima, Reliability of field determination in three cryptic whiskered bats (*Myotis alcathoe , M. mystacinus , M. brandtii*) and basic biometric characters: evidence from the Czech Republic. *Vespertilio*. **15**, 55–62 (2011). |

| | | | | | | |
|---|---|---|---|---|---|---|
| Chiroptera | Vespertilionidae | Myotis caucensis | *Myotis caucensis* | 3 | 2 | D. Ocampo, K. G. Borja-Acosta, J. Lozano-Flórez, S. Cifuentes-Acevedo, E. Arbeláez-Cortés, N. J. Bayly, Á. Caguazango, B. Coral-Jaramillo, D. Cueva, F. Forero, J. P. Gómez, C. Gómez, M. A. Loaiza-Muñoz, G. A. Londoño, S. Losada-Prado, S. Pérez-Peña, H. E. Ramírez-Chaves, M. E. Rodríguez-Posada, J. Sanabria-Mejía, M. Sánchez-Martínez, V. H. Serrano-Cardozo, M. S. Sierra-Buitrago, J. Soto-Patiño, O. Acevedo-Charry, Body mass data set for 1,317 bird and 270 mammal species from Colombia. *Ecology*. **102** (2021), doi:10.1002/ecy.3273. |
| Chiroptera | Vespertilionidae | Chilean myotis | *Myotis chiloensis* | 23 | 3 | G. Ossa, M. M. Díaz, R. M. Barquez, *Lasiurus varius* (Chiroptera: Vespertilionidae). *Mamm. Species*. **51**, 119–127 (2019). |
| Chiroptera | Vespertilionidae | Pond bat | *Myotis dasycneme* | 28 | 12 | Jaap van Schaik, pers. comm. |
| Chiroptera | Vespertilionidae | Daubenton's bat | *Myotis daubentonii* | 291 | 92 | Jaap van Schaik, pers. comm. |
| Chiroptera | Vespertilionidae | Geoffroy's bat | *Myotis emarginatus* | 159 | 34 | Jaap van Schaik, pers. comm. |
| Chiroptera | Vespertilionidae | | *Myotis evosi* | 37 | 32 | D. I. Solick, R. M. R. Barclay, Morphological differences among western long-eared myotis (*Myotis evotis*) populations in different environments. *J. Mammal.* **87**, 1020–1026 (2006). |

| | | | | | | |
|---|---|---|---|---|---|---|
| Chiroptera | Vespertilionidae | LaVal's myotis | *Myotis lavali* | 384 | 13 | F. Gonçalves, R. S. Bovendorp, G. Beca, C. Bello, R. Costa-Pereira, R. L. Muylaert, R. R. Rodarte, N. Villar, R. Souza, M. E. Graipel, J. J. Cherem, D. Faria, J. Baumgarten, M. R. Alvarez, E. M. Vieira, N. Cáceres, R. Pardini, Y. L. R. Leite, L. P. Costa, M. A. R. Mello, E. Fischer, F. C. Passos, L. H. Varzinczak, J. A. Prevedello, A. P. Cruz-Neto, F. Carvalho, A. R. Percequillo, A. Paviolo, A. Nava, J. M. B. Duarte, N. U. de la Sancha, E. Bernard, R. G. Morato, J. F. Ribeiro, R. G. Becker, G. Paise, P. S. Tomasi, F. Vélez-Garcia, G. L. Melo, J. Sponchiado, F. Cerezer, M. A. S. Barros, A. Q. S. de Souza, C. C. dos Santos, G. A. F. Giné, P. Kerches-Rogeri, M. M. Weber, G. Ambar, L. V. Cabrera-Martinez, A. Eriksson, M. Silveira, C. F. Santos, L. Alves, E. Barbier, G. C. Rezende, G. S. T. Garbino, É. O. Rios, A. Silva, A. T. A. Nascimento, R. S. de Carvalho, A. Feijó, J. Arrabal, I. Agostini, D. Lamattina, S. Costa, E. Vanderhoeven, F. R. de Melo, P. de Oliveira Laroque, L. Jerusalinsky, M. M. Valença-Montenegro, A. B. Martins, G. Ludwig, R. B. de Azevedo, A. Anzóategui, M. X. da Silva, M. Figuerêdo Duarte Moraes, A. Vogliotti, A. Gatti, T. Püttker, C. S. Barros, T. K. Martins, A. Keuroghlian, D. P. Eaton, C. L. Neves, M. S. Nardi, C. Braga, P. R. Gonçalves, A. C. Srbek-Araujo, P. Mendes, J. A. de Oliveira, F. A. M. Soares, P. A. Rocha, P. Crawshaw, M. C. Ribeiro, M. Galetti, Atlantic Mammal Traits: a data set of morphological traits of mammals in the Atlantic Forest of South America. *Ecology*. **99**, 498 (2018). |
| Chiroptera | Vespertilionidae | Yellowish myotis | *Myotis levis* | 50 | 24 | R. A. Araújo, B. D. Amaro, S. A. Talamoni, H. P. Godinho, Seasonal reproduction of yellowish myotis, *Myotis levis* (Chiroptera: Vespertilionidae), from a Neotropical highland. *J. Morphol.* **274**, 1230–1238 (2013). |

| | | | | | | |
|---|---|---|---|---|---|---|
| Chiroptera | Vespertilionidae | Greater mouse-eared bat | *Myotis myotis* | 9 | 12 | Jaap van Schaik, pers. comm. |
| Chiroptera | Vespertilionidae | Whiskered bat | *Myotis mystacinus* | 26 | 91 | R. Lucan, P. Benda, A. Reiter, J. Zima, Reliability of field determination in three cryptic whiskered bats (*Myotis alcathoe , M. mystacinus , M. brandtii*) and basic biometric characters: evidence from the Czech Republic. *Vespertilio*. **15**, 55–62 (2011). |
| Chiroptera | Vespertilionidae | Natterer's bat | *Myotis nattereri* | 200 | 97 | Jaap van Schaik, pers. comm. |

| | | | | | | |
|---|---|---|---|---|---|---|
| Chiroptera | Vespertilionidae | Cave Myotis | *Myotis velifer* | 148 | 212 | W. Caire, L. S. Loucks, Loss in mass by hibernating cave myotis, *Myotis velifer* (Chiroptera: Vespertilionidae) in western Oklahoma. *Southwest. Nat.* **55**, 323–330 (2010). |
| Chiroptera | Vespertilionidae | Fish-eating myotis | *Myotis vivesi* | 31 | 18 | A. Otálora-Ardila, J. J. Flores-Martínez, C. Rosales, A. Salame-Méndez, L. G. H. Montalvo, Physiological and ecological correlates of the cellular and humoral innate immune responses in an insular desert bat: the fish-eating myotis (*Myotis vivesi*). *Diversity*. **14** (2022), doi:10.3390/d14100781. |
| Chiroptera | Vespertilionidae | Lesser noctule | *Nyctalus leisleri* | 5 | 5 | R. Toffoli, P. Culasso, A. G. Locatelli, L. Giraudo, Bats of Alpi Marittime Nature Park (north west Italy) and site of community importance IT1160056: Distribution and status. *Nat. Hist. Sci.* **3**, 3–13 (2016). |
| Chiroptera | Vespertilionidae | Common noctule | *Nyctalus nyctalus* | 88 | 357 | M. T. O'Mara, K. Bauer, D. Blank, J. W. Baldwin, K. Dina, N. Dechmann, Common noctule bats are sexually dimorphic in migratory behaviour and body size but not wing shape. *PLoS One*. **11**, e0167027 (2016). |
| Chiroptera | Vespertilionidae | Kuhl's pipistrelle | *Pipistrellus kuhlii* | 5 | 3 | B. M. McAllan, J. R. Roberts, T. O'Shea, Seasonal changes in glomerular filtration rate in *Antechinus stuartii* (Marsupialia: Dasyuridae). *J. Comp. Physiol. - B Biochem. Syst. Environ. Physiol.* **168**, 41–49 (1998). |
| Chiroptera | Vespertilionidae | Common pipistrelle | *Pipistrellus pipistrellus* | 50 | 61 | Jaap van Schaik, pers. comm. |
| Chiroptera | Vespertilionidae | Brown long-eared bat | *Plecotus auritus* | 43 | 10 | Jaap van Schaik, pers. comm. |

| | | | | | | |
|---|---|---|---|---|---|---|
| Cingulata | Chlamyphoridae | Squealing Hairy Armadillo | *Chaetophractus vellerosus* | 2 | 3 | V. P. Seitz, S. Puig, Aboveground activity, reproduction, body temperature and weight of armadillos (Xenarthra, Chlamyphoridae) according to atmospheric conditions in the central Monte (Argentina). *Mamm. Biol.* **88**, 43–51 (2018). |
| Cingulata | Chlamyphoridae | Big hairy armadillo | *Chaetophractus villosus* | 4 | 7 | V. P. Seitz, S. Puig, Aboveground activity, reproduction, body temperature and weight of armadillos (Xenarthra, Chlamyphoridae) according to atmospheric conditions in the central Monte (Argentina). *Mamm. Biol.* **88**, 43–51 (2018). |
| Cingulata | Chlamyphoridae | Giant armadillo | *Priodontes maximus* | 9 | 9 | A. L. J. Desbiez, G. F. Massocato, D. Kluyber, C. D. N. Luba, N. Attias, How giant are giant armadillos? The morphometry of giant armadillos (*Priodontes maximus* Kerr, 1792) in the Pantanal of Brazil. *Mamm. Biol.* **95**, 9–14 (2019). |
| Cingulata | Chlamyphoridae | Pichi | *Zaedyus pichiy* | 30 | 21 | V. P. Seitz, S. Puig, Aboveground activity, reproduction, body temperature and weight of armadillos (Xenarthra, Chlamyphoridae) according to atmospheric conditions in the central Monte (Argentina). *Mamm. Biol.* **88**, 43–51 (2018). |
| Cingulata | Dasypodidae | Nine-banded armadillo | *Dasypus novemcinctus* | 11 | 13 | R. G. Tague, Pelvic sexual dimorphism among species monomorphic in body size: Relationship to relative newborn body mass. *J. Mammal.* **97**, 503–517 (2016). |
| Dasyuromorphia | Dasyuridae | Brown antechinus | *Antechinus stuartii* | 17 | 10 | B. M. McAllan, J. R. Roberts, T. O’Shea, Seasonal changes in glomerular filtration rate in *Antechinus stuartii* (Marsupialia: Dasyuridae). *J. Comp. Physiol. - B Biochem. Syst. Environ. Physiol.* **168**, 41–49 (1998). |
| Dasyuromorphia | Dasyuridae | Northern quoll | *Dasyurus hallucatus* | 20 | 16 | L. Hernandez-Santin, J. A. Dunlop, A. W. Goldizen, D. O. Fisher, Demography of the northern quoll (*Dasyurus hallucatus*) in the most arid part of its range. *J. Mammal.* **100**, 1191–1198 (2019). |

| | | | | | | |
|---|---|---|---|---|---|---|
| Dasyuromo rphia | Dasyuridae | Tiger quoll | *Dasyurus maculatus* | 41 | 20 | A. S. Glen, Population attributes of the spotted-tailed quoll (*Dasyurus maculatus*) in north-eastern New South Wales. *Aust. J. Zool.* **56**, 137–142 (2008). |
| Dasyuromo rphia | Dasyuridae | Red-tailed phascogale | *Phascogale calura* | 15 | 56 | A. J. Bradley, Reproduction and life history in the red-tailed phascogale, *Phascogale calura* (Marsupialia: Dasyuridae): The adaptive-stress senescence hypothesis. *J. Zool.* **241**, 739–755 (1997). |
| Dasyuromo rphia | Dasyuridae | Tasmanian devil | *Sarcophilus harrisii* | 35 3 | 51 3 | S. Belkhir, R. Hamede, F. Thomas, B. Ujvari, A. M. Dujon, Season, weight, and age, but not transmissible cancer, affect tick loads in the endangered Tasmanian devil. *Infect. Genet. Evol.* **98** (2022), doi:10.1016/j.meegid.2022.105221. |

| | | | | | | |
|---|---|---|---|---|---|---|
| Didelphimorphia | Didelphidae | Southeastern four-eyed opossum | *Philander frenatus* | 18 | 30 | F. Gonçalves, R. S. Bovendorp, G. Beca, C. Bello, R. Costa-Pereira, R. L. Muylaert, R. R. Rodarte, N. Villar, R. Souza, M. E. Graipel, J. J. Cherem, D. Faria, J. Baumgarten, M. R. Alvarez, E. M. Vieira, N. Cáceres, R. Pardini, Y. L. R. Leite, L. P. Costa, M. A. R. Mello, E. Fischer, F. C. Passos, L. H. Varzinczak, J. A. Prevedello, A. P. Cruz-Neto, F. Carvalho, A. R. Percequillo, A. Paviolo, A. Nava, J. M. B. Duarte, N. U. de la Sancha, E. Bernard, R. G. Morato, J. F. Ribeiro, R. G. Becker, G. Paise, P. S. Tomasi, F. Vélez-Garcia, G. L. Melo, J. Sponchiado, F. Cerezer, M. A. S. Barros, A. Q. S. de Souza, C. C. dos Santos, G. A. F. Giné, P. Kerches-Rogeri, M. M. Weber, G. Ambar, L. V. Cabrera-Martinez, A. Eriksson, M. Silveira, C. F. Santos, L. Alves, E. Barbier, G. C. Rezende, G. S. T. Garbino, É. O. Rios, A. Silva, A. T. A. Nascimento, R. S. de Carvalho, A. Feijó, J. Arrabal, I. Agostini, D. Lamattina, S. Costa, E. Vanderhoeven, F. R. de Melo, P. de Oliveira Laroque, L. Jerusalinsky, M. M. Valença-Montenegro, A. B. Martins, G. Ludwig, R. B. de Azevedo, A. Anzóategui, M. X. da Silva, M. Figuerêdo Duarte Moraes, A. Vogliotti, A. Gatti, T. Püttker, C. S. Barros, T. K. Martins, A. Keuroghlian, D. P. Eaton, C. L. Neves, M. S. Nardi, C. Braga, P. R. Gonçalves, A. C. Srbek-Araujo, P. Mendes, J. A. de Oliveira, F. A. M. Soares, P. A. Rocha, P. Crawshaw, M. C. Ribeiro, M. Galetti, Atlantic Mammal Traits: a data set of morphological traits of mammals in the Atlantic Forest of South America. *Ecology*. **99**, 498 (2018). |
| Diprotodontia | Burramyidae | Cercartetus nanus | *Eastern pygmy possum* | 33 | 32 | J. M. Harris, R. L. Goldingay, L. O. Brooks, Population ecology of the eastern pygmy-possum (*Cercartetus nanus*) in a montane woodland in southern New South Wales. *Aust. Mammal.* **36**, 212–218 (2014). |

| | | | | | | |
|---|---|---|---|---|---|---|
| Diprotodontia | Macropodidae | Lumholt's tree-kangaroo | *Dendrolagus lumholtzi* | 12 | 7 | G. R. Newell, Home range and habitat use by Lumholtz's tree-kangaroo (*Dendrolagus lumholtzi*) within a rainforest fragment in north Queensland. *Wildl. Res.* **26**, 129–145 (1999). |
| Diprotodontia | Macropodidae | Rufous hare-wallaby | *Lagorchestes hirsutus* | 59 | 64 | J. D. Richards, J. Short, R. I. T. Prince, J. A. Friend, J. M. Courtenay, The biology of banded (*Lagostrophus fasciatus*) and rufous (*Lagorchestes hirsutus*) hare-wallabies (Diprotodontia: Macropodidae) on Dorre and Bernier Islands, Western Australia. *Wildl. Res.* **28**, 311–322 (2001). |
| Diprotodontia | Macropodidae | Banded hare-wallaby | *Lagostrophus fasciatus* | 77 | 105 | J. D. Richards, J. Short, R. I. T. Prince, J. A. Friend, J. M. Courtenay, The biology of banded (*Lagostrophus fasciatus*) and rufous (*Lagorchestes hirsutus*) hare-wallabies (Diprotodontia: Macropodidae) on Dorre and Bernier Islands, Western Australia. *Wildl. Res.* **28**, 311–322 (2001). |
| Diprotodontia | Macropodidae | Tammar wallaby | *Macropus eugenii* | 34 | 35 | A. C. Taylor, P. Sunnucks, D. W. Cooper, Retention of reproductive barriers and ecological differences between two introduced sympatric *Macropus* spp. in New Zealand. *Anim. Conserv.* **2**, 195–202 (1999). |
| Diprotodontia | Macropodidae | Parma wallaby | *Macropus parma* | 21 | 14 | A. C. Taylor, P. Sunnucks, D. W. Cooper, Retention of reproductive barriers and ecological differences between two introduced sympatric *Macropus* spp. in New Zealand. *Anim. Conserv.* **2**, 195–202 (1999). |
| Diprotodontia | Macropodidae | Red-necked wallaby | *Macropus rufogriseus rufogriseus* | 4 | 2 | K. Le Mar, C. McArthur, M. Statham, Home ranges of sympatric red-necked wallabies, redbellied pademelons and common brushtail possums in a temperate eucalypt forestry environment. *Aust. Mammal.* **25**, 183–191 (2003). |
| Diprotodontia | Macropodidae | Western brush wallaby | *Notamacropus irma* | 7 | 6 | L. F. Povh, R. Bencini, B. K. Chambers, T. L. Kreplins, N. Willers, P. J. Adams, J. Wann, H. T. Kobryn, P. A. Fleming, Shedding light on a cryptic macropodid: home ranges and habitat preferences of translocated western brush wallabies (*Notamacropus irma*). *Aust. Mammal.* **41**, 82–91 (2018). |

| | | | | | | |
|---|---|---|---|---|---|---|
| Diprotodontia | Macropodidae | Red-bellied pademelon | *Thylogale billardierii* | 3 | 3 | K. Le Mar, C. McArthur, M. Statham, Home ranges of sympatric red-necked wallabies, redbellied pademelons and common brushtail possums in a temperate eucalypt forestry environment. *Aust. Mammal.* **25**, 183–191 (2003). |
| Diprotodontia | Petauridae | Sugar glider | *Petaurus breviceps* | 9 | 12 | K. A. Nagy, G. C. Suckling, Field energetics and water balance of sugar gliders, *Petaurus breviceps* (Marsupialia: Petauridae). *Aust. J. Zool.* **33**, 683–691 (1985). |
| Diprotodontia | Phalangeridae | Brush-tail possum | *Trichosurus vulpecula hypoleucus* | 43 | 35 | A. Wayne, J. F. Rooney, C. G. Ward, C. V. Vellios, D. B. Lindenmayer, The life history of *Pseudocheirus occidentalis* (Pseudocheiridae) in the jarrah forest of south-western Australia. *Aust. J. Zool.* **53**, 325–337 (2005). |
| Diprotodontia | Phalangeridae | Common brushtail possum | *Trichosurus vulpecula* | 311 | 279 | D. P. Gilmore, Organ-body weight relationships in the common brushtail possum *Trichosurus vulpecula* (Marsupialia: Phalangeridae). *Aust. Mammal.* **7**, 131–138 (1984). |
| Diprotodontia | Phascolarctidae | Koala | *Phascolarctos cinereus* | 35 | 35 | W. A. H. Ellis, F. B. Bercovitch, Body size and sexual selection in the koala. *Behav. Ecol. Sociobiol.* **65**, 1229–1235 (2011). |
| Diprotodontia | Potoroidae | Woylie | *Bettongia penicillata ogilbyi* | 21 | 19 | K. D. Page, L. Ruykys, D. W. Miller, P. J. Adams, P. W. Bateman, P. A. Fleming, Influences of behaviour and physiology on body mass gain in the woylie (*Bettongia penicillata ogilbyi*) post-translocation. *Wildl. Res.* **46**, 429–443 (2019). |
| Diprotodontia | Pseudocheiridae | Western ring-tail possum | *Pseudocheirus occidentalis* | 46 | 77 | A. Wayne, J. F. Rooney, C. G. Ward, C. V. Vellios, D. B. Lindenmayer, The life history of *Pseudocheirus occidentalis* (Pseudocheiridae) in the jarrah forest of south-western Australia. *Aust. J. Zool.* **53**, 325–337 (2005). |

| | | | | | | |
|---|---|---|---|---|---|---|
| Diprotodontia | Vombatidae | Southern hairy-nosed wombat | *Lasiorhinus latifrons* | 26 | 24 | G. R. Finlayson, G. A. Shimmin, P. D. Temple-Smith, K. A. Handasyde, D. A. Taggart, Burrow use and ranging behaviour of the southern hairy-nosed wombat (*Lasiorhinus latifrons*) in the Murraylands, South Australia. *J. Zool.* **265**, 189–200 (2005). |
| Eulipotyphla | Erinaceidae | Four-toed hedgehog | *Atelerix albiventris* | 5 | 7 | O. M. Coker, S. G. Olukole, O. A. Udje, External and internal morphometry of the four-toed hedgehog (*Atelerix albiventris* Wagner, 1841) in Ibadan, Nigeria. *Anim. Res. Int.* **15**, 3003–3012 (2018). |
| Eulipotyphla | Erinaceidae | Algerian hedgehog | *Atelerix algirus* | 4 | 7 | S. García-Rodríguez, X. Puig-Montserrat, Algerian hedgehog (*Atelerix algirus* Lereboullet, 1842) habitat selection at the northern limit of its range. *Galemys, Spanish J. Mammal.* **22**, 49–56 (2014). |
| Eulipotyphla | Erinaceidae | European hedgehog | *Erinaceus europaeus* | 57 | 34 | A. Rautio, A. Valtonen, M. Kunnasranta, The effects of sex and season on home range in european hedgehogs at the northern edge of the species range. *Ann. Zool. Fennici*. **50**, 107–123 (2013). |
| Eulipotyphla | Erinaceidae | Ethiopian hedgehog | *Paraechinus aethiopicus* | 50 | 35 | O. B. Mohammed, N. M. S. Amor, S. A. Omer, A. N. Alagaili, Haematology and biochemistry panels in the Ethiopian hedgehog, *Paraechinus aethiopicus* (Ehrenberg, 1833) from central Saudi Arabia: Establishing reference intervals and assessing variability across sex and hibernation. *J. King Saud Univ. - Sci.* **33**, 101228 (2021). |
| Eulipotyphla | Erinaceidae | Mindanao gymnure | *Podogymnura truei minima* | 33 | 18 | L. R. Heaney, B. R. Tabaranza, E. A. Rickart, D. S. Balete, N. R. Ingle, The mammals of Mt. Kitanglad Nature Park, Mindanao, Philippines. *Fieldiana Zool.* **112**, 1–63 (2006). |
| Eulipotyphla | Soricidae | Short-tailed shrew | *Blarina brevicauda* | 44 | 37 | R. Boonstra, Demography of short-tailed shrew populations living on PCB contaminated sites. **4**, 1–23 (2002). |

| | | | | | | |
|---|---|---|---|---|---|---|
| Eulipotyphla | Soricidae | Buettikofer's shrew | *Crocidura buettikoferi* | 25 | 24 | C. Denys, F. Jacquet, B. Kadjo, A. D. Missoup, V. Aniskine, J. Goüy de Bel-Locq, B. Soropogui, M. Douno, M. Sylla, V. Nicolas, A. Lalis, A. Monadjem, Shrews (Mammalia, Euplipotyphla) from a biodiversity hotspot, Mount Nimba (West Africa), with a field identification key to species. *Zoosystema*. **43**, 729–757 (2021). |
| Eulipotyphla | Soricidae | Ivory Coast white-toothed shrew | *Crocidura eburnea* | 5 | 9 | C. Denys, F. Jacquet, B. Kadjo, A. D. Missoup, V. Aniskine, J. Goüy de Bel-Locq, B. Soropogui, M. Douno, M. Sylla, V. Nicolas, A. Lalis, A. Monadjem, Shrews (Mammalia, Euplipotyphla) from a biodiversity hotspot, Mount Nimba (West Africa), with a field identification key to species. *Zoosystema*. **43**, 729–757 (2021). |
| Eulipotyphla | Soricidae | Large-headed shrew | *Crocidura grandiceps* | 7 | 13 | C. Denys, F. Jacquet, B. Kadjo, A. D. Missoup, V. Aniskine, J. Goüy de Bel-Locq, B. Soropogui, M. Douno, M. Sylla, V. Nicolas, A. Lalis, A. Monadjem, Shrews (Mammalia, Euplipotyphla) from a biodiversity hotspot, Mount Nimba (West Africa), with a field identification key to species. *Zoosystema*. **43**, 729–757 (2021). |
| Eulipotyphla | Soricidae | Jouvenet's shrew | *Crocidura jouvenetae* | 14 | 20 | C. Denys, F. Jacquet, B. Kadjo, A. D. Missoup, V. Aniskine, J. Goüy de Bel-Locq, B. Soropogui, M. Douno, M. Sylla, V. Nicolas, A. Lalis, A. Monadjem, Shrews (Mammalia, Euplipotyphla) from a biodiversity hotspot, Mount Nimba (West Africa), with a field identification key to species. *Zoosystema*. **43**, 729–757 (2021). |
| Eulipotyphla | Soricidae | Crocidura makeda | *Crocidura makeda* | 2 | 3 | A. Konečný, R. Hutterer, Y. Meheretu, J. Bryja, Two new species of *Crocidura* (Mammalia: Soricidae) from Ethiopia and updates on the Ethiopian shrew fauna. *J. Vertebr. Biol.* **69** (2020), doi:10.25225/jvb.20064. |

| | | | | | | |
|---|---|---|---|---|---|---|
| Eulipotyphla | Soricidae | West African long-tailed shrew | *Crocidura muricauda* | 2 | 10 | C. Denys, F. Jacquet, B. Kadjo, A. D. Missoup, V. Aniskine, J. Goüy de Bel-Locq, B. Soropogui, M. Douno, M. Sylla, V. Nicolas, A. Lalis, A. Monadjem, Shrews (Mammalia, Euplipotyphla) from a biodiversity hotspot, Mount Nimba (West Africa), with a field identification key to species. *Zoosystema*. **43**, 729–757 (2021). |
| Eulipotyphla | Soricidae | Nimba shrew | *Crocidura nimbae* | 3 | 4 | C. Denys, F. Jacquet, B. Kadjo, A. D. Missoup, V. Aniskine, J. Goüy de Bel-Locq, B. Soropogui, M. Douno, M. Sylla, V. Nicolas, A. Lalis, A. Monadjem, Shrews (Mammalia, Euplipotyphla) from a biodiversity hotspot, Mount Nimba (West Africa), with a field identification key to species. *Zoosystema*. **43**, 729–757 (2021). |
| Eulipotyphla | Soricidae | Mount Nimba giant forest shrew | *Crocidura nimbasilvanus* | 7 | 4 | C. Denys, F. Jacquet, B. Kadjo, A. D. Missoup, V. Aniskine, J. Goüy de Bel-Locq, B. Soropogui, M. Douno, M. Sylla, V. Nicolas, A. Lalis, A. Monadjem, Shrews (Mammalia, Euplipotyphla) from a biodiversity hotspot, Mount Nimba (West Africa), with a field identification key to species. *Zoosystema*. **43**, 729–757 (2021). |
| Eulipotyphla | Soricidae | West African pygmy shrew | *Crocidura obscurior* | 21 | 22 | C. Denys, F. Jacquet, B. Kadjo, A. D. Missoup, V. Aniskine, J. Goüy de Bel-Locq, B. Soropogui, M. Douno, M. Sylla, V. Nicolas, A. Lalis, A. Monadjem, Shrews (Mammalia, Euplipotyphla) from a biodiversity hotspot, Mount Nimba (West Africa), with a field identification key to species. *Zoosystema*. **43**, 729–757 (2021). |
| Eulipotyphla | Soricidae | African giant shrew | *Crocidura olivieri* | 17 | 12 | C. Denys, F. Jacquet, B. Kadjo, A. D. Missoup, V. Aniskine, J. Goüy de Bel-Locq, B. Soropogui, M. Douno, M. Sylla, V. Nicolas, A. Lalis, A. Monadjem, Shrews (Mammalia, Euplipotyphla) from a biodiversity hotspot, Mount Nimba (West Africa), with a field identification key to species. *Zoosystema*. **43**, 729–757 (2021). |

| | | | | | | |
|---|---|---|---|---|---|---|
| Eulipotyphla | Soricidae | Greater white-toothed shrew | *Crocidura russula* | 76 | 71 | C. Bouteiller-Reuter, N. Perrin, Sex-specific selective pressures on body mass in the greater white-toothed shrew, *Crocidura russula*. *J. Evol. Biol.* **18**, 290–300 (2005). |
| Eulipotyphla | Soricidae | Therese's shrew | *Crocidura theresae* | 20 | 22 | C. Denys, F. Jacquet, B. Kadjo, A. D. Missoup, V. Aniskine, J. Goüy de Bel-Locq, B. Soropogui, M. Douno, M. Sylla, V. Nicolas, A. Lalis, A. Monadjem, Shrews (Mammalia, Euplipotyphla) from a biodiversity hotspot, Mount Nimba (West Africa), with a field identification key to species. *Zoosystema*. **43**, 729–757 (2021). |
| Eulipotyphla | Soricidae | Merida small-eared shrew | *Cryptotis meridensis* | 7 | 7 | N. Woodman, A. Díaz de Pascual, *Cryptotis meridensis*. *Mamm. Species*. **761**, 1 (2004). |
| Eulipotyphla | Soricidae | North American least shrew | *Cryptotis parvus* | 7 | 7 | D. W. Kaufman, G. A. Kaufman, Least shrews in north-central Kansas: Habitat and individual characteristics. *Trans. Kansas Acad. Sci.* **119**, 129–135 (2016). |
| Eulipotyphla | Soricidae | Forest shrew | *Myosorex varius* | 40 | 57 | D. T. Rowe-Rowe, J. Meester, Altitudinal variation in external measurements of two small-mammal species in the Natal Drakensberg. *Ann. Transvaal Museum*. **34**, 49–53 (1985). |
| Eulipotyphla | Soricidae | Water shrew | *Neomys fodiens* | 18 | 21 | L. Balčiauskas, Sex- and age-related differences in tooth row length of small mammals: Shrews. *Acta Zool. Litu.* **14**, 41–47 (2004). |
| Eulipotyphla | Soricidae | Alpine shrew | *Sorex alpinus* | 78 | 57 | I. Baláž, M. Ambros, Shrews (*Sorex* spp.) somatometry and reproduction in Slovakia. *Biologia (Bratisl).* **61**, 611–620 (2006). |

| | | | | | | |
|---|---|---|---|---|---|---|
| Eulipotyphla | Soricidae | Common shrew | *Sorex araneus* | 147 | 113 | L. Balčiauskas, Sex- and age-related differences in tooth row length of small mammals: Shrews. *Acta Zool. Litu.* **14**, 41–47 (2004). |
| Eulipotyphla | Soricidae | Pygmy shrew | *Sorex minutus* | 62 | 39 | L. Balčiauskas, Sex- and age-related differences in tooth row length of small mammals: Shrews. *Acta Zool. Litu.* **14**, 41–47 (2004). |
| Eulipotyphla | Soricidae | Long-clawed shrew | *Sorex unguiculatus* | 5 | 3 | T. Namba, S. D. Ohdachi, Top-down cascade effects of the long-clawed shrew (*Sorex unguiculatus*) on the soil invertebrate community in a cool-temperate forest. *Mammal Study*. **41**, 119–130 (2016). |
| Eulipotyphla | Soricidae | Climbing shrew | *Suncus megalura* | 7 | 10 | C. Denys, F. Jacquet, B. Kadjo, A. D. Missoup, V. Aniskine, J. Goüy de Bel-Locq, B. Soropogui, M. Douno, M. Sylla, V. Nicolas, A. Lalis, A. Monadjem, Shrews (Mammalia, Euplipotyphla) from a biodiversity hotspot, Mount Nimba (West Africa), with a field identification key to species. *Zoosystema*. **43**, 729–757 (2021). |
| Eulipotyphla | Soricidae | Asian house shrew | *Suncus murinus* | 55 | 30 | S. F. Shoma, M. M. Feeroz, M. K. Hasan, Morphometry of Asian house shrew (*Suncus murinus*) in Jahangirnagar University campus. *Bangladesh J. Zool.* **43**, 313–319 (2016). |
| Eulipotyphla | Talpidae | Small Japanese mole | *Mogera imaizumii* | 37 | 37 | T. Hashimoto, M. Abe, Body size and reproductive schedules in two parapatric moles, *Mogera tokudae* and *Mogera imaizumii*, in the Echigo Plain. *Mammal Study*. **26** (2001). |
| Eulipotyphla | Talpidae | Sado mole | *Mogera tokudae* | 50 | 34 | T. Hashimoto, M. Abe, Body size and reproductive schedules in two parapatric moles, *Mogera tokudae* and *Mogera imaizumii*, in the Echigo Plain. *Mammal Study*. **26** (2001). |

| | | | | | | |
|---|---|---|---|---|---|---|
| Eulipotyphla | Talpidae | Eastern mole | *Scalopus aquaticus* | 66 | 100 | G. A. Feldhamer, J. C. Devine, S. M. Ressing, Age-related morphometrics and sex ratios in the eastern mole (*Scalopus aquaticus*). *Am. Midl. Nat.* **169**, 409–415 (2013). |
| Eulipotyphla | Talpidae | European mole | *Talpa europea* | 13 | 8 | G. J. K. Komarnicki, Tissue, sex and age specific accumulation of heavy metals (Zn, Cu, Pb, Cd) by populations of the mole (*Talpa europaea* L.) in a central urban area. *Chemosphere*. **41**, 1593–1602 (2000). |
| Eulipotyphla | Talpidae | Japanese shrew mole | *Urotrichus talpoides* | 36 | 27 | K. Shirai, M. A. Iwasa, A histological study of tail width variation in the greater Japanese shrew-mole, *Urotrichus talpoides*. *Folia Zool.* **66**, 203–210 (2017). |
| Lagomorpha | Leporidae | Snowshoe hare | *Lepus americanus* | 23 | 15 | E. Ellsworth, M. R. Boudreau, K. Nagy, J. L. Rachlow, D. L. Murray, Differential sex-related winter energetics in free-ranging snowshoe hares (*Lepus americanus*). *Can. J. Zool.* **94**, 115–121 (2015). |
| Lagomorpha | Leporidae | Brown hare | *Lepus europaeus* | 289 | 315 | E. Donadio, J. N. Pauli, N. Bonino, A method to estimate body mass and relative age of exotic lagomorphs in the southern Neotropics. *Acta Theriol. (Warsz).* **50**, 81–89 (2005). |
| Lagomorpha | Leporidae | European rabbit | *Oryctolagus cuniculus* | 55 | 86 | E. Donadio, J. N. Pauli, N. Bonino, A method to estimate body mass and relative age of exotic lagomorphs in the southern Neotropics. *Acta Theriol. (Warsz).* **50**, 81–89 (2005). |
| Lagomorpha | Leporidae | Eastern cottontail | *Sylvilagus floridanus* | 10 | 7 | R. G. Tague, Pelvic sexual dimorphism among species monomorphic in body size: Relationship to relative newborn body mass. *J. Mammal.* **97**, 503–517 (2016). |
| Lagomorpha | Leporidae | New England cottontail rabbit | *Sylvilagus transitionalis* | 33 | 19 | M. S. Barbour, J. A. Litvaitis, Niche dimensions of New England cottontails in relation to habitat patch size. *Oecologia*. **95**, 321–327 (1993). |

| | | | | | | |
|---|---|---|---|---|---|---|
| Lagomorpha | Ochotonidae | Plateau pika | *Ochotona curzoniae* | 147 | 140 | H. Nie, J. Liu, B. Chen, Life-history traits and fitness of plateau pika (*Ochotona curzoniae*) in alpine meadow ecosystem. *Ecol. Evol.* **12**, 1–17 (2022). |
| Lagomorpha | Ochotonidae | American pika | *Ochotona princeps* | 30 | 17 | C. A. M. Blair, thesis, University of Alberta (1998). |
| Lagomorpha | Ochotonidae | Collared pika | *Ochotona rufescens* | 105 | 48 | M. M. Shafi, S. W. A. Rizvi, A. Pervez, R. Ali, S. Z. Shah, Some observations on the reproductive biology of collared pika *Ochotona rufescens* Gray, 1842 from Ziarat Valley, Baluchistan, Pakistan. *Acta Theriol. (Warsz).* **37**, 423–427 (1992). |
| Macroscelidea | Macroscelididae | Cape elephant shrew | *Elephantulus edwardii* | 2 | 6 | H. A. Smit, T. J. Robinson, J. Watson, B. J. van Vuuren, A new species of elephant-shrew (Afrotheria: Macroscelidea: *Elephantulus*) from South Africa. *J. Mammal.* **89**, 1257–1269 (2008). |
| Macroscelidea | Macroscelididae | Eastern rock elephant shrew | *Elephantulus myurus* | 36 | 37 | K. Medger, thesis, University of Pretoria (2011). |
| Macroscelidea | Macroscelididae | Western rock elephant shrew | *Elephantulus rupestris* | 6 | 8 | H. A. Smit, T. J. Robinson, J. Watson, B. J. van Vuuren, A new species of elephant-shrew (Afrotheria: Macroscelidea: *Elephantulus*) from South Africa. *J. Mammal.* **89**, 1257–1269 (2008). |
| Macroscelidea | Macroscelididae | Karoo rock elephant shrew | *Elephantulus pilicaudus* | 6 | 6 | H. A. Smit, T. J. Robinson, J. Watson, B. J. van Vuuren, A new species of elephant-shrew (Afrotheria: Macroscelidea: *Elephantulus*) from South Africa. *J. Mammal.* **89**, 1257–1269 (2008). |

| | | | | | | |
|---|---|---|---|---|---|---|
| Macroscelidea | Macroscelididae | Round-eared sengi | *Macroscelides proboscideus* | 33 | 28 | M. Schubert, N. Pillay, D. O. Ribble, C. Schradin, The round-eared sengi and the evolution of social monogamy: Factors that constrain males to live with a single female. *Ethology*. **115**, 972–985 (2009). |
| Peramelemorphia | Peramelidae | Brown bandicoot | *Isoodon obesulus* | 31 | 36 | D. M. Stoddart, R. W. Braithwaite, A strategy for utilization of regenerating heathland habitat by the brown bandicoot (*Isoodon obesulus*; Marsupialia, Peramelidae). *J. Anim. Ecol.* **48**, 165–179 (1979). |
| Peramelemorphia | Peramelidae | Northern brown bandicoot | *Isoodon macrourus* | 82 | 56 | C. Kemper, D. J. Kitchener, W. F. Humphreys, R. A. How, L. H. Schmitt, A. Bradley, The biology of the northern brown bandicoot, *Isoodon macrourus* (Marsupialia: Peramelidae) at Mitchell Plateau, Western Australia. *Aust. J. Zool.* **37**, 609–616 (1990). |
| Peramelemorphia | Peramelidae | Eastern barred bandicoot | *Perameles gunnii* | 396 | 403 | S. A. Mallick, M. M. Driessen, G. J. Hocking, Demography and home range of the eastern barred bandicoot (*Perameles gunnii*) in south-eastern Tasmania. *Wildl. Res.* **27**, 103–115 (2000). |
| Peramelemorphia | Peramelidae | Long-nosed bandicoot | *Perameles nasuta* | 14 | 6 | C. I. MacGregor, J. T. Wood, N. Dexter, D. B. Lindenmayer, Home range size and use by long-nosed bandicoot (*Perameles nasuta*) following fire. *Aust. Mammal.* **35**, 206–216 (2013). |

| | | | | | | |
|---|---|---|---|---|---|---|
| Perissodactyla | Tapiridae | South American tapir | *Tapirus terrestris* | 4 | 2 | F. Gonçalves, R. S. Bovendorp, G. Beca, C. Bello, R. Costa-Pereira, R. L. Muylaert, R. R. Rodarte, N. Villar, R. Souza, M. E. Graipel, J. J. Cherem, D. Faria, J. Baumgarten, M. R. Alvarez, E. M. Vieira, N. Cáceres, R. Pardini, Y. L. R. Leite, L. P. Costa, M. A. R. Mello, E. Fischer, F. C. Passos, L. H. Varzinczak, J. A. Prevedello, A. P. Cruz-Neto, F. Carvalho, A. R. Percequillo, A. Paviolo, A. Nava, J. M. B. Duarte, N. U. de la Sancha, E. Bernard, R. G. Morato, J. F. Ribeiro, R. G. Becker, G. Paise, P. S. Tomasi, F. Vélez-Garcia, G. L. Melo, J. Sponchiado, F. Cerezer, M. A. S. Barros, A. Q. S. de Souza, C. C. dos Santos, G. A. F. Giné, P. Kerches-Rogeri, M. M. Weber, G. Ambar, L. V. Cabrera-Martinez, A. Eriksson, M. Silveira, C. F. Santos, L. Alves, E. Barbier, G. C. Rezende, G. S. T. Garbino, É. O. Rios, A. Silva, A. T. A. Nascimento, R. S. de Carvalho, A. Feijó, J. Arrabal, I. Agostini, D. Lamattina, S. Costa, E. Vanderhoeven, F. R. de Melo, P. de Oliveira Laroque, L. Jerusalinsky, M. M. Valença-Montenegro, A. B. Martins, G. Ludwig, R. B. de Azevedo, A. Anzóategui, M. X. da Silva, M. Figuerêdo Duarte Moraes, A. Vogliotti, A. Gatti, T. Püttker, C. S. Barros, T. K. Martins, A. Keuroghlian, D. P. Eaton, C. L. Neves, M. S. Nardi, C. Braga, P. R. Gonçalves, A. C. Srbek-Araujo, P. Mendes, J. A. de Oliveira, F. A. M. Soares, P. A. Rocha, P. Crawshaw, M. C. Ribeiro, M. Galetti, Atlantic Mammal Traits: a data set of morphological traits of mammals in the Atlantic Forest of South America. *Ecology*. **99**, 498 (2018). |
| Pilosa | Bradypodidae | Maned sloth | *Bradypus torquatus* | 15 | 21 | P. Lara-Ruiz, A. G. Chiarello, Life-history traits and sexual dimorphism of the Atlantic forest maned sloth *Bradypus torquatus* (Xenarthra: Bradypodidae). *J. Zool.* **267**, 63–73 (2005). |

| | | | | | | |
|---|---|---|---|---|---|---|
| Pilosa | Bradypodidae | Pale-throated three-toed sloth | *Bradypus tridactylus* | 46 | 44 | M. Pool, R. Boateng, A.-M. Ako-Adounvo, R. Allen-McFarlane, D. Elizondo, H. Paturault, H. Alhawas, G. Middendorf, Sloths in the city: unexpectedly high density of pale-throated three-toed sloths (*Bradypus tridactylus*) found in an urban forest patch in Paramaribo, Suriname. *Edentata Newsl. IUCN/SSC Anteater, Sloth Armadillo Spec. Gr.* (2016), doi:10.2305/iucn.ch.2016.edentata-17-1.5.en. |

| | | | | | | |
|---|---|---|---|---|---|---|
| Pilosa | Bradypodidae | Brown-throated sloth | *Bradypus variegatus* | 4 | 5 | F. Gonçalves, R. S. Bovendorp, G. Beca, C. Bello, R. Costa-Pereira, R. L. Muylaert, R. R. Rodarte, N. Villar, R. Souza, M. E. Graipel, J. J. Cherem, D. Faria, J. Baumgarten, M. R. Alvarez, E. M. Vieira, N. Cáceres, R. Pardini, Y. L. R. Leite, L. P. Costa, M. A. R. Mello, E. Fischer, F. C. Passos, L. H. Varzinczak, J. A. Prevedello, A. P. Cruz-Neto, F. Carvalho, A. R. Percequillo, A. Paviolo, A. Nava, J. M. B. Duarte, N. U. de la Sancha, E. Bernard, R. G. Morato, J. F. Ribeiro, R. G. Becker, G. Paise, P. S. Tomasi, F. Vélez-Garcia, G. L. Melo, J. Sponchiado, F. Cerezer, M. A. S. Barros, A. Q. S. de Souza, C. C. dos Santos, G. A. F. Giné, P. Kerches-Rogeri, M. M. Weber, G. Ambar, L. V. Cabrera-Martinez, A. Eriksson, M. Silveira, C. F. Santos, L. Alves, E. Barbier, G. C. Rezende, G. S. T. Garbino, É. O. Rios, A. Silva, A. T. A. Nascimento, R. S. de Carvalho, A. Feijó, J. Arrabal, I. Agostini, D. Lamattina, S. Costa, E. Vanderhoeven, F. R. de Melo, P. de Oliveira Laroque, L. Jerusalinsky, M. M. Valença-Montenegro, A. B. Martins, G. Ludwig, R. B. de Azevedo, A. Anzóategui, M. X. da Silva, M. Figuerêdo Duarte Moraes, A. Vogliotti, A. Gatti, T. Püttker, C. S. Barros, T. K. Martins, A. Keuroghlian, D. P. Eaton, C. L. Neves, M. S. Nardi, C. Braga, P. R. Gonçalves, A. C. Srbek-Araujo, P. Mendes, J. A. de Oliveira, F. A. M. Soares, P. A. Rocha, P. Crawshaw, M. C. Ribeiro, M. Galetti, Atlantic Mammal Traits: a data set of morphological traits of mammals in the Atlantic Forest of South America. *Ecology*. **99**, 498 (2018). |
| Pilosa | Choloepodidae | Hoffman's two-toed sloth | *Choloepus hoffmanni* | 38 | 48 | M. Z. Peery, J. N. Pauli, The mating system of a “lazy” mammal, Hoffmann’s two-toed sloth. *Anim. Behav.* **84**, 555–562 (2012). |

| | | | | | | |
|---|---|---|---|---|---|---|
| Pilosa | Myrmecophagidae | Giant anteater | *Myrmecophaga tridactyla* | 19 | 27 | D. Kluyber, N. Attias, M. H. Alves, A. C. Alves, G. Massocato, A. L. J. Desbiez, Physical capture and chemical immobilization procedures for a mammal with singular anatomy: the giant anteater (*Myrmecophaga tridactyla*). *Eur. J. Wildl. Res.* **67**, 1–10 (2021). |
| Primates | Atelidae | Red-handed howler | *Alouatta belzebul* | 4 | 4 | F. Gonçalves, R. S. Bovendorp, G. Beca, C. Bello, R. Costa-Pereira, R. L. Muylaert, R. R. Rodarte, N. Villar, R. Souza, M. E. Graipel, J. J. Cherem, D. Faria, J. Baumgarten, M. R. Alvarez, E. M. Vieira, N. Cáceres, R. Pardini, Y. L. R. Leite, L. P. Costa, M. A. R. Mello, E. Fischer, F. C. Passos, L. H. Varzinczak, J. A. Prevedello, A. P. Cruz-Neto, F. Carvalho, A. R. Percequillo, A. Paviolo, A. Nava, J. M. B. Duarte, N. U. de la Sancha, E. Bernard, R. G. Morato, J. F. Ribeiro, R. G. Becker, G. Paise, P. S. Tomasi, F. Vélez-Garcia, G. L. Melo, J. Sponchiado, F. Cerezer, M. A. S. Barros, A. Q. S. de Souza, C. C. dos Santos, G. A. F. Giné, P. Kerches-Rogeri, M. M. Weber, G. Ambar, L. V. Cabrera-Martinez, A. Eriksson, M. Silveira, C. F. Santos, L. Alves, E. Barbier, G. C. Rezende, G. S. T. Garbino, É. O. Rios, A. Silva, A. T. A. Nascimento, R. S. de Carvalho, A. Feijó, J. Arrabal, I. Agostini, D. Lamattina, S. Costa, E. Vanderhoeven, F. R. de Melo, P. de Oliveira Laroque, L. Jerusalinsky, M. M. Valença-Montenegro, A. B. Martins, G. Ludwig, R. B. de Azevedo, A. Anzóategui, M. X. da Silva, M. Figuerêdo Duarte Moraes, A. Vogliotti, A. Gatti, T. Püttker, C. S. Barros, T. K. Martins, A. Keuroghlian, D. P. Eaton, C. L. Neves, M. S. Nardi, C. Braga, P. R. Gonçalves, A. C. Srbek-Araujo, P. Mendes, J. A. de Oliveira, F. A. M. Soares, P. A. Rocha, P. Crawshaw, M. C. Ribeiro, M. Galetti, Atlantic Mammal Traits: a data set of morphological traits of mammals in the Atlantic Forest of South America. *Ecology*. **99**, 498 (2018). |

| | | | | | | |
|---|---|---|---|---|---|---|
| Primates | Atelidae | Black howler monkey | *Alouatta caraya* | 58 | 11 7 | R. J. Smith, W. L. Jungers, Body mass in comparative primatology. *J. Hum. Evol.* **32**, 523–559 (1997).; N. Rowe, M. Myers, “All the World’s Primates” (Charlestown RI, 2022), (available at www.alltheworldsprimates.org). |
| Primates | Atelidae | Mantled howler monkey | *Alouatta palliata* | 34 4 | 73 8 | N. Rowe, M. Myers, “All the World’s Primates” (Charlestown RI, 2022), (available at www.alltheworldsprimates.org). |
| Primates | Atelidae | Columbian red howler | *Alouatta seniculus* | 31 | 29 | R. J. Smith, W. L. Jungers, Body mass in comparative primatology. *J. Hum. Evol.* **32**, 523–559 (1997).; N. Rowe, M. Myers, “All the World’s Primates” (Charlestown RI, 2022), (available at www.alltheworldsprimates.org). |

| | | | | | | |
|---|---|---|---|---|---|---|
| Primates | Cebidae | Cotton-top tamarin | *Saguinus oedipus* | 181 | 171 | R. J. Smith, W. L. Jungers, Body mass in comparative primatology. *J. Hum. Evol.* **32**, 523–559 (1997). |
| Primates | Cebidae | Azara's night monkey | *Aotus azarae* | 45 | 46 | Fernandez-Duque 2011 |
| Primates | Cebidae | Gray-legged night monkey | *Aotus griseimembra* | 20 | 16 | N. Rowe, M. Myers, “All the World’s Primates” (Charlestown RI, 2022), (available at www.alltheworldsprimates.org). |
| Primates | Cebidae | Lemurine night monkey | *Aotus lemurinus* | 7 | 6 | N. Rowe, M. Myers, “All the World’s Primates” (Charlestown RI, 2022), (available at www.alltheworldsprimates.org). |
| Primates | Cebidae | Nancy Ma's night monkey | *Aotus nancymaae* | 130 | 124 | R. J. Smith, W. L. Jungers, Body mass in comparative primatology. *J. Hum. Evol.* **32**, 523–559 (1997). |
| Primates | Cebidae | Humboldt's white-fronted capuchin | *Cebus albifrons* | 24 | 43 | R. J. Smith, W. L. Jungers, Body mass in comparative primatology. *J. Hum. Evol.* **32**, 523–559 (1997). |
| Primates | Cebidae | Guianan squirrel monkey | *Saimiri sciureus* | 17 | 58 | R. J. Smith, W. L. Jungers, Body mass in comparative primatology. *J. Hum. Evol.* **32**, 523–559 (1997). |

| | | | | | | |
|---|---|---|---|---|---|---|
| Primates | Cercopithecidae | Tana river mangabey | *Cercocebus galeritus* | 2 | 2 | M. Colyn, Données pondérales sur les primates Cercopithecidae d’Afrique Centrale (Bassin du Zaire/Congo). *Mammalia*. **58**, 483–487 (1994). |
| Primates | Cercopithecidae | Vervet monkey | *Cercopithecus aethiops* | 26 | 36 | R. J. Smith, W. L. Jungers, Body mass in comparative primatology. *J. Hum. Evol.* **32**, 523–559 (1997). |
| Primates | Cercopithecidae | Red-tailed monkey | *Cercopithecus ascanius* | 32 | 187 | M. Colyn, Données pondérales sur les primates Cercopithecidae d’Afrique Centrale (Bassin du Zaire/Congo). *Mammalia*. **58**, 483–487 (1994). |
| Primates | Cercopithecidae | Hamlyn's monkey | *Cercopithecus hamlyni* | 11 | 9 | M. Colyn, Données pondérales sur les primates Cercopithecidae d’Afrique Centrale (Bassin du Zaire/Congo). *Mammalia*. **58**, 483–487 (1994). |
| Primates | Cercopithecidae | L'Hoest's monkey | *Cercopithecus l'hoesti* | 19 | 50 | M. Colyn, Données pondérales sur les primates Cercopithecidae d’Afrique Centrale (Bassin du Zaire/Congo). *Mammalia*. **58**, 483–487 (1994). |
| Primates | Cercopithecidae | Blue monkey | *Cercopithecus mitis* | 41 | 94 | M. Colyn, Données pondérales sur les primates Cercopithecidae d’Afrique Centrale (Bassin du Zaire/Congo). *Mammalia*. **58**, 483–487 (1994). |
| Primates | Cercopithecidae | Greater spot-nosed monkey | *Cercopithecus nictitans* | 17 | 21 | M. Colyn, Données pondérales sur les primates Cercopithecidae d’Afrique Centrale (Bassin du Zaire/Congo). *Mammalia*. **58**, 483–487 (1994). |
| Primates | Cercopithecidae | Grivet | *Chlorocebus aethiops* | 48 | 61 | F. Anapol, T. R. Turner, C. S. Mott, C. J. Jolly, Comparative postcranial body shape and locomotion in *Chlorocebus aethiops* and *Cercopithecus mitis*. *Am. J. Phys. Anthropol.* **127**, 231–239 (2005). |

| | | | | | | |
|---|---|---|---|---|---|---|
| Primates | Cercopitheci dae | Angola colobus | *Colobus angolensis* | 8 | 5 | M. Colyn, Données pondérales sur les primates Cercopithecidae d'Afrique Centrale (Bassin du Zaire/Congo). *Mammalia*. **58**, 483–487 (1994). |
| Primates | Cercopitheci dae | Grey-cheeked mangabey | *Lophocebus albigena* | 4 | 6 | M. Colyn, Données pondérales sur les primates Cercopithecidae d'Afrique Centrale (Bassin du Zaire/Congo). *Mammalia*. **58**, 483–487 (1994). |
| Primates | Cercopitheci dae | Olive baboon | *Papio anubis* | 39 | 35 | R. J. Smith, W. L. Jungers, Body mass in comparative primatology. *J. Hum. Evol.* **32**, 523–559 (1997). |
| Primates | Cercopitheci dae | Yellow baboon | *Papio cynocephalus* | 20 | 18 | J. Altmann, D. Schoeller, S. A. Altmann, P. Muruthi, R. M. Sapolsky, Body size and fatness of free-living baboons reflect food availability and activity levels. *Am. J. Primatol.* **30**, 149–161 (1993). |
| Primates | Cercopitheci dae | Hamadryas baboon | *Papio hamadryas* | 41 | 39 | R. J. Smith, W. L. Jungers, Body mass in comparative primatology. *J. Hum. Evol.* **32**, 523–559 (1997). |
| Primates | Cheirogaleid ae | Greater dwarf lemur | *Cheirogaleus major* | 3 | 3 | P. M. Kappeler, Patterns of sexual dimorphism in body weight among prosimian primates. *Folia Primatol.* **57**, 132–146 (1991). |
| Primates | Cheirogaleid ae | Fat-tailed dwarf lemur | *Cheirogaleus medius* | 23 | 25 | P. M. Kappeler, Patterns of sexual dimorphism in body weight among prosimian primates. *Folia Primatol.* **57**, 132–146 (1991). |
| Primates | Cheirogaleid ae | Gray mouse lemur | *Microcebus murinus* | 33 | 27 | P. M. Kappeler, Patterns of sexual dimorphism in body weight among prosimian primates. *Folia Primatol.* **57**, 132–146 (1991). |

| | | | | | | |
|---|---|---|---|---|---|---|
| Primates | Cheirogaleidae | Coquerel's giant mouse lemur | *Mirza coquereli* | 10 | 9 | P. M. Kappeler, Patterns of sexual dimorphism in body weight among prosimian primates. *Folia Primatol.* **57**, 132–146 (1991). |
| Primates | Daubentoniidae | Aye-aye | *Daubentonia madagascariensis* | 2 | 2 | P. M. Kappeler, Patterns of sexual dimorphism in body weight among prosimian primates. *Folia Primatol.* **57**, 132–146 (1991). |
| Primates | Galagidae | Brown greater galago | *Otolemur crassicaudatus argentatus* | 17 | 19 | P. M. Kappeler, Patterns of sexual dimorphism in body weight among prosimian primates. *Folia Primatol.* **57**, 132–146 (1991). |
| Primates | Galagidae | Northern greater galago | *Otolemur garnettii* | 17 | 19 | P. M. Kappeler, Patterns of sexual dimorphism in body weight among prosimian primates. *Folia Primatol.* **57**, 132–146 (1991). |
| Primates | Galagidae | Mohol bushbaby | *Galago moholi* | 25 | 20 | P. M. Kappeler, Patterns of sexual dimorphism in body weight among prosimian primates. *Folia Primatol.* **57**, 132–146 (1991). |
| Primates | Galagidae | Prince Demidoff's bushbaby | *Galagoides demidovii* | 4 | 8 | P. M. Kappeler, Patterns of sexual dimorphism in body weight among prosimian primates. *Folia Primatol.* **57**, 132–146 (1991). |
| Primates | Galagidae | Zanzibar bushbaby | *Galagoides zanzibaricus* | 35 | 38 | R. J. Smith, W. L. Jungers, Body mass in comparative primatology. *J. Hum. Evol.* **32**, 523–559 (1997). |
| Primates | Galagidae | *Otolemur crassicaudatus* | | 40 | 44 | P. M. Kappeler, Patterns of sexual dimorphism in body weight among prosimian primates. *Folia Primatol.* **57**, 132–146 (1991). |

| | | | | | | |
|---|---|---|---|---|---|---|
| Primates | Hominidae | Chimpanzee | *Pan troglodytes* | 18 | 23 | R. J. Smith, W. L. Jungers, Body mass in comparative primatology. *J. Hum. Evol.* **32**, 523–559 (1997). |
| Primates | Hylobatidae | Lar gibbon | *Hylobates lar* | 84 | 66 | R. J. Smith, W. L. Jungers, Body mass in comparative primatology. *J. Hum. Evol.* **32**, 523–559 (1997). |
| Primates | Hylobatidae | Siamang | *Hylobates syndactylus* | 89 | 87 | R. J. Smith, W. L. Jungers, Body mass in comparative primatology. *J. Hum. Evol.* **32**, 523–559 (1997). |
| Primates | Indriidae | Coquerel's sifaka | *Propithecus coquereli* | 8 | 10 | P. M. Kappeler, Patterns of sexual dimorphism in body weight among prosimian primates. *Folia Primatol.* **57**, 132–146 (1991). |
| Primates | Indriidae | Golden-crowned sifaka | *Propithecus tattersalli* | 4 | 5 | P. M. Kappeler, Patterns of sexual dimorphism in body weight among prosimian primates. *Folia Primatol.* **57**, 132–146 (1991). |
| Primates | Indriidae | Verreaux's sifaka | *Propithecus verreauxi* | 10 | 12 | P. M. Kappeler, Patterns of sexual dimorphism in body weight among prosimian primates. *Folia Primatol.* **57**, 132–146 (1991). |
| Primates | Lemuridae | White-headed lemur | *Eulemur albifrons* | 13 | 11 | P. M. Kappeler, Patterns of sexual dimorphism in body weight among prosimian primates. *Folia Primatol.* **57**, 132–146 (1991). |
| Primates | Lemuridae | Collared brown lemur | *Eulemur collaris* | 13 | 4 | P. M. Kappeler, Patterns of sexual dimorphism in body weight among prosimian primates. *Folia Primatol.* **57**, 132–146 (1991). |

| | | | | | | |
|---|---|---|---|---|---|---|
| Primates | Lemuridae | Blue-eyed black lemur | *Eulemur flavifrons* | 3 | 5 | P. M. Kappeler, Patterns of sexual dimorphism in body weight among prosimian primates. *Folia Primatol.* **57**, 132–146 (1991). |
| Primates | Lemuridae | Common brown lemur | *Eulemur fulvus* | 60 | 47 | P. M. Kappeler, Patterns of sexual dimorphism in body weight among prosimian primates. *Folia Primatol.* **57**, 132–146 (1991). |
| Primates | Lemuridae | Black lemur | *Eulemur macaco* | 23 | 20 | P. M. Kappeler, Patterns of sexual dimorphism in body weight among prosimian primates. *Folia Primatol.* **57**, 132–146 (1991). |
| Primates | Lemuridae | Mongoose lemur | *Eulemur mongoz* | 14 | 13 | P. M. Kappeler, Patterns of sexual dimorphism in body weight among prosimian primates. *Folia Primatol.* **57**, 132–146 (1991). |
| Primates | Lemuridae | Red-bellied lemur | *Eulemur rubriventer* | 4 | 4 | P. M. Kappeler, Patterns of sexual dimorphism in body weight among prosimian primates. *Folia Primatol.* **57**, 132–146 (1991). |
| Primates | Lemuridae | Red lemur | *Eulemur rufus* | 22 | 17 | P. M. Kappeler, Patterns of sexual dimorphism in body weight among prosimian primates. *Folia Primatol.* **57**, 132–146 (1991). |
| Primates | Lemuridae | Sanford's brown lemur | *Eulemur sanfordi* | 7 | 6 | P. M. Kappeler, Patterns of sexual dimorphism in body weight among prosimian primates. *Folia Primatol.* **57**, 132–146 (1991). |
| Primates | Lemuridae | Eastern lesser bamboo lemur | *Hapalemur griseus griseus* | 10 | 9 | P. M. Kappeler, Patterns of sexual dimorphism in body weight among prosimian primates. *Folia Primatol.* **57**, 132–146 (1991). |

| | | | | | | |
|---|---|---|---|---|---|---|
| Primates | Lemuridae | Ringtailed lemur | *Lemur catta* | 24 | 16 | P. M. Kappeler, Patterns of sexual dimorphism in body weight among prosimian primates. *Folia Primatol.* **57**, 132–146 (1991). |
| Primates | Lemuridae | Crowned lemur | *Lemur coronatus* | 11 | 8 | P. M. Kappeler, Patterns of sexual dimorphism in body weight among prosimian primates. *Folia Primatol.* **57**, 132–146 (1991). |
| Primates | Lemuridae | Black-and-white ruffed lemur | *Varecia variegata* | 46 | 35 | P. M. Kappeler, Patterns of sexual dimorphism in body weight among prosimian primates. *Folia Primatol.* **57**, 132–146 (1991). |
| Primates | Lemuridae | Red ruffed lemur | *Varecia variegata rubra* | 20 | 17 | P. M. Kappeler, Patterns of sexual dimorphism in body weight among prosimian primates. *Folia Primatol.* **57**, 132–146 (1991). |
| Primates | Lepilemuridae | Milne-Edward's sportive lemur | *Lepilemur edwardsi* | 6 | 6 | N. Rowe, M. Myers, "All the World's Primates" (Charlestown RI, 2022), (available at www.alltheworldsprimates.org). |
| Primates | Lepilemuridae | Petter's sportive lemur | *Lepilemur petteri* | 8 | 2 | N. Rowe, M. Myers, "All the World's Primates" (Charlestown RI, 2022), (available at www.alltheworldsprimates.org). |
| Primates | Lepilemuridae | Red-tailed sportive lemur | *Lepilemur ruficaudatus* | 15 | 19 | N. Rowe, M. Myers, "All the World's Primates" (Charlestown RI, 2022), (available at www.alltheworldsprimates.org). |
| Primates | Lorisidae | Red slender loris | *Loris tardigradus malabaricus* | 10 | 8 | P. M. Kappeler, Patterns of sexual dimorphism in body weight among prosimian primates. *Folia Primatol.* **57**, 132–146 (1991). |

| | | | | | | |
|---|---|---|---|---|---|---|
| Primates | Lorisidae | Sunda slow loris | *Nycticebus coucang* | 14 | 15 | P. M. Kappeler, Patterns of sexual dimorphism in body weight among prosimian primates. *Folia Primatol.* **57**, 132–146 (1991). |
| Primates | Lorisidae | Pygmy slow loris | *Nycticebus pygmaeus* | 7 | 5 | P. M. Kappeler, Patterns of sexual dimorphism in body weight among prosimian primates. *Folia Primatol.* **57**, 132–146 (1991). |
| Primates | Lorisidae | Potto | *Perodicticus potto* | 5 | 6 | P. M. Kappeler, Patterns of sexual dimorphism in body weight among prosimian primates. *Folia Primatol.* **57**, 132–146 (1991). |

| | | | | | | |
|---|---|---|---|---|---|---|
| Primates | Pitheciidae | Black-fronted titi | *Callicebus nigrifrons* | 2 | 4 | F. Gonçalves, R. S. Bovendorp, G. Beca, C. Bello, R. Costa-Pereira, R. L. Muylaert, R. R. Rodarte, N. Villar, R. Souza, M. E. Graipel, J. J. Cherem, D. Faria, J. Baumgarten, M. R. Alvarez, E. M. Vieira, N. Cáceres, R. Pardini, Y. L. R. Leite, L. P. Costa, M. A. R. Mello, E. Fischer, F. C. Passos, L. H. Varzinczak, J. A. Prevedello, A. P. Cruz-Neto, F. Carvalho, A. R. Percequillo, A. Paviolo, A. Nava, J. M. B. Duarte, N. U. de la Sancha, E. Bernard, R. G. Morato, J. F. Ribeiro, R. G. Becker, G. Paise, P. S. Tomasi, F. Vélez-Garcia, G. L. Melo, J. Sponchiado, F. Cerezer, M. A. S. Barros, A. Q. S. de Souza, C. C. dos Santos, G. A. F. Giné, P. Kerches-Rogeri, M. M. Weber, G. Ambar, L. V. Cabrera-Martinez, A. Eriksson, M. Silveira, C. F. Santos, L. Alves, E. Barbier, G. C. Rezende, G. S. T. Garbino, É. O. Rios, A. Silva, A. T. A. Nascimento, R. S. de Carvalho, A. Feijó, J. Arrabal, I. Agostini, D. Lamattina, S. Costa, E. Vanderhoeven, F. R. de Melo, P. de Oliveira Laroque, L. Jerusalinsky, M. M. Valença-Montenegro, A. B. Martins, G. Ludwig, R. B. de Azevedo, A. Anzóategui, M. X. da Silva, M. Figuerêdo Duarte Moraes, A. Vogliotti, A. Gatti, T. Püttker, C. S. Barros, T. K. Martins, A. Keuroghlian, D. P. Eaton, C. L. Neves, M. S. Nardi, C. Braga, P. R. Gonçalves, A. C. Srbek-Araujo, P. Mendes, J. A. de Oliveira, F. A. M. Soares, P. A. Rocha, P. Crawshaw, M. C. Ribeiro, M. Galetti, Atlantic Mammal Traits: a data set of morphological traits of mammals in the Atlantic Forest of South America. *Ecology*. **99**, 498 (2018). |
| Primates | Tarsiidae | Philippine tarsier | *Carlito syrichta* | 10 | 17 | P. M. Kappeler, Patterns of sexual dimorphism in body weight among prosimian primates. *Folia Primatol.* **57**, 132–146 (1991). |

| | | | | | | |
|---|---|---|---|---|---|---|
| Primates | Tarsiidae | Horsfield's tarsier | *Cephalopachus bancanus* | 3 | 3 | P. M. Kappeler, Patterns of sexual dimorphism in body weight among prosimian primates. *Folia Primatol.* **57**, 132–146 (1991). |
| Rodentia | Bathyergidae | Cape dune mole-rat | *Bathyergus suillus* | 11 | 12 | H. G. Thomas, P. W. Bateman, M. Scantlebury, N. C. Bennett, Seasonal effects on digging activity and burrow architecture in the Cape dune mole-rat, *Bathyergus suillus* (Rodentia: Bathyergidae). *African Zool.* **47**, 332–340 (2012). |
| Rodentia | Bathyergidae | Damaraland mole-rat | *Cryptomys damarensis* | 11 | 14 | N. C. Bennett, J. U. M. Jarvis, D. B. Wallace, The relative age structure and body masses of complete wild-captured colonies of two social mole-rats, the common mole-rat, *Cryptomys hottentotus hottentotus* and the Damaraland mole-rat, *Cryptomys damarensis*. *J. Zool.* **220**, 469–485 (1990). |
| Rodentia | Bathyergidae | Common mole-rat | *Cryptomys hottentotus hottentotus* | 8 | 5 | N. C. Bennett, J. U. M. Jarvis, D. B. Wallace, The relative age structure and body masses of complete wild-captured colonies of two social mole-rats, the common mole-rat, *Cryptomys hottentotus hottentotus* and the Damaraland mole-rat, *Cryptomys damarensis*. *J. Zool.* **220**, 469–485 (1990). |
| Rodentia | Bathyergidae | Giant mole-rat | *Cryptomys mechowi* | 18 | 15 | A. I. Schulte-Hostedde, "Sexual size dimorphism in rodents" in *Rodent Societies: An Ecological and Evolutionary Perspective*, J. O. Wolff, P. W. Sherman, Eds. (University of Chicago Press, Chicago, 2007), pp. 115–128. |
| Rodentia | Bathyergidae | Ansell's mole-rat | *Fukomys anselli* | 30 | 40 | J. Šklíba, V. Mazoch, H. Patzenhauerová, E. Hrouzková, M. Lövy, O. Kott, R. Šumbera, A maze-lover’s dream: Burrow architecture, natural history and habitat characteristics of Ansell’s mole-rat (*Fukomys anselli*). *Mamm. Biol.* **77**, 420–427 (2012). |

| | | | | | | |
|---|---|---|---|---|---|---|
| Rodentia | Bathyergidae | Micklem's mole-rat | *Fukomys micklemi* | 7 | 13 | P. A. A. G. Van Daele, N. Desmet, R. Šumbera, D. Adriaens, Work behaviour and biting performance in the cooperative breeding Micklem’s mole-rat *Fukomys micklemi* (Bathyergidae, Rodentia). *Mamm. Biol.* **95**, 69–76 (2019). |
| Rodentia | Bathyergidae | Cape mole-rat | *Georychus capensis* | 11 | 29 | H. G. Thomas, P. W. Bateman, M. Scantlebury, N. C. Bennett, Season but not sex influences burrow length and complexity in the non-sexually dimorphic solitary Cape mole-rat (Rodentia: Bathyergidae). *J. Zool.* **288**, 214–221 (2012). |
| Rodentia | Bathyergidae | Silvery mole-rat | *Heliophobius argente-eocinereus* | 74 | 70 | A. I. Schulte-Hostedde, "Sexual size dimorphism in rodents" in *Rodent Societies: An Ecological and Evolutionary Perspective*, J. O. Wolff, P. W. Sherman, Eds. (University of Chicago Press, Chicago, 2007), pp. 115–128. |
| Rodentia | Capromyidae | Black-tailed hutia | *Mesocapromys melanurus* | 27 | 13 | R. Borroto-Páez, A. C. Pérez, Systematics and comparative biology of the black-tailed hutia (*Mesocapromys melanurus*). *Terr. Mamm. West Indies*. **10**, 90–98 (2012). |
| Rodentia | Castoridae | Canadian beaver | *Castor canadensis* | 21 | 22 | D. W. Soprovich, thesis, University of Manitoba (1994). |
| Rodentia | Caviidae | Greater guinea pig | *Cavia magna* | 18 | 22 | C. Kraus, F. Trillmich, J. Künkele, Reproduction and growth in a precocial small mammal, *Cavia magna*. *J. Mammal.* **86**, 763–772 (2005). |
| Rodentia | Caviidae | Capybara | *Hydrochoerus hydrochaeris* | 9 | 6 | K. M. P. M. de B. Ferraz, K. Bonach, L. M. Verdade, Relationship between body mass and body length in capybaras (*Hydrochoerus hydrochaeris*). *Biota Neotrop.* **5**, 197–200 (2005). |

| | | | | | | |
|---|---|---|---|---|---|---|
| Rodentia | Cricetidae | Chilomys instans | *Chilomys instans* | 7 | 3 | D. Ocampo, K. G. Borja-Acosta, J. Lozano-Flórez, S. Cifuentes-Acevedo, E. Arbeláez-Cortés, N. J. Bayly, Á. Caguazango, B. Coral-Jaramillo, D. Cueva, F. Forero, J. P. Gómez, C. Gómez, M. A. Loaiza-Muñoz, G. A. Londoño, S. Losada-Prado, S. Pérez-Peña, H. E. Ramírez-Chaves, M. E. Rodríguez-Posada, J. Sanabria-Mejía, M. Sánchez-Martínez, V. H. Serrano-Cardozo, M. S. Sierra-Buitrago, J. Soto-Patiño, O. Acevedo-Charry, Body mass data set for 1,317 bird and 270 mammal species from Colombia. *Ecology*. **102** (2021), doi:10.1002/ecy.3273. |
| Rodentia | Cricetidae | Red-backed vole | *Clethrionomys gapperi* | 74 | 60 | S. Bondrup-Nielsen, R. A. Ims, Reversed sexual size dimorphism in microtines: Are females larger than males or are males smaller than females? *Evol. Ecol.* **4**, 261–272 (1990). |
| Rodentia | Cricetidae | Bank vole | *Clethrionomys glareolus* | 176 | 70 | S. Bondrup-Nielsen, R. A. Ims, Reversed sexual size dimorphism in microtines: Are females larger than males or are males smaller than females? *Evol. Ecol.* **4**, 261–272 (1990). |
| Rodentia | Cricetidae | Grey red-backed vole | *Clethrionomys rufocanus* | 159 | 109 | S. Bondrup-Nielsen, R. A. Ims, Reversed sexual size dimorphism in microtines: Are females larger than males or are males smaller than females? *Evol. Ecol.* **4**, 261–272 (1990). |
| Rodentia | Cricetidae | Northern red-backed vole | *Clethrionomys rutilus* | 39 | 20 | S. Bondrup-Nielsen, R. A. Ims, Reversed sexual size dimorphism in microtines: Are females larger than males or are males smaller than females? *Evol. Ecol.* **4**, 261–272 (1990). |

| | | | | | | |
|---|---|---|---|---|---|---|
| Rodentia | Cricetidae | Russet oryzomys | *Euryoryzomys russatus* | 209 | 157 | F. Gonçalves, R. S. Bovendorp, G. Beca, C. Bello, R. Costa-Pereira, R. L. Muylaert, R. R. Rodarte, N. Villar, R. Souza, M. E. Graipel, J. J. Cherem, D. Faria, J. Baumgarten, M. R. Alvarez, E. M. Vieira, N. Cáceres, R. Pardini, Y. L. R. Leite, L. P. Costa, M. A. R. Mello, E. Fischer, F. C. Passos, L. H. Varzinczak, J. A. Prevedello, A. P. Cruz-Neto, F. Carvalho, A. R. Percequillo, A. Paviolo, A. Nava, J. M. B. Duarte, N. U. de la Sancha, E. Bernard, R. G. Morato, J. F. Ribeiro, R. G. Becker, G. Paise, P. S. Tomasi, F. Vélez-Garcia, G. L. Melo, J. Sponchiado, F. Cerezer, M. A. S. Barros, A. Q. S. de Souza, C. C. dos Santos, G. A. F. Giné, P. Kerches-Rogeri, M. M. Weber, G. Ambar, L. V. Cabrera-Martinez, A. Eriksson, M. Silveira, C. F. Santos, L. Alves, E. Barbier, G. C. Rezende, G. S. T. Garbino, É. O. Rios, A. Silva, A. T. A. Nascimento, R. S. de Carvalho, A. Feijó, J. Arrabal, I. Agostini, D. Lamattina, S. Costa, E. Vanderhoeven, F. R. de Melo, P. de Oliveira Laroque, L. Jerusalinsky, M. M. Valença-Montenegro, A. B. Martins, G. Ludwig, R. B. de Azevedo, A. Anzóategui, M. X. da Silva, M. Figuerêdo Duarte Moraes, A. Vogliotti, A. Gatti, T. Püttker, C. S. Barros, T. K. Martins, A. Keuroghlian, D. P. Eaton, C. L. Neves, M. S. Nardi, C. Braga, P. R. Gonçalves, A. C. Srbek-Araujo, P. Mendes, J. A. de Oliveira, F. A. M. Soares, P. A. Rocha, P. Crawshaw, M. C. Ribeiro, M. Galetti, Atlantic Mammal Traits: a data set of morphological traits of mammals in the Atlantic Forest of South America. *Ecology*. **99**, 498 (2018). |
| Rodentia | Cricetidae | Elegant rice rat | *Euryoryzomys nitidus* | 30 | 18 | H. de G. Bergallo, Ecology of a small mammal community in an Atlantic Forest Area in Southeastern Brazil. *Stud. Neotrop. Fauna Environ.* **29**, 197–217 (1994). |

| | | | | | | |
|---|---|---|---|---|---|---|
| Rodentia | Cricetidae | Long-clawed mole mouse | *Geoxus valdivianus* | 15 | 8 | A. I. Schulte-Hostedde, "Sexual size dimorphism in rodents" in *Rodent Societies: An Ecological and Evolutionary Perspective*, J. O. Wolff, P. W. Sherman, Eds. (University of Chicago Press, Chicago, 2007), pp. 115–128. |
| Rodentia | Cricetidae | Handleyomys alfaroi | *Handleyomys alfaroi* | 27 | 11 | D. Ocampo, K. G. Borja-Acosta, J. Lozano-Flórez, S. Cifuentes-Acevedo, E. Arbeláez-Cortés, N. J. Bayly, Á. Caguazango, B. Coral-Jaramillo, D. Cueva, F. Forero, J. P. Gómez, C. Gómez, M. A. Loaiza-Muñoz, G. A. Londoño, S. Losada-Prado, S. Pérez-Peña, H. E. Ramírez-Chaves, M. E. Rodríguez-Posada, J. Sanabria-Mejía, M. Sánchez-Martínez, V. H. Serrano-Cardozo, M. S. Sierra-Buitrago, J. Soto-Patiño, O. Acevedo-Charry, Body mass data set for 1,317 bird and 270 mammal species from Colombia. *Ecology*. **102** (2021), doi:10.1002/ecy.3273. |

| | | | | | | |
|---|---|---|---|---|---|---|
| Rodentia | Cricetidae | Melanomys caliginosus | *Melanomys caliginosus* | 10<br>6 | 54 | D. Ocampo, K. G. Borja-Acosta, J. Lozano-Flórez, S. Cifuentes-Acevedo, E. Arbeláez-Cortés, N. J. Bayly, Á. Caguazango, B. Coral-Jaramillo, D. Cueva, F. Forero, J. P. Gómez, C. Gómez, M. A. Loaiza-Muñoz, G. A. Londoño, S. Losada-Prado, S. Pérez-Peña, H. E. Ramírez-Chaves, M. E. Rodríguez-Posada, J. Sanabria-Mejía, M. Sánchez-Martínez, V. H. Serrano-Cardozo, M. S. Sierra-Buitrago, J. Soto-Patiño, O. Acevedo-Charry, Body mass data set for 1,317 bird and 270 mammal species from Colombia. *Ecology*. **102** (2021), doi:10.1002/ecy.3273. |
| Rodentia | Cricetidae | Short-tailed field vole | *Microtus agrestis* | 43 | 29 | E. Pankakoski, K. M. Tähkä, Relation of adrenal weight to sex, maturity and season in five species of small mammals. *Ann. Zool. Fenn.* **19**, 225–232 (2017). |
| Rodentia | Cricetidae | California vole | *Microtus californicus* | 97 | 68 | S. Bondrup-Nielsen, R. A. Ims, Reversed sexual size dimorphism in microtines: Are females larger than males or are males smaller than females? *Evol. Ecol.* **4**, 261–272 (1990). |
| Rodentia | Cricetidae | Gerbe's vole | *Microtus gerbei* | 5 | 9 | S. M. Giannoni, C. E. Borghi, M. R. Laconi, Swimming ability of the European fossorial vole *Microtus (T. ) gerbei* under different laboratory conditions. *Mammalia*. **65**, 143–150 (2001). |
| Rodentia | Cricetidae | Prairie vole | *Microtus ochrogaster* | 15<br>7 | 81 | S. Bondrup-Nielsen, R. A. Ims, Reversed sexual size dimorphism in microtines: Are females larger than males or are males smaller than females? *Evol. Ecol.* **4**, 261–272 (1990). |
| Rodentia | Cricetidae | Root vole | *Microtus oeconomus* | 19 | 30 | S. Bondrup-Nielsen, R. A. Ims, Reversed sexual size dimorphism in microtines: Are females larger than males or are males smaller than females? *Evol. Ecol.* **4**, 261–272 (1990). |

| | | | | | | |
|---|---|---|---|---|---|---|
| Rodentia | Cricetidae | Meadow vole | *Microtus pennsylvanicus* | 40 | 43 | A. I. Schulte-Hostedde, "Sexual size dimorphism in rodents" in *Rodent Societies: An Ecological and Evolutionary Perspective*, J. O. Wolff, P. W. Sherman, Eds. (University of Chicago Press, Chicago, 2007), pp. 115–128. |
| Rodentia | Cricetidae | Water vole | *Microtus richardsoni* | 86 | 142 | A. I. Schulte-Hostedde, "Sexual size dimorphism in rodents" in *Rodent Societies: An Ecological and Evolutionary Perspective*, J. O. Wolff, P. W. Sherman, Eds. (University of Chicago Press, Chicago, 2007), pp. 115–128. |
| Rodentia | Cricetidae | Wood lemming | *Myopus schisticolor* | 42 | 45 | S. Bondrup-Nielsen, R. A. Ims, Reversed sexual size dimorphism in microtines: Are females larger than males or are males smaller than females? *Evol. Ecol.* **4**, 261–272 (1990). |
| Rodentia | Cricetidae | Neacomys spinosus | *Neacomys spinosus* | 2 | 3 | D. Ocampo, K. G. Borja-Acosta, J. Lozano-Flórez, S. Cifuentes-Acevedo, E. Arbeláez-Cortés, N. J. Bayly, Á. Caguazango, B. Coral-Jaramillo, D. Cueva, F. Forero, J. P. Gómez, C. Gómez, M. A. Loaiza-Muñoz, G. A. Londoño, S. Losada-Prado, S. Pérez-Peña, H. E. Ramírez-Chaves, M. E. Rodríguez-Posada, J. Sanabria-Mejía, M. Sánchez-Martínez, V. H. Serrano-Cardozo, M. S. Sierra-Buitrago, J. Soto-Patiño, O. Acevedo-Charry, Body mass data set for 1,317 bird and 270 mammal species from Colombia. *Ecology*. **102** (2021), doi:10.1002/ecy.3273. |

| | | | | | | |
|---|---|---|---|---|---|---|
| Rodentia | Cricetidae | Bushy-tailed woodrat | *Neotoma cinerea* | 22 | 40 | A. I. Schulte-Hostedde, "Sexual size dimorphism in rodents" in Rodent Societies: An Ecological and Evolutionary Perspective, J. O. Wolff, P. W. Sherman, Eds. (University of Chicago Press, Chicago, 2007), pp. 115–128. |
| Rodentia | Cricetidae | Nephelomys childi | *Nephelomys childi* | 14 | 9 | D. Ocampo, K. G. Borja-Acosta, J. Lozano-Flórez, S. Cifuentes-Acevedo, E. Arbeláez-Cortés, N. J. Bayly, Á. Caguazango, B. Coral-Jaramillo, D. Cueva, F. Forero, J. P. Gómez, C. Gómez, M. A. Loaiza-Muñoz, G. A. Londoño, S. Losada-Prado, S. Pérez-Peña, H. E. Ramírez-Chaves, M. E. Rodríguez-Posada, J. Sanabria-Mejía, M. Sánchez-Martínez, V. H. Serrano-Cardozo, M. S. Sierra-Buitrago, J. Soto-Patiño, O. Acevedo-Charry, Body mass data set for 1,317 bird and 270 mammal species from Colombia. *Ecology*. **102** (2021), doi:10.1002/ecy.3273. |
| Rodentia | Cricetidae | Nephelomys meridensis | *Nephelomys meridensis* | 7 | 10 | D. Ocampo, K. G. Borja-Acosta, J. Lozano-Flórez, S. Cifuentes-Acevedo, E. Arbeláez-Cortés, N. J. Bayly, Á. Caguazango, B. Coral-Jaramillo, D. Cueva, F. Forero, J. P. Gómez, C. Gómez, M. A. Loaiza-Muñoz, G. A. Londoño, S. Losada-Prado, S. Pérez-Peña, H. E. Ramírez-Chaves, M. E. Rodríguez-Posada, J. Sanabria-Mejía, M. Sánchez-Martínez, V. H. Serrano-Cardozo, M. S. Sierra-Buitrago, J. Soto-Patiño, O. Acevedo-Charry, Body mass data set for 1,317 bird and 270 mammal species from Colombia. *Ecology*. **102** (2021), doi:10.1002/ecy.3273. |

| | | | | | | |
|---|---|---|---|---|---|---|
| Rodentia | Cricetidae | Darwin's leaf-eared mouse | *Phyllotys darwini* | 1361 | 1253 | M. Lima, F. Bozinovic, F. M. Jaksic, Body mass dynamics and growth patterns of leaf-eared mice *Phyllotis darwini* in a semi-arid region of the Neotropics. *Acta Theriol. (Warsz).* **42**, 15–24 (1997). |
| Rodentia | Cricetidae | Reithrodontomys mexicanus | *Reithrodontomys mexicanus* | 8 | 10 | D. Ocampo, K. G. Borja-Acosta, J. Lozano-Flórez, S. Cifuentes-Acevedo, E. Arbeláez-Cortés, N. J. Bayly, Á. Caguazango, B. Coral-Jaramillo, D. Cueva, F. Forero, J. P. Gómez, C. Gómez, M. A. Loaiza-Muñoz, G. A. Londoño, S. Losada-Prado, S. Pérez-Peña, H. E. Ramírez-Chaves, M. E. Rodríguez-Posada, J. Sanabria-Mejía, M. Sánchez-Martínez, V. H. Serrano-Cardozo, M. S. Sierra-Buitrago, J. Soto-Patiño, O. Acevedo-Charry, Body mass data set for 1,317 bird and 270 mammal species from Colombia. *Ecology*. **102** (2021), doi:10.1002/ecy.3273. |
| Rodentia | Cricetidae | Western harvest mouse | *Reithrodontomys megalotis* | 41 | 11 | Sara Weinstein, pers. comm. |
| Rodentia | Cricetidae | Rhipidomys fulviventer | *Rhipidomys fulviventer* | 51 | 12 | D. Ocampo, K. G. Borja-Acosta, J. Lozano-Flórez, S. Cifuentes-Acevedo, E. Arbeláez-Cortés, N. J. Bayly, Á. Caguazango, B. Coral-Jaramillo, D. Cueva, F. Forero, J. P. Gómez, C. Gómez, M. A. Loaiza-Muñoz, G. A. Londoño, S. Losada-Prado, S. Pérez-Peña, H. E. Ramírez-Chaves, M. E. Rodríguez-Posada, J. Sanabria-Mejía, M. Sánchez-Martínez, V. H. Serrano-Cardozo, M. S. Sierra-Buitrago, J. Soto-Patiño, O. Acevedo-Charry, Body mass data set for 1,317 bird and 270 mammal species from Colombia. *Ecology*. **102** (2021), doi:10.1002/ecy.3273. |

| | | | | | | |
|---|---|---|---|---|---|---|
| Rodentia | Cricetidae | Thomasomys baeops | *Thomasomys baeops* | 2 | 2 | D. Ocampo, K. G. Borja-Acosta, J. Lozano-Flórez, S. Cifuentes-Acevedo, E. Arbeláez-Cortés, N. J. Bayly, Á. Caguazango, B. Coral-Jaramillo, D. Cueva, F. Forero, J. P. Gómez, C. Gómez, M. A. Loaiza-Muñoz, G. A. Londoño, S. Losada-Prado, S. Pérez-Peña, H. E. Ramírez-Chaves, M. E. Rodríguez-Posada, J. Sanabria-Mejía, M. Sánchez-Martínez, V. H. Serrano-Cardozo, M. S. Sierra-Buitrago, J. Soto-Patiño, O. Acevedo-Charry, Body mass data set for 1,317 bird and 270 mammal species from Colombia. *Ecology*. **102** (2021), doi:10.1002/ecy.3273. |
| Rodentia | Cricetidae | Thomasomys cinereiventer | *Thomasomys cinereiventer* | 8 | 10 | D. Ocampo, K. G. Borja-Acosta, J. Lozano-Flórez, S. Cifuentes-Acevedo, E. Arbeláez-Cortés, N. J. Bayly, Á. Caguazango, B. Coral-Jaramillo, D. Cueva, F. Forero, J. P. Gómez, C. Gómez, M. A. Loaiza-Muñoz, G. A. Londoño, S. Losada-Prado, S. Pérez-Peña, H. E. Ramírez-Chaves, M. E. Rodríguez-Posada, J. Sanabria-Mejía, M. Sánchez-Martínez, V. H. Serrano-Cardozo, M. S. Sierra-Buitrago, J. Soto-Patiño, O. Acevedo-Charry, Body mass data set for 1,317 bird and 270 mammal species from Colombia. *Ecology*. **102** (2021), doi:10.1002/ecy.3273. |
| Rodentia | Cricetidae | Thomasomys contradictus | *Thomasomys contradictus* | 26 | 20 | D. Ocampo, K. G. Borja-Acosta, J. Lozano-Flórez, S. Cifuentes-Acevedo, E. Arbeláez-Cortés, N. J. Bayly, Á. Caguazango, B. Coral-Jaramillo, D. Cueva, F. Forero, J. P. Gómez, C. Gómez, M. A. Loaiza-Muñoz, G. A. Londoño, S. Losada-Prado, S. Pérez-Peña, H. E. Ramírez-Chaves, M. E. Rodríguez-Posada, J. Sanabria-Mejía, M. Sánchez-Martínez, V. H. Serrano-Cardozo, M. S. Sierra-Buitrago, J. Soto-Patiño, O. Acevedo-Charry, Body mass data set for 1,317 bird and 270 mammal species from Colombia. *Ecology*. **102** (2021), doi:10.1002/ecy.3273. |

| | | | | | | |
|---|---|---|---|---|---|---|
| Rodentia | Cricetidae | Thomasomys nicefori | *Thomasomys nicefori* | 12 | 11 | D. Ocampo, K. G. Borja-Acosta, J. Lozano-Flórez, S. Cifuentes-Acevedo, E. Arbeláez-Cortés, N. J. Bayly, Á. Caguazango, B. Coral-Jaramillo, D. Cueva, F. Forero, J. P. Gómez, C. Gómez, M. A. Loaiza-Muñoz, G. A. Londoño, S. Losada-Prado, S. Pérez-Peña, H. E. Ramírez-Chaves, M. E. Rodríguez-Posada, J. Sanabria-Mejía, M. Sánchez-Martínez, V. H. Serrano-Cardozo, M. S. Sierra-Buitrago, J. Soto-Patiño, O. Acevedo-Charry, Body mass data set for 1,317 bird and 270 mammal species from Colombia. *Ecology*. **102** (2021), doi:10.1002/ecy.3273. |
| Rodentia | Cricetidae | Thomasomys niveipes | *Thomasomys niveipes* | 38 | 16 | D. Ocampo, K. G. Borja-Acosta, J. Lozano-Flórez, S. Cifuentes-Acevedo, E. Arbeláez-Cortés, N. J. Bayly, Á. Caguazango, B. Coral-Jaramillo, D. Cueva, F. Forero, J. P. Gómez, C. Gómez, M. A. Loaiza-Muñoz, G. A. Londoño, S. Losada-Prado, S. Pérez-Peña, H. E. Ramírez-Chaves, M. E. Rodríguez-Posada, J. Sanabria-Mejía, M. Sánchez-Martínez, V. H. Serrano-Cardozo, M. S. Sierra-Buitrago, J. Soto-Patiño, O. Acevedo-Charry, Body mass data set for 1,317 bird and 270 mammal species from Colombia. *Ecology*. **102** (2021), doi:10.1002/ecy.3273. |
| Rodentia | Cricetidae | Transandinomys bolivaris | *Transandinomys bolivaris* | 10 | 5 | D. Ocampo, K. G. Borja-Acosta, J. Lozano-Flórez, S. Cifuentes-Acevedo, E. Arbeláez-Cortés, N. J. Bayly, Á. Caguazango, B. Coral-Jaramillo, D. Cueva, F. Forero, J. P. Gómez, C. Gómez, M. A. Loaiza-Muñoz, G. A. Londoño, S. Losada-Prado, S. Pérez-Peña, H. E. Ramírez-Chaves, M. E. Rodríguez-Posada, J. Sanabria-Mejía, M. Sánchez-Martínez, V. H. Serrano-Cardozo, M. S. Sierra-Buitrago, J. Soto-Patiño, O. Acevedo-Charry, Body mass data set for 1,317 bird and 270 mammal species from Colombia. *Ecology*. **102** (2021), doi:10.1002/ecy.3273. |

| | | | | | | |
|---|---|---|---|---|---|---|
| Rodentia | Cricetidae | Transandinomys talamancae | *Transandinomys talamancae* | 5 | 2 | D. Ocampo, K. G. Borja-Acosta, J. Lozano-Flórez, S. Cifuentes-Acevedo, E. Arbeláez-Cortés, N. J. Bayly, Á. Caguazango, B. Coral-Jaramillo, D. Cueva, F. Forero, J. P. Gómez, C. Gómez, M. A. Loaiza-Muñoz, G. A. Londoño, S. Losada-Prado, S. Pérez-Peña, H. E. Ramírez-Chaves, M. E. Rodríguez-Posada, J. Sanabria-Mejía, M. Sánchez-Martínez, V. H. Serrano-Cardozo, M. S. Sierra-Buitrago, J. Soto-Patiño, O. Acevedo-Charry, Body mass data set for 1,317 bird and 270 mammal species from Colombia. *Ecology*. **102** (2021), doi:10.1002/ecy.3273. |
| Rodentia | Ctenomyidae | Sand dune tuco-tuco | *Ctenomys australis* | 13 | 27 | M. S. Mora, F. Becerra, A. I. Vassallo, An allometric analysis of sexual dimorphism in *Ctenomys australis*: integrating classic morphometry and functional performance in vivo. *Zoology*. **127**, 27–39 (2018). |
| Rodentia | Ctenomyidae | Flamarion's tuco tuco | *Ctenomys flamarioni* | 6 | 9 | B. B. Kubiak, R. Maestri, L. R. Borges, D. Galiano, T. R. de Freitas, Interspecific interactions may not influence home range size in subterranean rodents: A case study of two tuco-tuco species (Rodentia: Ctenomyidae). *J. Mammal.* **98**, 1753–1759 (2017). |
| Rodentia | Ctenomyidae | Tiny tuco-tuco | *Ctenomys minutus* | 27 | 35 | J. R. Marinho, T. R. O. De Freitas, Population structure of *Ctenomys minutus* (Rodentia, Ctenomyidae) on the coastal plain of Rio Grande do Sul, Brazil. *Acta Theriol. (Warsz).* **51**, 53–59 (2006). |
| Rodentia | Ctenomyidae | Highland tuco-tuco | *Ctenomys opimus* | 10 | 16 | S. L. O'Brien, M. N. Tammone, P. A. Cuello, E. A. Lacey, Facultative sociality in a subterranean rodent, the highland tuco-tuco (*Ctenomys opimus*). *Biol. J. Linn. Soc.* **129**, 918–930 (2020). |
| Rodentia | Ctenomyidae | Tuco-tuco | *Ctenomys talarum* | 95 | 110 | A. I. Schulte-Hostedde, "Sexual size dimorphism in rodents" in *Rodent Societies: An Ecological and Evolutionary Perspective*, J. O. Wolff, P. W. Sherman, Eds. (University of Chicago Press, Chicago, 2007), pp. 115–128. |

| | | | | | | |
|---|---|---|---|---|---|---|
| Rodentia | Dasyproctidae | Azara's agouti | *Dasyprocta azarae* | 3 | 7 | F. Gonçalves, R. S. Bovendorp, G. Beca, C. Bello, R. Costa-Pereira, R. L. Muylaert, R. R. Rodarte, N. Villar, R. Souza, M. E. Graipel, J. J. Cherem, D. Faria, J. Baumgarten, M. R. Alvarez, E. M. Vieira, N. Cáceres, R. Pardini, Y. L. R. Leite, L. P. Costa, M. A. R. Mello, E. Fischer, F. C. Passos, L. H. Varzinczak, J. A. Prevedello, A. P. Cruz-Neto, F. Carvalho, A. R. Percequillo, A. Paviolo, A. Nava, J. M. B. Duarte, N. U. de la Sancha, E. Bernard, R. G. Morato, J. F. Ribeiro, R. G. Becker, G. Paise, P. S. Tomasi, F. Vélez-Garcia, G. L. Melo, J. Sponchiado, F. Cerezer, M. A. S. Barros, A. Q. S. de Souza, C. C. dos Santos, G. A. F. Giné, P. Kerches-Rogeri, M. M. Weber, G. Ambar, L. V. Cabrera-Martinez, A. Eriksson, M. Silveira, C. F. Santos, L. Alves, E. Barbier, G. C. Rezende, G. S. T. Garbino, É. O. Rios, A. Silva, A. T. A. Nascimento, R. S. de Carvalho, A. Feijó, J. Arrabal, I. Agostini, D. Lamattina, S. Costa, E. Vanderhoeven, F. R. de Melo, P. de Oliveira Laroque, L. Jerusalinsky, M. M. Valença-Montenegro, A. B. Martins, G. Ludwig, R. B. de Azevedo, A. Anzóategui, M. X. da Silva, M. Figuerêdo Duarte Moraes, A. Vogliotti, A. Gatti, T. Püttker, C. S. Barros, T. K. Martins, A. Keuroghlian, D. P. Eaton, C. L. Neves, M. S. Nardi, C. Braga, P. R. Gonçalves, A. C. Srbek-Araujo, P. Mendes, J. A. de Oliveira, F. A. M. Soares, P. A. Rocha, P. Crawshaw, M. C. Ribeiro, M. Galetti, Atlantic Mammal Traits: a data set of morphological traits of mammals in the Atlantic Forest of South America. *Ecology*. **99**, 498 (2018). |

| | | | | | | |
|---|---|---|---|---|---|---|
| Rodentia | Dinomyidae | Dinomys branickii | *Dinomys branickii* | 2 | 2 | D. Ocampo, K. G. Borja-Acosta, J. Lozano-Flórez, S. Cifuentes-Acevedo, E. Arbeláez-Cortés, N. J. Bayly, Á. Caguazango, B. Coral-Jaramillo, D. Cueva, F. Forero, J. P. Gómez, C. Gómez, M. A. Loaiza-Muñoz, G. A. Londoño, S. Losada-Prado, S. Pérez-Peña, H. E. Ramírez-Chaves, M. E. Rodríguez-Posada, J. Sanabria-Mejía, M. Sánchez-Martínez, V. H. Serrano-Cardozo, M. S. Sierra-Buitrago, J. Soto-Patiño, O. Acevedo-Charry, Body mass data set for 1,317 bird and 270 mammal species from Colombia. *Ecology*. **102** (2021), doi:10.1002/ecy.3273. |
| Rodentia | Dipodidae | Lesser jerboa A | *Jaculus jaculus sp. A* | 11 | 15 | A. B. Faleh, J. F. Cosson, C. Tatard, A. B. Othmen, K. Said, L. Granjon, Are there two cryptic species of the lesser jerboa *Jaculus jaculus* (Rodentia: Dipodidae) in Tunisia? Evidence from molecular, morphometric, and cytogenetic data. *Biol. J. Linn. Soc.* **99**, 673–686 (2010). |
| Rodentia | Dipodidae | Lesser jerboa B | *Jaculus jaculus sp.B* | 3 | 6 | A. B. Faleh, J. F. Cosson, C. Tatard, A. B. Othmen, K. Said, L. Granjon, Are there two cryptic species of the lesser jerboa *Jaculus jaculus* (Rodentia: Dipodidae) in Tunisia? Evidence from molecular, morphometric, and cytogenetic data. *Biol. J. Linn. Soc.* **99**, 673–686 (2010). |
| Rodentia | Echimyidae | Hoplomys gymnurus | *Hoplomys gymnurus* | 6 | 2 | D. Ocampo, K. G. Borja-Acosta, J. Lozano-Flórez, S. Cifuentes-Acevedo, E. Arbeláez-Cortés, N. J. Bayly, Á. Caguazango, B. Coral-Jaramillo, D. Cueva, F. Forero, J. P. Gómez, C. Gómez, M. A. Loaiza-Muñoz, G. A. Londoño, S. Losada-Prado, S. Pérez-Peña, H. E. Ramírez-Chaves, M. E. Rodríguez-Posada, J. Sanabria-Mejía, M. Sánchez-Martínez, V. H. Serrano-Cardozo, M. S. Sierra-Buitrago, J. Soto-Patiño, O. Acevedo-Charry, Body mass data set for 1,317 bird and 270 mammal species from Colombia. *Ecology*. **102** (2021), doi:10.1002/ecy.3273. |

| | | | | | | |
|---|---|---|---|---|---|---|
| Rodentia | Echimyidae | Boyacá spiny rat | *Proechimys chrysaeolus* | 18 | 13 | D. Ocampo, K. G. Borja-Acosta, J. Lozano-Flórez, S. Cifuentes-Acevedo, E. Arbeláez-Cortés, N. J. Bayly, Á. Caguazango, B. Coral-Jaramillo, D. Cueva, F. Forero, J. P. Gómez, C. Gómez, M. A. Loaiza-Muñoz, G. A. Londoño, S. Losada-Prado, S. Pérez-Peña, H. E. Ramírez-Chaves, M. E. Rodríguez-Posada, J. Sanabria-Mejía, M. Sánchez-Martínez, V. H. Serrano-Cardozo, M. S. Sierra-Buitrago, J. Soto-Patiño, O. Acevedo-Charry, Body mass data set for 1,317 bird and 270 mammal species from Colombia. *Ecology*. **102** (2021), doi:10.1002/ecy.3273. |
| Rodentia | Echimyidae | O'Connell's spiny rat | *Proechimys oconnelli* | 2 | 6 | D. Ocampo, K. G. Borja-Acosta, J. Lozano-Flórez, S. Cifuentes-Acevedo, E. Arbeláez-Cortés, N. J. Bayly, Á. Caguazango, B. Coral-Jaramillo, D. Cueva, F. Forero, J. P. Gómez, C. Gómez, M. A. Loaiza-Muñoz, G. A. Londoño, S. Losada-Prado, S. Pérez-Peña, H. E. Ramírez-Chaves, M. E. Rodríguez-Posada, J. Sanabria-Mejía, M. Sánchez-Martínez, V. H. Serrano-Cardozo, M. S. Sierra-Buitrago, J. Soto-Patiño, O. Acevedo-Charry, Body mass data set for 1,317 bird and 270 mammal species from Colombia. *Ecology*. **102** (2021), doi:10.1002/ecy.3273. |

| | | | | | | |
|---|---|---|---|---|---|---|
| Rodentia | Erethizontidae | Bahia porcupine | *Coendou insidiosus* | 10 | 8 | F. Gonçalves, R. S. Bovendorp, G. Beca, C. Bello, R. Costa-Pereira, R. L. Muylaert, R. R. Rodarte, N. Villar, R. Souza, M. E. Graipel, J. J. Cherem, D. Faria, J. Baumgarten, M. R. Alvarez, E. M. Vieira, N. Cáceres, R. Pardini, Y. L. R. Leite, L. P. Costa, M. A. R. Mello, E. Fischer, F. C. Passos, L. H. Varzinczak, J. A. Prevedello, A. P. Cruz-Neto, F. Carvalho, A. R. Percequillo, A. Paviolo, A. Nava, J. M. B. Duarte, N. U. de la Sancha, E. Bernard, R. G. Morato, J. F. Ribeiro, R. G. Becker, G. Paise, P. S. Tomasi, F. Vélez-Garcia, G. L. Melo, J. Sponchiado, F. Cerezer, M. A. S. Barros, A. Q. S. de Souza, C. C. dos Santos, G. A. F. Giné, P. Kerches-Rogeri, M. M. Weber, G. Ambar, L. V. Cabrera-Martinez, A. Eriksson, M. Silveira, C. F. Santos, L. Alves, E. Barbier, G. C. Rezende, G. S. T. Garbino, É. O. Rios, A. Silva, A. T. A. Nascimento, R. S. de Carvalho, A. Feijó, J. Arrabal, I. Agostini, D. Lamattina, S. Costa, E. Vanderhoeven, F. R. de Melo, P. de Oliveira Laroque, L. Jerusalinsky, M. M. Valença-Montenegro, A. B. Martins, G. Ludwig, R. B. de Azevedo, A. Anzóategui, M. X. da Silva, M. Figuerêdo Duarte Moraes, A. Vogliotti, A. Gatti, T. Püttker, C. S. Barros, T. K. Martins, A. Keuroghlian, D. P. Eaton, C. L. Neves, M. S. Nardi, C. Braga, P. R. Gonçalves, A. C. Srbek-Araujo, P. Mendes, J. A. de Oliveira, F. A. M. Soares, P. A. Rocha, P. Crawshaw, M. C. Ribeiro, M. Galetti, Atlantic Mammal Traits: a data set of morphological traits of mammals in the Atlantic Forest of South America. *Ecology*. **99**, 498 (2018). |
| Rodentia | Geomyidae | Botta's pocket gopher | *Thomomys bottae* | 69 | 123 | J. C. Daly, J. L. Patton, Growth, reproduction, and sexual dimorphism in *Thomomys bottae* pocket gophers. *J. Mammal.* **67**, 256–265 (1986). |

| | | | | | | |
|---|---|---|---|---|---|---|
| Rodentia | Gliridae | European edible dormouse | *Glis glis* | 9 | 30 | A. Čanády, L. Mošanský, P. Krišovský, Sexual size monomorphism and body variation in the fat dormouse *Glis glis* in Slovakia. *Biologia (Bratisl).* **71**, 1061–1066 (2016). |
| Rodentia | Gliridae | Haze, dormouse | *Muscardinus avellanarius* | 17 | 13 | I. Pretzlaff, D. Rau, K. H. Dausmann, Energy expenditure increases during the active season in the small, free-living hibernator *Muscardinus avellanarius*. *Mamm. Biol.* **79**, 208–214 (2014). |
| Rodentia | Heteromyidae | Merriam's kangaroo rat | *Dipodomys merriami* | 162 | 154 | G. J. Kenagy, G. A. Bartholomew, Seasonal reproductive patterns in five coexisting California desert rodent species. *Ecol. Monogr.* **55**, 371–396 (1985). |
| Rodentia | Heteromyidae | Chisel-toothed kangaroo rat | *Dipodomys microps* | 202 | 165 | G. J. Kenagy, G. A. Bartholomew, Seasonal reproductive patterns in five coexisting California desert rodent species. *Ecol. Monogr.* **55**, 371–396 (1985). |
| Rodentia | Heteromyidae | Heteromys anomalus | *Heteromys anomalus* | 5 | 3 | D. Ocampo, K. G. Borja-Acosta, J. Lozano-Flórez, S. Cifuentes-Acevedo, E. Arbeláez-Cortés, N. J. Bayly, Á. Caguazango, B. Coral-Jaramillo, D. Cueva, F. Forero, J. P. Gómez, C. Gómez, M. A. Loaiza-Muñoz, G. A. Londoño, S. Losada-Prado, S. Pérez-Peña, H. E. Ramírez-Chaves, M. E. Rodríguez-Posada, J. Sanabria-Mejía, M. Sánchez-Martínez, V. H. Serrano-Cardozo, M. S. Sierra-Buitrago, J. Soto-Patiño, O. Acevedo-Charry, Body mass data set for 1,317 bird and 270 mammal species from Colombia. *Ecology*. **102** (2021), doi:10.1002/ecy.3273. |
| Rodentia | Heteromyidae | Perognathus formosus | *Perognathus formosus* | 97 | 118 | G. J. Kenagy, G. A. Bartholomew, Seasonal reproductive patterns in five coexisting California desert rodent species. *Ecol. Monogr.* **55**, 371–396 (1985). |

| | | | | | | |
|---|---|---|---|---|---|---|
| Rodentia | Heteromyidae | Little pocket mouse | *Perognathus longimembris* | 66 | 67 | G. J. Kenagy, G. A. Bartholomew, Seasonal reproductive patterns in five coexisting California desert rodent species. *Ecol. Monogr.* **55**, 371–396 (1985). |
| Rodentia | Muridae | Spiny mouse | *Acomys cahirinus* | 77 | 20 | E. Shargal, N. Kronfeld-Schor, T. Dayan, Population biology and spatial relationships of coexisting spiny mice (*Acomys*) in Israel. *J. Mammal.* **81**, 1046–1052 (2000). |
| Rodentia | Muridae | Arabian spiny mouse | *Acomys dimidiatus* | 73 | 63 | J. Sarli, H. Luterman, A. N. Alagaili, O. B. Mohammed, N. C. Bennett, Seasonal reproduction in the Arabian spiny mouse, *Acomys dimidiatus* (Rodentia: Muridae) from Saudi Arabia: The role of rainfall and temperature. *J. Arid Environ.* **124**, 352–359 (2016). |
| Rodentia | Muridae | Southern African spiny mouse | *Acomys spinosissimus* | 27 | 17 | W. Verheyen, J. Hulselmans, W. Wendelen, H. Leirs, M. Corti, Contribution to the systematics and zoogeography of the East-African *Acomys spinosissimus* Peters 1852 species complex and the description of two new species (Rodentia: Muridae). *Zootaxa*. **3059**, 1–35 (2011). |
| Rodentia | Muridae | Golden spiny mouse | *Acomys russatus* | 119 | 44 | E. Shargal, N. Kronfeld-Schor, T. Dayan, Population biology and spatial relationships of coexisting spiny mice (*Acomys*) in Israel. *J. Mammal.* **81**, 1046–1052 (2000). |
| Rodentia | Muridae | Tete veld rat | *Aethomys ineptus* | 25 | 27 | S. P. Muteka, C. T. Chimimba, N. C. Bennett, Reproductive seasonality in the Tete veld rat (*Aethomys ineptus*) (Rodentia: Muridae) from southern Africa. *J. Zool.* **270**, 314–322 (2006). |
| Rodentia | Muridae | Namaqua rock rat | *Aethomys namaquensis* | 47 | 53 | S. P. Muteka, C. T. Chimimba, N. C. Bennett, Reproductive seasonality in *Aethomys namaquensis* (Rodentia: Muridae) from southern Africa. *J. Mammal.* **87**, 67–74 (2006). |

| | | | | | | |
|---|---|---|---|---|---|---|
| Rodentia | Muridae | Striped field mouse | *Apodemus agrarius* | 57 | 63 | L. Balćiauskiene, L. Balćiauskas, Pelvis of the striped field mouse *Apodemus agrarius* (Pallas, 1771): Sexual dimorphism and relation to body weight. *North. West. J. Zool.* **12**, 50–57 (2016). |
| Rodentia | Muridae | Small Japanese field mouse | *Apodemus argenteus* | 17 | 9 | H. Suzuki, H. Kashiwagi, H. Kajimura, How does the 120-year cycle mast seeding of dwarf bamboo affect the rodent population? Ecol. Process. 11 (2022), doi:10.1186/s13717-022-00385-x. |
| Rodentia | Muridae | Yellow-necked mouse | *Apodemus flavicollis* | 17 | 13 | E. Wołk, J. Kozłowski, Changes of body weight and hematological parameters in a fluctuating population of *Apodemus flavicollis*. *Acta Theriol. (Warsz).* **34** (1989). |
| Rodentia | Muridae | Ural field mouse | *Apodemus microps* | 4 | 7 | V. Vohralík, Distribution, skull morphometrics and systematic status of an isolated population of *Apodemus microps* (Mammalia: Rodentia) in NW Bohemia, Czech Republic. *Acta Soc. Zool. Bohemicae*. **66**, 67–80 (2002). |
| Rodentia | Muridae | Taiwan field mouse | *Apodemus semotus* | 160 | 24 | L.-K. Lin, S. Shiraishi, Reproductive biology of the Formosan wood mouse, *Apodemus semotus*. *J. Fac. Agric. Kyushu Univ.* **36**, 183–200 (1992). |
| Rodentia | Muridae | Large Japanese field mouse | *Apodemus speciosus* | 45 | 30 | H. Ueda, S. Takatsuki, Sexual dimorphism of *Apodemus speciosus* in wild populations. *Mammal Study*. **30**, 65–68 (2005). |
| Rodentia | Muridae | Wood mouse | *Apodemus sylvaticus* | 10 | 10 | I. T. Rosário, M. L. Mathias, Annual weight variaiton and reproductive cycle of the wood mouse (*Apodemus sylvaticus*) in a Mediterranean environment. *Mammalia*. **68**, 133–140 (2004). |
| Rodentia | Muridae | Pygmy field mouse | *Apodemus uralensis* | 12 | 23 | L. Balčiauskas, L. Balčiauskienė, R. Juškaitis, Body size and craniometry of the herb field mouse from Lithuania in the context of species range. *Biol.* **73**, 351–359 (2018). |

| | | | | | | |
|---|---|---|---|---|---|---|
| Rodentia | Muridae | Mindanao mossy forest mouse | *Apomys hylocetes* | 31 | 13 | L. R. Heaney, B. R. Tabaranza, E. A. Rickart, D. S. Balete, N. R. Ingle, The mammals of Mt. Kitanglad Nature Park, Mindanao, Philippines. *Fieldiana Zool.* **112**, 1–63 (2006). |
| Rodentia | Muridae | Mindanao montane forest mouse | *Apomys insignis* | 15 | 13 | L. R. Heaney, B. R. Tabaranza, E. A. Rickart, D. S. Balete, N. R. Ingle, The mammals of Mt. Kitanglad Nature Park, Mindanao, Philippines. *Fieldiana Zool.* **112**, 1–63 (2006). |
| Rodentia | Muridae | Lesser bandicoot rat | *Bandicota benaglensis* | 52 | 25 | S. A. Rana, N. Rana, M. Z. Iqbal, F. Shahbaz, Seasonal demographic variation in bandicoot rats, *Bandicota bengalensis* (Gray) in irrigated crops. *Pakistan J. Agric. Sci.* **43**, 63–68 (2006). |
| Rodentia | Muridae | Greater bandicoot rat | *Bandicota indica* | 15 | 33 | N. Thitipramote, J. Swanjarat, W. G. Breed, Reproductive biology of the greater bandicoot rat *Bandicota indica* (Rodentia: Muridae) in the rice fields of southern Thailand. *Curr. Zool.* **55**, 48–55 (2009). |
| Rodentia | Muridae | Luzon hairy-tailed rat | *Batomys granti* | 5 | 5 | D. S. Balete, E. A. Rickart, L. R. Heaney, S. A. Jansa, A new species of *Batomys* (Muridae, Rodentia) from southern Luzon Island, Philippines. *Proc. Biol. Soc. Washingt.* **128**, 22–39 (2015). |
| Rodentia | Muridae | Mindanao hairy-tailed rat | *Batomys salomonseni* | 14 | 10 | L. R. Heaney, B. R. Tabaranza, E. A. Rickart, D. S. Balete, N. R. Ingle, The mammals of Mt. Kitanglad Nature Park, Mindanao, Philippines. *Fieldiana Zool.* **112**, 1–63 (2006). |
| Rodentia | Muridae | Batomys uragon | *Batomys uragon* | 5 | 5 | D. S. Balete, E. A. Rickart, L. R. Heaney, S. A. Jansa, A new species of *Batomys* (Muridae, Rodentia) from southern Luzon Island, Philippines. *Proc. Biol. Soc. Washingt.* **128**, 22–39 (2015). |

| | | | | | | |
|---|---|---|---|---|---|---|
| Rodentia | Muridae | Blazed Luzon Shrew rat | *Celaenomys silaceus* | 4 | 11 | E. A. Rickart, L. R. Heaney, S. M. Goodman, S. A. Jansa, Review of the Philippine genera *Chrotomys* and *Celaenomys* (Murinae) and description of a new species. *J. Mammal.* **86**, 415–428 (2005). |
| Rodentia | Muridae | Isarog striped shrew-rat | *Chrotomys gonzalesi* | 2 | 6 | E. A. Rickart, L. R. Heaney, S. M. Goodman, S. A. Jansa, Review of the Philippine genera *Chrotomys* and *Celaenomys* (Murinae) and description of a new species. *J. Mammal.* **86**, 415–428 (2005). |
| Rodentia | Muridae | Luzon striped rat | *Chrotomys whiteheadi* | 7 | 19 | E. A. Rickart, L. R. Heaney, S. M. Goodman, S. A. Jansa, Review of the Philippine genera *Chrotomys* and *Celaenomys* (Murinae) and description of a new species. *J. Mammal.* **86**, 415–428 (2005). |
| Rodentia | Muridae | Namibian gerbil | *Gerbillicus leucocaster* | 93 | 65 | S. P. Muteka, C. T. Chimimba, A. D. Bastos, N. C. Bennett, The reproductive pattern of the *Gerbillicus* cf. *leucocaster* (Rodentia: Muridae) from Namibia. *Can. J. Zool.* **97** (2019). |
| Rodentia | Muridae | Spiny Boki Mekot rat | *Halmaheramys bokimekot* | 3 | 3 | P. H. Fabre, A. H. Reeve, Y. S. Fitriana, K. P. Aplin, K. M. Helgen, A new species of *Halmaheramys* (Rodentia: Muridae) from Bisa and Obi Islands (North Maluku Province, Indonesia). *J. Mammal.* **99**, 187–208 (2018). |
| Rodentia | Muridae | African Crested Rat | *Lophiomys imhausi* | 12 | 10 | S. B. Weinstein, K. N. Malanga, B. Agwanda, J. E. Maldonado, M. D. Dearing, The secret social lives of African crested rats, *Lophiomys imhausi*. *J. Mammal.* **101**, 1680–1691 (2020). |
| Rodentia | Muridae | Rajah spiny rat | *Maxomys rajah* | 19 | 13 | M. Nakagawa, H. Miguchi, K. Sato, T. Nakashizuka, A preliminary study of two sympatric *Maxomys* rats in Sarawak, Malaysia: spacing patterns and population dynamics. *Raffles Bull. Zool.* **55**, 381–387 (2007). |

| | | | | | | |
|---|---|---|---|---|---|---|
| Rodentia | Muridae | Whitehead's spiny rat | *Maxomys whiteadi* | 3 | 12 | M. Nakagawa, H. Miguchi, K. Sato, T. Nakashizuka, A preliminary study of two sympatric *Maxomys* rats in Sarawak, Malaysia: spacing patterns and population dynamics. *Raffles Bull. Zool.* **55**, 381–387 (2007). |
| Rodentia | Muridae | Sundevall's jird | *Meriones crassus* | 37 | 30 | A. I. Schulte-Hostedde, "Sexual size dimorphism in rodents" in *Rodent Societies: An Ecological and Evolutionary Perspective*, J. O. Wolff, P. W. Sherman, Eds. (University of Chicago Press, Chicago, 2007), pp. 115–128. |
| Rodentia | Muridae | Mongolian gerbil | *Meriones unguiculatus* | 61 | 84 | G. Wang, Sexual size dimorphism of group-living Mongolian gerbils *Meriones unguicutalus* (Muridae: Gerbillinae). *Eur. Zool. J.* **84**, 536–540 (2017). |
| Rodentia | Muridae | Spinifex hopping mouse | *Notomyx alexis* | 22 | 18 | M. Bauer, W. G. Breed, Testis mass of the spinifex hopping mouse and its impact on fertility potential. *J. Zool.* **274**, 349–356 (2008). |
| Rodentia | Muridae | Bush Karoo rat | *Otomys unisulcatus* | 11 | 9 | L. Wolhuter, J. Thomson, C. Schradin, N. Pillay, Life history traits of free-living bush Karoo rats (*Otomys unisulcatus*) in the semi-arid Succulent Karoo. *Mammal Res.* **67**, 73–81 (2022). |
| Rodentia | Muridae | Delectable soft-furred mouse | *Praomys delectorum* | 33 | 19 | W. T. Stanley, S. M. Goodman, W. T. Stanley, S. M. Goodman, Small mammal inventories in the East and West Usambara Mountains, Tanzania: Rodentia. *Fieldiana Life Earth Sci.* **4**, 53–73 (2011). |
| Rodentia | Muridae | Forest Soft Furred Mouse | *Praomys rostratus* | 25 | 35 | E. Fichet-Calvet, L. Audenaert, P. Barrière, E. Verheyen, Diversity, dynamics and reproduction in a community of small mammals in Upper Guinea, with emphasis on pygmy mice ecology. *Afr. J. Ecol.* **48**, 600–614 (2010). |

| | | | | | | |
|---|---|---|---|---|---|---|
| Rodentia | Muridae | Polynesian rat | *Rattus exulans* | 11 | 6 | J. Theuerkauf, S. Rouys, C. Chatreau, Mortality of radio-tracked wild rats in relation to transmitter weight and resilience of transmitters in relation to their design. *J. R. Soc. New Zeal.* **37**, 85–90 (2007). |
| Rodentia | Muridae | Brown rat | *Rattus norvegicus* | 13 | 14 | E. Mori, A. Iacucci, R. Castiglia, L. Santini, Sexual-size dimorphism in two synanthropic rat species: Comparison and eco-evolutionary perspectives. *Mamm. Biol.* **83**, 78–80 (2017). |
| Rodentia | Muridae | Black rat | *Rattus rattus* | 20 | 20 | E. Mori, A. Iacucci, R. Castiglia, L. Santini, Sexual-size dimorphism in two synanthropic rat species: Comparison and eco-evolutionary perspectives. *Mamm. Biol.* **83**, 78–80 (2017). |
| Rodentia | Muridae | African striped mouse | *Rhabdomys pumilio* | 43 | 38 | R. Rimbach, J. Jäger, N. Pillay, C. Schradin, Food availability is the main driver of seasonal changes in resting metabolic rate in African striped mice (*Rhabdomys pumilio*). *Physiol. Biochem. Zool.* **91**, 826–833 (2018). |
| Rodentia | Muridae | Labo shrew-rat | *Rhynchomys labo* | 14 | 7 | E. A. Rickart, D. S. Balete, R. M. Timm, P. A. Alviola, J. A. Esselstyn, L. R. Heaney, Two new species of shrew-rats (*Rhynchomys*: Muridae: Rodentia) from Luzon Island, Philippines. *J. Mammal.* **100**, 1112–1129 (2019). |
| Rodentia | Muridae | Isarog shrew-rat | *Rhynchomys isarogensis* | 10 | 2 | E. A. Rickart, D. S. Balete, R. M. Timm, P. A. Alviola, J. A. Esselstyn, L. R. Heaney, Two new species of shrew-rats (*Rhynchomys*: Muridae: Rodentia) from Luzon Island, Philippines. *J. Mammal.* **100**, 1112–1129 (2019). |
| Rodentia | Muridae | Mingan shrew-rat | *Rhynchomys mingan* | 8 | 10 | E. A. Rickart, D. S. Balete, R. M. Timm, P. A. Alviola, J. A. Esselstyn, L. R. Heaney, Two new species of shrew-rats (*Rhynchomys*: Muridae: Rodentia) from Luzon Island, Philippines. *J. Mammal.* **100**, 1112–1129 (2019). |

| | | | | | | |
|---|---|---|---|---|---|---|
| Rodentia | Muridae | Mount Data shrew-rat | *Rhynchomys soricoides* | 9 | 4 | D. S. Balete, E. A. Rickart, R. G. B. Rosell-Ambal, S. Jansa, L. R. Heaney, Descriptions of two new species of *Rhynchomys* Thomas (Rodentia: Muridae: Murinae) from Luzon Island, Philippines. *J. Mammal.* **88**, 287–301 (2007). |
| Rodentia | Muridae | Indian Gerbil | *Tatera indica* | 13 | 6 | A. H. Shehab, S. A. Al-Ahmad, F. F. Samara, Morphology and distribution of the Indian gerbil, *Tatera indica* (Hardwicke, 1807), in Syria: (Rodentia: Gerbillinae). *Zool. Middle East*. **52**, 3–10 (2011). |
| Rodentia | Muridae | Black-tailed tree rat | *Thallomys nigricauda* | 26 | 44 | J. Meyer, thesis, Philipps-University Marburg (2004). |
| Rodentia | Nesomyidae | White-bellied nesomys | *Nesomys audeberti* | 8 | 7 | J. M. Ryan, G. K. Creighton, L. H. Emmons, Activity patterns of two species of *Nesomys* (Muridae: Nesomyinae) in a Madagascar rain forest. *J. Trop. Ecol.* **9**, 101–107 (1993). |
| Rodentia | Nesomyidae | Island mouse | *Nesomys rufus* | 19 | 11 | J. M. Ryan, G. K. Creighton, L. H. Emmons, Activity patterns of two species of *Nesomys* (Muridae: Nesomyinae) in a Madagascar rain forest. *J. Trop. Ecol.* **9**, 101–107 (1993). |
| Rodentia | Nesomyidae | South African pouched mouse | *Saccostomus campestris* | 8 | 8 | N. Mzilikazi, thesis, University of Natal (2000). |
| Rodentia | Octodontidae | Common degu | *Octodon degu* | 10 | 8 | L. A. Ebensperger, F. Bozinovic, Energetics and burrowing behaviour in the semifossorial degu *Octodon degus* (Rodentia: Octodontidae). *J. Zool.* **252**, 179–186 (2000). |
| Rodentia | Octodontidae | Coruro | *Spalacopus cyanus* | 13 | 22 | S. Begall, H. Burda, M. H. Gallardo, Reproduction, postnatal development, and growth of social coruros, *Spalacopus cyanus* (Rodentia: Octodontidae), from Chile. *J. Mammal.* **80**, 210–217 (1999). |

| | | | | | | |
|---|---|---|---|---|---|---|
| Rodentia | Sciuridae | White-tailed antelope squirrel | *Ammospermophilus leucurus* | 130 | 144 | G. J. Kenagy, G. A. Bartholomew, Seasonal reproductive patterns in five coexisting California desert rodent species. *Ecol. Monogr.* **55**, 371–396 (1985). |
| Rodentia | Sciuridae | Gunnison's prairie dog | *Cynomys gunnisoni* | 12 | 25 | L. S. Rayor, Effects of habitat quality on growth, age of first reproduction, and dispersal in Gunnison's prairie dogs (*Cynomys gunnisoni*). *Can. J. Zool.* **63**, 2835–2840 (1985). |

| | | | | | | |
|---|---|---|---|---|---|---|
| Rodentia | Sciuridae | Brazilian squirrel | *Guerlinguetus brasiliensis* | 10 | 2 | F. Gonçalves, R. S. Bovendorp, G. Beca, C. Bello, R. Costa-Pereira, R. L. Muylaert, R. R. Rodarte, N. Villar, R. Souza, M. E. Graipel, J. J. Cherem, D. Faria, J. Baumgarten, M. R. Alvarez, E. M. Vieira, N. Cáceres, R. Pardini, Y. L. R. Leite, L. P. Costa, M. A. R. Mello, E. Fischer, F. C. Passos, L. H. Varzinczak, J. A. Prevedello, A. P. Cruz-Neto, F. Carvalho, A. R. Percequillo, A. Paviolo, A. Nava, J. M. B. Duarte, N. U. de la Sancha, E. Bernard, R. G. Morato, J. F. Ribeiro, R. G. Becker, G. Paise, P. S. Tomasi, F. Vélez-Garcia, G. L. Melo, J. Sponchiado, F. Cerezer, M. A. S. Barros, A. Q. S. de Souza, C. C. dos Santos, G. A. F. Giné, P. Kerches-Rogeri, M. M. Weber, G. Ambar, L. V. Cabrera-Martinez, A. Eriksson, M. Silveira, C. F. Santos, L. Alves, E. Barbier, G. C. Rezende, G. S. T. Garbino, É. O. Rios, A. Silva, A. T. A. Nascimento, R. S. de Carvalho, A. Feijó, J. Arrabal, I. Agostini, D. Lamattina, S. Costa, E. Vanderhoeven, F. R. de Melo, P. de Oliveira Laroque, L. Jerusalinsky, M. M. Valença-Montenegro, A. B. Martins, G. Ludwig, R. B. de Azevedo, A. Anzóategui, M. X. da Silva, M. Figuerêdo Duarte Moraes, A. Vogliotti, A. Gatti, T. Püttker, C. S. Barros, T. K. Martins, A. Keuroghlian, D. P. Eaton, C. L. Neves, M. S. Nardi, C. Braga, P. R. Gonçalves, A. C. Srbek-Araujo, P. Mendes, J. A. de Oliveira, F. A. M. Soares, P. A. Rocha, P. Crawshaw, M. C. Ribeiro, M. Galetti, Atlantic Mammal Traits: a data set of morphological traits of mammals in the Atlantic Forest of South America. *Ecology*. **99**, 498 (2018). |
| Rodentia | Sciuridae | Mexican ground squirrel | *Ictidomys mexicanus* | 17 | 12 | L. E. Schwanz, Annual cycle of activity, reproduction, and body mass in Mexican ground squirrels (*Spermophilus mexicanus*). *J. Mammal.* **87**, 1086–1095 (2006). |

| | | | | | | |
|---|---|---|---|---|---|---|
| Rodentia | Sciuridae | Hoary marmot | *Marmota caligata* | 69 | 63 | B. Carrier, S. Hamel, M. Garel, S. D. Côté, Coping with seasonality: dynamics of adult body mass and survival in an alpine hibernator. *Oikos*, 1–15 (2022). |
| Rodentia | Sciuridae | Yellow-bellied marmot | *Marmota flaviventris* | 12 | 26 | V. A. Matrosova, D. T. Blumstein, I. A. Volodin, E. V. Volodina, The potential to encode sex, age, and individual identity in the alarm calls of three species of Marmotinae. *Naturwissenschaften*. **98**, 181–192 (2011). |
| Rodentia | Sciuridae | Notosciuru s granatensis | *Notosciurus granatensis* | 17 | 10 | D. Ocampo, K. G. Borja-Acosta, J. Lozano-Flórez, S. Cifuentes-Acevedo, E. Arbeláez-Cortés, N. J. Bayly, Á. Caguazango, B. Coral-Jaramillo, D. Cueva, F. Forero, J. P. Gómez, C. Gómez, M. A. Loaiza-Muñoz, G. A. Londoño, S. Losada-Prado, S. Pérez-Peña, H. E. Ramírez-Chaves, M. E. Rodríguez-Posada, J. Sanabria-Mejía, M. Sánchez-Martínez, V. H. Serrano-Cardozo, M. S. Sierra-Buitrago, J. Soto-Patiño, O. Acevedo-Charry, Body mass data set for 1,317 bird and 270 mammal species from Colombia. *Ecology*. **102** (2021), doi:10.1002/ecy.3273. |
| Rodentia | Sciuridae | Siberian flying squirrel | *Pteromys volans* | 33 | 28 | I. K. Hanski, P. C. Stevens, P. Ihalempia, V. Selonen, Home-range size, movements, and nest-site use in the Siberian flying squirrel, *Pteromys volans*. *J. Mammal.* **81**, 798–809 (2000). |
| Rodentia | Sciuridae | Eastern grey squirrel | *Sciurus carolinensis* | 16 | 23 | R. G. Tague, Pelvic sexual dimorphism among species monomorphic in body size: Relationship to relative newborn body mass. *J. Mammal.* **97**, 503–517 (2016). |
| Rodentia | Sciuridae | Red squirrel | *Sciurus vulgaris* | 51 | 39 | L. A. Wauters, G. Amori, G. Aloise, S. Gippoliti, P. Agnelli, A. Galimberti, M. Casiraghi, D. Preatoni, A. Martinoli, New endemic mammal species for Europe: *Sciurus meridionalis* (Rodentia, Sciuridae). *Hystrix, Ital. J. Mammal.* **28**, 1–8 (2017). |

| | | | | | | |
|---|---|---|---|---|---|---|
| Rodentia | Sciuridae | Calabrian black squirrel | *Sciurus meridionalis* | 6 | 5 | L. A. Wauters, G. Amori, G. Aloise, S. Gippoliti, P. Agnelli, A. Galimberti, M. Casiraghi, D. Preatoni, A. Martinoli, New endemic mammal species for Europe: *Sciurus meridionalis* (Rodentia, Sciuridae). *Hystrix, Ital. J. Mammal.* **28**, 1–8 (2017). |
| Rodentia | Sciuridae | Yellow ground squirrel | *Spermophilus fulvus* | 25 | 25 | V. A. Matrosova, D. T. Blumstein, I. A. Volodin, E. V. Volodina, The potential to encode sex, age, and individual identity in the alarm calls of three species of Marmotinae. *Naturwissenschaften*. **98**, 181–192 (2011). |
| Rodentia | Sciuridae | Golden-mantled ground squirrel | *Spermophillus saturatus* | 8 | 12 | G. J. Kenagy, S. M. Sharbaugh, K. A. Nagy, Annual cycle of energy and time expenditure in a golden-mantled ground squirrel population. *Oecologia*. **78**, 269–282 (1989). |
| Rodentia | Sciuridae | Speckled ground squirrel | *Spermophilus suslicus* | 26 | 26 | V. A. Matrosova, D. T. Blumstein, I. A. Volodin, E. V. Volodina, The potential to encode sex, age, and individual identity in the alarm calls of three species of Marmotinae. *Naturwissenschaften*. **98**, 181–192 (2011). |
| Rodentia | Sciuridae | Anatolian ground squirrel | *Spermophillus xanthoprymnus* | 78 | 90 | M. K. Gür, H. Gür, *Spermophilus xanthoprymnus* (Rodentia: Sciuridae). *Mamm. Species*. **42**, 183–194 (2010). |
| Rodentia | Sciuridae | Cliff chipmunk | *Tamias dorsalis* | 98 | 123 | A. L. Kilanowski, J. L. Koprowski, Female-biased sexual size dimorphism: Ontogeny, seasonality, and fecundity of the cliff chipmunk (*Tamias dorsalis*). *J. Mammal.* **98**, 204–210 (2017). |
| Rodentia | Sciuridae | North American red squirrel | *Tamiasciurus hudsonicus* | 33 | 32 | J. E. H. Patterson, P. Neuhaus, S. J. Kutz, K. E. Ruckstuhl, Patterns of ectoparasitism in North American red squirrels (*Tamiasciurus hudsonicus*): Sex-biases, seasonality, age, and effects on male body condition. *Int. J. Parasitol. Parasites Wildl.* **4**, 301–306 (2015). |

| | | | | | | |
|---|---|---|---|---|---|---|
| Rodentia | Sciuridae | Richardson's ground squirrel | *Urocitellus richardsonii* | 8 | 8 | R. J. Keeley, D. K. Burger, D. M. Saucier, A. N. Iwaniuk, The size of non-hippocampal brain regions varies by season and sex in Richardson's ground squirrel. *Neuroscience*. **289**, 194–206 (2015). |
| Rodentia | Sciuridae | Cape ground squirrel | *Xerus inauris* | 50 | 55 | J. M. Waterman, Reproductive biology of a tropical, non-hibernating ground squirrel. *J. Mammal.* **77**, 134–146 (1996). |
| Rodentia | Sminthidae | Northern birch mouse | *Sicista betulina* | 7 | 4 | I. A. Volodin, A. V. Klenova, O. G. Ilchenko, E. V. Volodina, High frequency audible calls in northern birch mice *Sicista betulina* in response to handling: Effects of individuality, sex and body mass on the acoustics. *BMC Res. Notes*. **12**, 1–7 (2019). |
| Rodentia | Spalacidae | Upper Galilee Mountains blind mole rat | *Spalax galili* | 5 | 5 | J. Šklíba, M. Lövy, S. C. W. Koeppen, L. Pleštilová, M. Vitámvás, E. Nevo, R. Šumbera, Activity of free-living subterranean blind mole rats *Spalax galili* (Rodentia: Spalacidae) in an area of supposed sympatric speciation. *Biol. J. Linn. Soc.* **118**, 280–291 (2016). |
| Rodentia | Zapodidae | Preble's meadow jumping mouse | *Zapus hudsonius preblei* | 96 | 76 | R. A. Schorr, P. M. Lukacs, G. L. Florant, Body mass and winter severity as predictors of overwinter survival in Preble's meadow jumping mouse. *J. Mammal.* **90**, 17–24 (2009). |
| Scandentia | Tupaiidae | Northern tree shrew | *Tupaia belangeri chinensis* | 454 | 538 | X. Wu, Q. Chang, Y. Zhang, X. Zou, L. Chen, L. Zhang, L. Lv, B. Liang, Relationships between body weight, fasting blood glucose concentration, sex and age in tree shrews (*Tupaia belangeri chinensis*). *J. Anim. Physiol. Anim. Nutr. (Berl).* **97**, 1179–1188 (2013). |

| | | | | | | |
|---|---|---|---|---|---|---|
| Scandentia | Tupaiidae | Common treeshrew | *Tupaia glis* | 3 | 2 | A. Mariana, M. N. Shukor, N. H. Muhd, N. B. Intan, T. M. Ho, Movements and home range of a common species of tree-shrew, *Tupaia glis*, surrounding houses of otoacariasis cases in Kuantan, Pahang, Malaysia. *Asian Pac. J. Trop. Med.* **3**, 427–434 (2010). |
| Scandentia | Tupaiidae | Large treeshrew | *Tupaia tana* | 12 | 10 | J. Munshi-South, thesis, University of Maryland (2006). |
